# A cross-biomes bacterial diversity shed light on ocean-atmosphere microbial transmission

**DOI:** 10.1101/2021.06.06.445733

**Authors:** Naama Lang-Yona, J. Michel Flores, Rotem Haviv, Adriana Alberti, Julie Poulain, Caroline Belser, Miri Trainic, Daniella Gat, Hans-Joachim Ruscheweyh, Patrick Wincker, Shinichi Sunagawa, Yinon Rudich, Ilan Koren, Assaf Vardi

## Abstract

Microbes are ubiquitous in the oceans and the atmosphere, playing essential roles in biogeochemical processes. The bio-exchanges between the two environments can provide important insights into microbial distribution and diversity but are still not well understood. We simultaneously surveyed the genomic diversity of airborne and marine bacterial communities across 15 000 kilometers in the Atlantic and Pacific oceans. Higher variability of microbial community composition was observed in the atmosphere than in the ocean surface waters. In addition, a greater similarity was observed between oceans than their overlaying atmosphere, and between atmospheric samples than with the ocean beneath. We additionally detected a higher coverage rate and relative abundance of marine bacteria in the Pacific atmosphere as compared to the Atlantic, while the dominant fraction in the Atlantic atmosphere was annotated as soil-associated bacteria. This study advances our understanding of microbial dispersion in the ocean, the atmosphere, and the exchange between them, as well as their potential impact on microbial composition, ecology, and biogeochemistry.

## Introduction

Microbes are ubiquitous in oceanic and atmospheric environments. Within the ocean, they account for approximately 70% of the total marine biomass (1), playing a crucial role in biogeochemical cycles (such as the carbon, nitrogen and sulfur cycles) (2). In the atmosphere, the microbial community comprises a major part of atmospheric bioaerosols (3), but little is known about the factors that affect their diversity, abundance (4), and role in this environment (5).

Understanding the ocean role as a source and sink of microorganisms and the transport of airborne bacteria can provide important insights into microbial spatial distribution, diversity, and the interplay between terrestrial communities and their transmission over oceanic regions. Over land, airborne bacteria are emitted from a wide range of sources, from anthropogenic to natural ecosystems. Over the oceans, bacteria can be locally emitted as sea spray aerosols, generated at the ocean’s surface by wind-driven processes, or transferred into the marine atmosphere through long-range transport from terrestrial sources (4). Due to their aerodynamic sizes, it has been hypothesized that all airborne bacteria can disperse globally and may proliferate in any habitat with suitable environmental conditions (6, 7). Thus, it can be expected that the airborne bacterial community aloft a given marine environment would exhibit a large fraction of the local marine microbiome.

Indeed, biodiversity studies indicated a link between microbial community composition and geographic locations (7, 8). Nevertheless, the geo-distribution of microorganisms in the interface between the ocean and the atmosphere is still underexplored, as the type, amount, and efficiency of particle emissions from the ocean, or atmosphere deposition hold large uncertainties. Exploring such distributions and microbial fluxes can improve our understanding of the metabolic capabilities introduced into the ocean and the enrichment of local marine diversity (9, 10). They may also affect biogeochemical processes upon deposition of bacteria into the ocean or their release to the atmosphere.

In this study, we present the spatially resolved composition of bacterial communities in the atmospheric marine boundary layer (AMBL) and the ocean surface along two basin-scale oceanic transects: The North Atlantic Ocean and the Western Pacific Ocean, covering approximately 15 000 kilometers. The similar sampling times, both placed in the north hemisphere (except for a few sampling points in the Pacific Ocean, southern to the equatorial), allow us to focus on non-seasonal changes in the microbial communities across these transects. We first explore microbial community compositions in the atmospheric and oceanic environments, revealing larger similarities between different oceans, in contrast to their overlaying atmospheres. By focusing on specific genera, we track terrestrial-associated bacteria in the airborne community and show distinct patterns of marine bacterial emission into the atmosphere. We thus suggest that bacterial geo-distribution theories and models should include constraints between the hydrosphere and atmosphere.

## Materials and Methods

### Sampling

Details of the expedition and sampling system used are described extensively in Flores *et al*. (11). In short, marine aerosols were collected aboard the *R/V* Tara during the first year of the Tara Pacific Expedition (12). Airborne particles were collected at ∼15 m above sea level (ASL) during the Atlantic crossing from Lorient, France, to Miami, USA. After Miami, the inlet was relocated to ∼27 m ASL. The inlet was constructed out of conductive tubing of 1.9 cm inner diameter and a funnel (allowing the collection of all diameters) and mounted on the rear backstay of Tara. A custom-made aerosol filter system consisting of four 47 mm filter holders and one vacuum pump (Diaphragm pump ME 16 NT, VACUUBRAND BmbH & Co KG, Wertheim, Germany) was installed at the end of the inlet and used to collect the marine particles.

The flow through the filter system was 80 lpm (20 lpm for each filter) during the Atlantic crossing and 120 lpm (30 lpm for each filter) for the Pacific crossing. Three of the four filter holders were loaded with PVDF filters (47 mm diameter, 0.45 μm pore size, PAL, Port Washington, New York, USA), and were used for the current study. The flow rates of each filter holder were monitored continuously and recorded at the beginning and the end of each sampling event. Aerosols were collected for periods between approximately 12 – 24 hours (see Table S1 for the exact times). The filters were folded into a 2 ml cryotube and immediately dropped into liquid nitrogen, and these conditions were maintained while on board. Blank filters were collected by placing filters on the filter holders, closing the system for a few seconds, reopening the holders, folding the filters into cryotubes, and dropping into liquid nitrogen. These conditions were maintained while on board. Thorough validation tests were conducted to verify no boat-originated contamination masked the airborne bacterial population composition (See SI).

The surface water sample collection is described in detail in Gorsky *et al*. (12). In short, a “Dolphin sampler”, collecting < 2000 μm size particles, and connected to a large volume peristaltic pump installed on the deck (max flow rate = 3 m^3^ h^−1^), was used for water sampling. Each sample endured for ∼ 120 min. Water serial filtration (<0.22, 0.22–3, and 3–20 µm) was performed using 142 mm diameter polycarbonate filters (Millipore, Burlington, Massachusetts, USA), and the 0.22-3 µm fraction, where free bacteria are expected to be concentrated, was used for the current study. The filters were folded into a 5 ml cryotube and immediately dropped into liquid nitrogen, and these conditions were maintained while on board. The samples were shipped to the laboratory on dry ice and kept at -80°C until DNA extraction was carried in the laboratory.

### Back-Trajectory Analysis

The air mass origins were tracked using the National Oceanic and Atmospheric Administration HYSPLIT trajectory model and Global Data Assimilation System meteorological data (13). Although the average residence time for 3 µm particles is of about 4.7 days (5), as considerable terrestrial influence was observed, the model was run to obtain the 240-hour (10 days) back trajectories using the “Ensemble option” at an endpoint height of 250 m, which is the minimum height for an optimal configuration of the ensemble. The presented back trajectories in Fig. 1 are the average of the 27 back trajectories produced by the Ensemble option.

**Figure 1.**
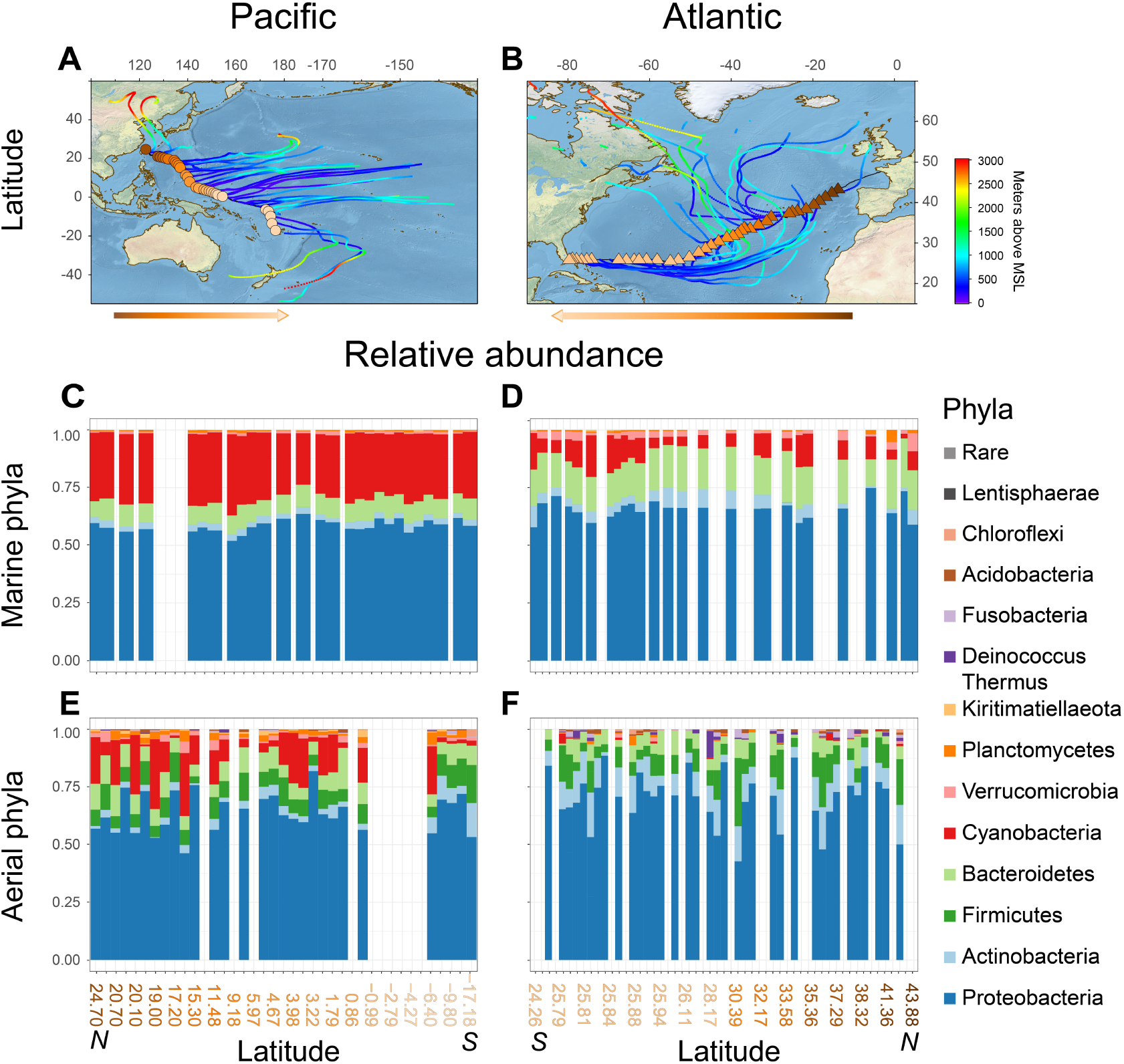
Regional distribution of airborne bacterial phyla above the Pacific and Atlantic oceans. Air-sampling locations and back trajectories, with height above mean sea levels (MSL) indicated with color scale for the Pacific (A) and Atlantic (B) atmospheric samples. Phyla relative abundance in the water (C and D) and the air (E and F) for taxa observed in > 5% of the samples in the Atlantic and Pacific transects, respectively.

### DNA extraction and sequencing

DNA extraction from 0.22-3 μm water filters was performed as described in Alberti *et al*. (14). Extraction of DNA from the air filters, with minute amounts of DNA, was carried out using the DNeasy PowerWater Kit (Qiagen, Hilden, Germany), following thorough optimization procedure for the extraction of low DNA concentrations from air filters (see SI text and Fig. S8). The processing of the water and air samples was performed separately, with random DNA extraction order for the different transects, thus preventing cross-contaminations between the different environments, and geographic location.

The DNA concentrations were evaluated with a Qubit® 3.0 Fluorometer (ThermoFischer, Massachusetts, USA), using the DeNovix (Wilmington, Delaware, USA) dsDNA Ultra High Sensitivity Assay. For DNA sequencing, the bacterial V4–V5 region of the 16S rRNA gene (515Fa: 5’–GTGYCAGCMGCCGCGGTAA–3’, and 926R: 5’–CCGYCAATTYMTTTRAGTTT–3’)(15), was amplified. A PCR mix of 25 μl was prepared in triplicate using 1X Mytaq mix (Bioline, London, UK), 0.2 μM primers, 4 μl DNA extract, and PCR-grade water (Merck, Darmstadt, Germany). A no-template control was included in all runs, as well as DNA from a mock community (ZymoBIOMICS Microbial Community DNA Standard; Zymo, Irvine, CA, USA). The PCR products were validated on 1% agarose gel, and triplicates were pooled and sent to Genoscope, the French National DNA Sequencing Facility. The PCR products of the water samples from the Pacific transect were sent to the DNA Sequencing Facility at the University of Illinois at Chicago. DNA sequencing in both facilities was conducted using Illumina MiSeq sequencing technology (maximum read length of 2×300 base pairs). No batch effects were observed, as validated by mock community positive control sequencing.

### Sequences Processing

Raw amplicon reads provided from the Genoscope sequencing facility were processed using the Tara Pacific metabarcoding (16S) pipeline up to the read margin stage using usearch v11(16), followed by the DADA2 pipeline (version 1.12) (17), using R (dada2 package). The merged reads (provided as .fastq files) were trimmed and filtered by removing reads exceeding the maximum expected error of 2 bp or an ambiguous read. The reads were dereplicated to acquire unique sequences, which were used to infer sequence variants with the trained error model. After chimeric sequence removal, amplicon sequence variants (ASVs) were used to assign taxa. ASVs were taxonomically assigned using dada2, with two steps: classification of sequences against the SILVA nr version 132 training dataset, followed by an exact matching between ASVs and the SILVA species assignment dataset (version 132) providing species-level assignment. Raw reads provided from the sequencing facility of the University of Illinois were also processed using the dada2 pipeline. Forward and reversed reads were merged after trimming and filtering steps. Dereplication and annotation processes were carried out as described above. A total of 11 577 bacterial ASVs were obtained after error correction, chimera, Archean, chloroplast, and mitochondrial DNA removal. Possible shifts in microbial composition due to differences in analyses were excluded by comparing mock community positive controls sequenced and analyzed using both methods. The microbial population sequencing analysis was performed only after a thorough decontamination procedure, to mitigate putative contamination in sequence libraries from the air samples with low microbial biomass (18), as described in the SI. The raw 16S amplicon sequences were deposited in the European Nucleotide Archive at EMBL-EBI (accession numbers: PRJEB39048, and PRJEB38899).

### Environmental ontology

Environmental descriptive terms were extracted from the closest matches (97% identity) using the SEQenv pipeline for Python (version 1.3.0) with default parameters (19) and ENVO terms. The input data included FASTA files of sequences after quality control check and removal of blank contaminants per each sample, to be compared to highly similar sequences from public repositories (such as GenBank, using the NCBI nucleotide data base). Detailed description of the pipeline flow is given in the SI file. In the current study, the terms were further clustered into five main groups: marine, terrestrial, fresh water, anthropogenic, and unclassified, as detailed in Table S7.

### Statistical analysis

Statistical significance was analyzed using R (3.5.2) and Origin 2019. Analysis of similarities (ANOSIM) was used to verify the significance of the nonmetric multidimensional scaling (NMDS) ordination for taxonomic grouping (using the Bray-Curtis dissimilarity score, vegan package), and differences in phyla composition between environments. Analysis of molecular variance (AMOVA) was used to verify significance of the Bray-Curtis dissimilarity differences of bacterial composition between biomes. Alpha diversity values were represented by the Shannon index and were tested for differences using analysis of variance (ANOVA) after normal distribution was verified.

## Results

### Regional distribution of airborne and surface water bacterial phyla in the Pacific and Atlantic oceans

The two open ocean sailing transects examined in this study were as follows: western Pacific, sampled in June 2016 (Table S1), starting from Keelung, Taiwan, towards Fiji (Fig. 1A), and the Atlantic crossing, sampled in May 2017 (Table S1) from Lorient, France, to Miami, USA. (Fig. 1B). In the water, we found a relatively homogeneous phyla distribution within each transect, with Proteobacteria dominating both oceans (58 ± 3% in the Pacific and 66 ± 4% in the Atlantic; Fig. 1C and D, respectively). Cyanobacteria (29 ± 2% and 9 ± 4%) and Bacteroidetes (9 ± 1% and 17 ± 2%) were the next two most abundant phyla. When comparing to other marine microbiome studies, the phyla distribution of the near-surface water environment is similar (2, 20). For example, the Cyanobacteria to Proteobacteria ratios in our study are 0.49 ± 0.06 and 0.14 ± 0.07 in the Pacific and Atlantic surface water, respectively. Similarly-calculated ratios characterized in these oceanic regions are 0.43 and 0.16, for the Pacific (20) and Atlantic (2) regions, respectively.

In the AMBL, Proteobacteria was also the most dominant airborne bacterial phylum in both the Pacific and Atlantic oceans, with 69 ± 12% and 64 ± 8% average percentile abundance, respectively (Fig. 1E and F). However, we found a more heterogeneous distribution of bacterial phyla than that in ocean surface water samples (Fig. S1), even when considering an air mass presence of at least 120 hours over an oceanic path prior to sampling (Fig. 1A and 1B, colored lines). Other abundant phyla in the Pacific AMBL included Cyanobacteria (11 ± 9%), Bacteroidetes (8 ± 3%), Firmicutes (8 ± 3%), Actinobacteria (4 ± 3%), and Planctomycetes (2 ± 1%). In the Atlantic AMBL, the abundant phyla included Actinobacteria (11 ± 5%), Firmicutes (10 ± 6%), and Bacteroidetes (6 ± 3%; Fig. 1F), while Cyanobacteria was observed with an average abundance less than 0.5%.

Firmicutes, Actinobacteria, Fusobacteria, Deinococcus-Thermos and Acidobacteria were higher in abundance in the atmospheric samples of both oceans compared to the waters. Firmicutes were predominantly observed in the airborne samples (8 ± 3% and 10 ± 6% in the Pacific and Atlantic AMBL, respectively), with low (<1% in average) to non-significant abundance in the water samples. Firmicutes were previously detected in the oceanic environment, specifically the Bacillus genus (21), and Sul *et al*. found in low relative abundance (< 6% on average) across latitudes with little latitudinal dependence (22). Actinobacteria abundance was also significantly higher in the Atlantic AMBL compared to the water (*t*-test, *p*-value < 0.0001). Airborne Actinobacteria and Firmicutes have been previously connected to desert dust samples in the Eastern Mediterranean (23-25), and detected in different studies of airborne bacteria (26-28). In addition, Firmicutes are usually more abundant in soils than in marine surface water (2, 21). Therefore, these groups may represent a terrestrial source.

Members of the airborne-detected Fusobacteria, Deinococcus-Thermos and Acidobacteria phyla exhibit high physiological diversity in cell shapes and sizes (21, 29). Their presence in remote locations, such as Antarctica (30), airborne dust (23), and precipitation over the alpine (31), together with their detection above the western Pacific Ocean, after 120 h transport above the ocean, suggests that they are ubiquitous in the atmospheric environment.

The local primary production imprint in the AMBL was estimated by calculating the ratio of autotrophic to heterotrophic bacteria in the atmospheric and oceanic samples (Fig. S2A and b). The ratios in the Pacific atmosphere were more than an order of magnitude higher than those measured in the Atlantic atmosphere (mean values: 0.113 ± 0.109 and 0.006 ± 0.009, respectively, two sample *t*-test *p*-value <0.001; Fig. S2A and B). Additionally, the average ratio in the Pacific water is approximately four times higher than that in the Atlantic (0.392 ± 0.083, compared to 0.103 ± 0.052, respectively; two sample *t*-test *p*-value < 0.001). Similarly, a significantly higher relative abundance of cyanobacterial 16S rRNA gene was observed in the Pacific compared to the Atlantic atmosphere (Fig. S2C and D, Table S3) based on qPCR analysis. The observed difference in oceanic cyanobacterial abundance is consistent with results reported by Flombaum *et al*., showing a higher abundance of marine Synechococcales in the Pacific compared to the Atlantic (33).

In general, the atmospheric phyla composition showed high variations between samples, whereas in the marine microbiome, a more homogeneous and stable community structure was observed. This may relate to the lower biomass sampled in the air, thus more prone to changes in community composition. Furthermore, while the wind-driven surface water currents show connectivity between oceans with time scales of years, the atmospheric circulation time scales are in the range of days to weeks (34, 35).

### Similarities and differences in the atmospheric and oceanic microbiomes

The marine and atmospheric microbiomes were further analyzed at the ASV level. We found the variability in the airborne bacterial diversity and composition to be significantly higher compared to the water samples (Fig. 2A and B; Two-sample *t*-test, *p*-value < 0.001, and ANOSIM, *p*-value = 0.043, respectively), with a distinct difference between the air and water ASV composition (Fig. 2C; AMOVA, *p*-value < 0.01 between the atmosphere and oceanic samples of both Atlantic and Pacific environments).

**Figure 2.**
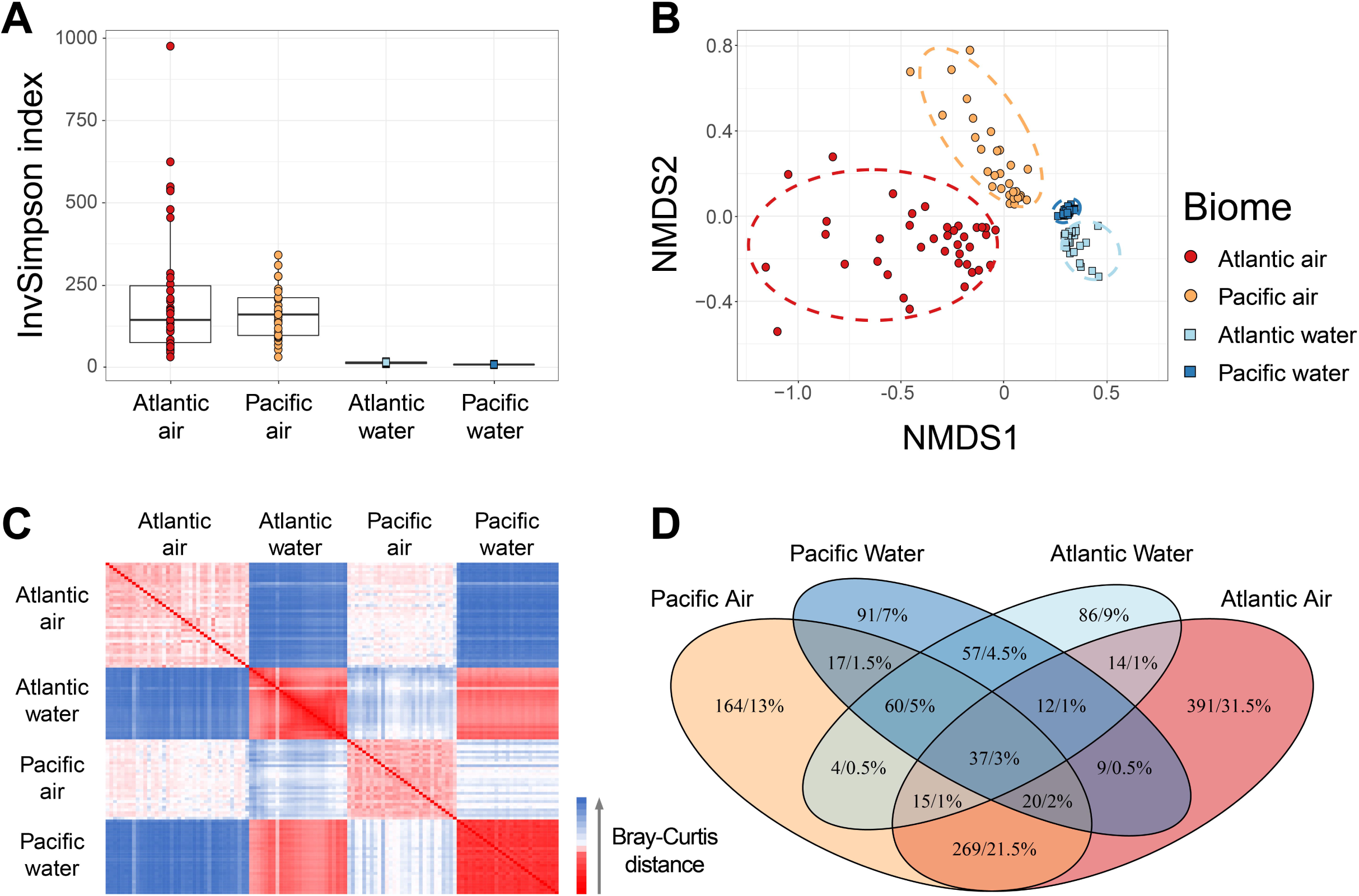
Similarities and differences in bacterial microbiomes of different oceanic environments. Biome-based diversity (based on the Inverse Simpson index) of amplicon sequence variants (ASVs) detected in more than 5% of the samples (A) and clusters represented by nonmetric multidimensional scaling (NMDS) ordination, based on Bray-Curtis dissimilarity metrics (B), with 95% confidence ellipses. The distance in ASV composition is presented in a heatmap of the Bray-Curtis dissimilarity between all analyzed samples (Bray-Curtis index varies between 0, for an identical ASV composition, and 1, for most distant ASVs on the samples) (C). A Venn diagram (*VennDiagram* 1.6.20) summarizes the number and the percentage out of the total number of bacterial taxa that were observed in the different biomes (D).

The surface water environments shared 166 taxa (comprising 28% of all ocean taxa; Fig. 2D), and the AMBL biomes shared 341 taxa (comprising 25% of all airborne taxa). Only 78 taxa in the Atlantic (7% of all Atlantic taxa) and 134 in the Pacific (15% of all Pacific taxa) were shared between an ocean and its corresponding atmosphere. We found different oceans to have a greater resemblance to one another than to their overlaying AMBL, and atmospheric samples from distinct locations (at least 13 000 km apart from each other) share more common taxa than with the ocean beneath. This suggests that the proximity of the sampled biomes is second in significance to the type of sampled environment.

A phylogenetic tree based on the bacterial 16S amplicon sequences provides an overview of the bacterial community and genetic distances between them in the observed marine environment (Fig. S3). Among the shared groups in the Atlantic and Pacific atmospheric biomes, the main phyla occurrences included 44% Proteobacteria, 19% Actinobacteria, 19% Firmicutes, and 10% Bacteroidetes. The high diversity in the common airborne bacterial taxa in both the Atlantic and Pacific is most likely a consequence of a short turnover time of the air mass, leading to continuously changing and dynamic community composition in the AMBL.

### Spatial distributions of bacteria across the Pacific and Atlantic environments

The spatial distribution of specific bacteria in the AMBL allows the identification of transported bacteria and indicates a nonrandom contribution from the marine environment. The bacterial ASVs were clustered into taxonomic groups, and >5% taxon occurrences were ranked based on the averaged abundance of the different phyla (Fig. 3 and listed in Tables S4-S6). The most abundant Proteobacteria detected only in the AMBL was *Paracoccus* (Table S4). *Paracoccus* strains have been isolated from different environments including soil (36), marine sediments (37), sewage (38), and have been detected in other atmospheric bacterial studies (39, 40). Some Proteobacteria, abundant in the air samples, were continuously detected in the surface water (*e*.*g*., *Halomonas, Pseudomonas, Idiomarina*, and *Methylobacterium*; Table S5), and thus assumed to be emitted from the local marine environment.

**Figure 3.**
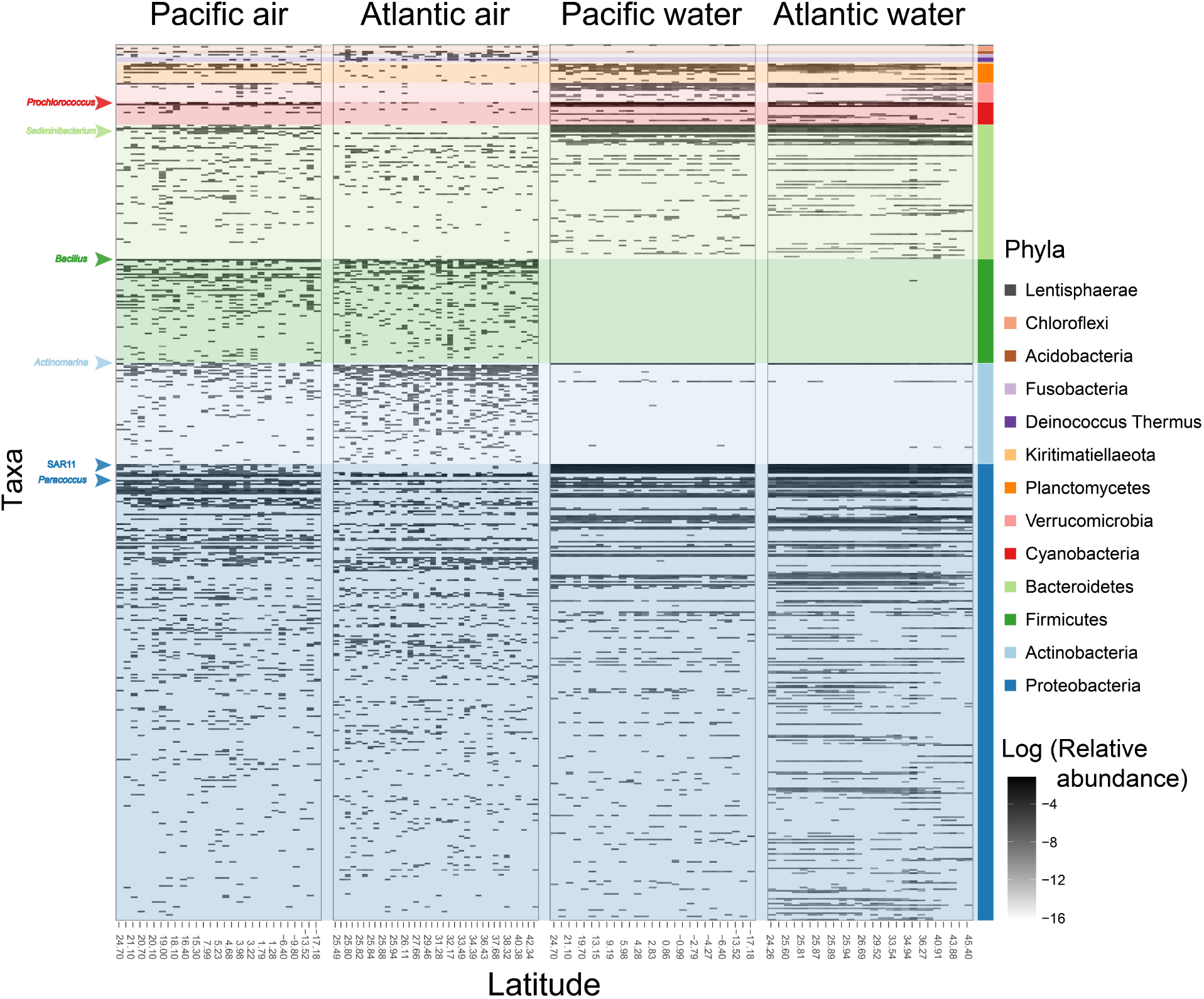
Spatial distribution of specific airborne microorganisms across the Atlantic and Pacific transects. The 16S amplicon sequence variants (ASVs) of Pacific air, Atlantic Air, Pacific water, and Atlantic water samples are categorized based on abundance for each phylum. ASVs were aggregated according to sequences and taxa annotation, and ASVs covering > 5% of all four environments are presented. Phyla are distinguished based on color code.

The most abundant genus in the phylum Actinobacteria was *Actinomarina*, which appeared in both oceans and in the Pacific air samples (Table S5). The main Firmicutes included Bacillus, known for their endospores that can remain dormant for years. *Bacillus* is a common bacterium found in transported desert dust (23), and the deep marine environment (41). Some *Bacillus* species are known for their unique metabolites and antagonistic activity against pathogens(42). We further explored the differences between the surface waters and the atmospheric Firmicutes population, which was dramatically under-represented in the water samples, with a focused phylogenetic tree targeting only Firmicutes ASVs (Fig. S4). Firmicutes ASVs were rarely detected in the water samples and differed phylogenetically from the atmospheric ASVs. The rare abundance of marine *bacilli* in the water may result from the preferential growth environment of the deep sea, coral and sediments (41), and their copiotrophic property (*i*.*e*., flourishing in environments with high nutrient availability) (43). Notably, *Tumebacillus*, and *Clostridium saccharobutylicum*, spore forming Firmicutes, were detected in the Atlantic AMBL. Endospores can survive harsh and dry conditions and thus might be transported through the air at higher survival rates than others. We also found ASVs assigned to genera that are known to include human-associated microbes (i.e., *Micrococcus, Actinomyces*; Table S4). Although not detected in the blank filters, we cannot exclude the possibility that those taxa may originate from the human activity onboard Tara. Nevertheless, the abundance of these genera is low, ranging on average between ∼0.1 – ∼2.7 % of the ASVs per filter. The most abundant genus in the phylum Bacteroidetes found in the atmospheric samples was *Sediminibacterium* (Table S5). This genus was previously found to contribute to the coral microbiome (44) and detected in air samples over the Great Barrier Reef (26) and the Mediterranean Sea (45). In this study, it was detected in the water samples of the Pacific Ocean solely, with low relative abundance (< 0.01%) and spatial coverage.

We continuously detected water-originated species in the air samples, with higher relative abundance in the Pacific AMBL. One such genus is the Cyanobacteria *Prochlorococcus*, considered a key and most abundant autotroph (46), found mainly in oligotrophic oceans (47). A notable difference in the relative abundance and appearance of marine bacteria in the Pacific compared to the Atlantic AMBL is also seen for different phyla, including Proteobacteria, Bacteroidetes, Verrucomicrobia, Planctomycetes, etc. (Fig. 3, and Table S5). While the increased fraction of dust-borne bacteria could partially explain the reduction in relative abundance of the local marine bacteria, marine-associated taxa were absent from a significantly high fraction of the sampled Atlantic atmosphere, suggesting other factors may also play a role in the observed difference between the two AMBLs. A clear case of such difference is seen for the Pelagibacterales (SAR-11 clade), representing approximately one-third of the oceanic surface water microbial community (48), and highly abundant in both oceans’ samples. Their appearance in the atmospheric samples is significantly lower (Two-sample *t*-test, *p*-value < 0.0001 in both tests; Fig. S4B). However, while in the Atlantic AMBL, the spatial coverage is minimal, the Pacific AMBL show almost a full spatial coverage of SAR-11 ASVs in these samples, with significantly higher relative abundance than the Atlantic (two-sample *t*-test, *p*-value < 0.001; Tables S5). A reduced aerosolized fraction of SAR-11 compared to the seawater was also observed in the Arctic Sea by Fahlgren *et al*. (49). The difference in abundance between the Atlantic and Pacific AMBLs could be related to properties of the sea-surface microlayer (SML), including thickness, concentration, and chemical composition, which was shown to differ according to changes in heat exchanges, microbial composition, oceanic waves, pollution and dust storms (50, 51). However, to fundamentally characterize the causing factors for the reduced detection of marine bacteria in the Atlantic AMBL, further investigation is required.

### Relationships between bacterial taxa in the marine boundary layer and other ecosystems

To better understand the environmental affiliation of the sampled microbiome we performed an environmental ontology (ENVO) analysis on all bacterial ASVs (52). The ENVO analysis allows a comparison of our dataset to published microbiomes found in other environments, by grouping all environments where the ASVs were previously detected (Fig. 4 and Table S7 detailing the annotations grouped into the five environments presented).

**Figure 4.**
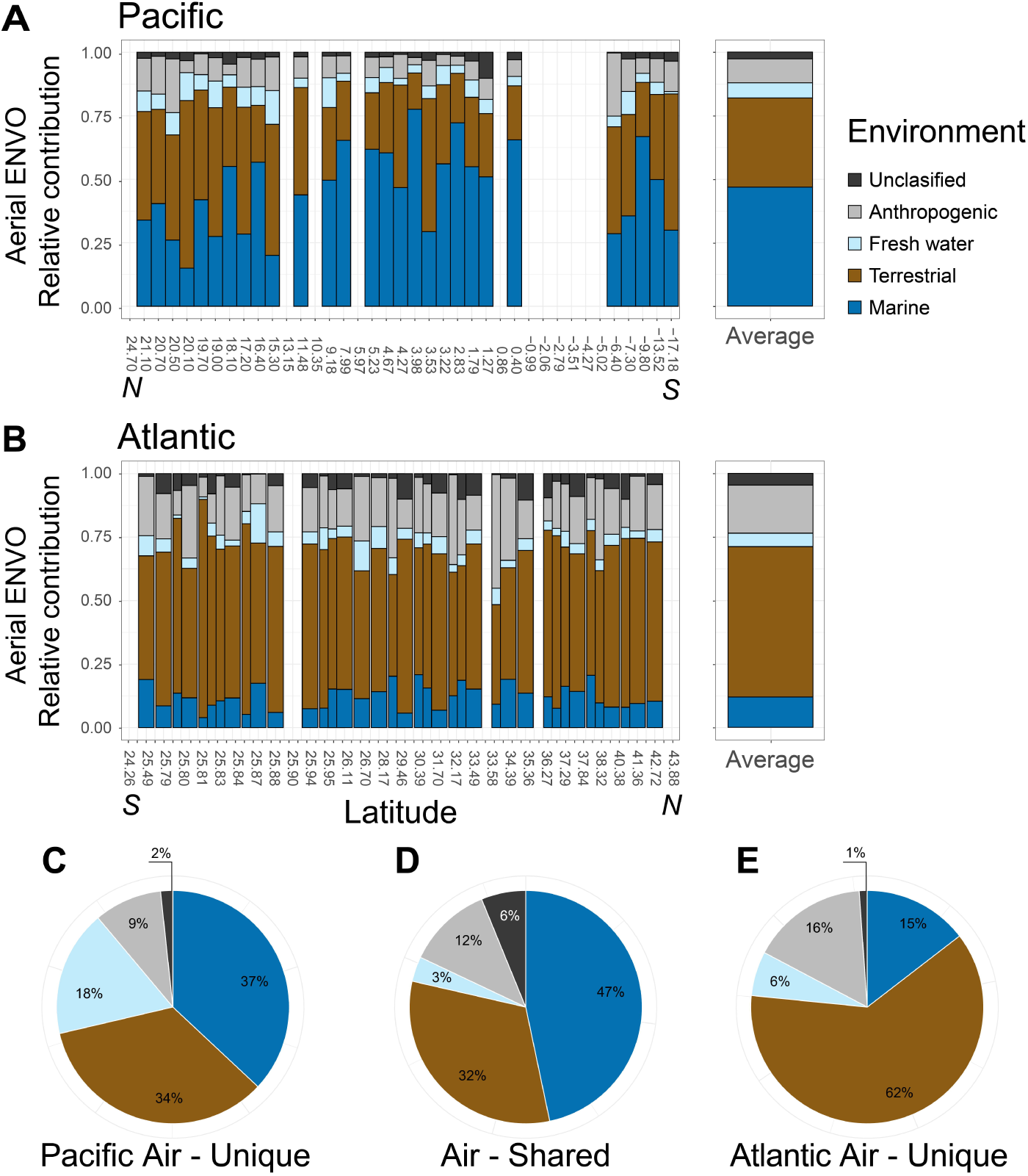
Association of airborne bacterial communities with other environments. The relative environmental ontology (ENVO) annotation of all amplicon sequence variants (ASVs) identified in the air samples clustered into five main groups (detailed terms are listed in Table S6) for the Pacific (A) and Atlantic (B) transects, as well as for Pacific air-unique ASVs (C), Pacific and Atlantic air-shared ASVs (D), and Atlantic air-unique ASVs (E).

The water samples showed 7% nonoceanic-annotated bacterial ASVs in the Pacific compared to 16% in the Atlantic (Fig. S5). Of these, 0.5%, in both environments, were from anthropogenic-annotated bacteria. In the Atlantic water samples, 9% were annotated as terrestrial bacteria, and might originate from sedimentation of dust particles or high maritime transportation activity in this environment (53).

The microbiome in the Pacific AMBL exhibited a higher fraction of marine-annotated bacteria, 47 ± 20% compared to 12 ± 5% in the Atlantic (Fig. 4A), while the Atlantic aerobiome was dominantly annotated as terrestrial (59 ± 16% compared to 35 ± 18% in the Pacific; Fig. 4B). A previous study by Mayol *et al*. (54) determined that overall, 25% of the airborne bacteria over the ocean originate from the marine environment, and 42% originate from terrestrial sources based on parameterizations of sea spray and deposition flux calculations. Higher soil-borne ENVO annotation was also observed in Firmicutes-targeted analysis (Fig. S6). Thus, airborne Firmicutes may originate from terrestrial long-range transport, but the extent in which their sedimentation and proliferation in the ocean is yet to be determined. Anthropogenic-annotated ASVs were higher in the Atlantic AMBL, with 19 ± 6%, compared to 9 ± 6% in the Pacific).

Both environments presented freshwater annotations (6 ± 3% and 5 ± 3% in the Pacific and Atlantic AMBL, respectively). The contribution of bacteria to the formation of water precipitation is of high interest, and studies revealed bacterial proteins from *e*.*g*., *Pseudomonas* and *Pantoea* sp., identified in the air samples (Table S4), can promote droplet freezing (55) and even detected bacterial activity in clouds (28). The freshwater annotation of the AMBL bacteria may indicate the presence of such bacteria in this environment.

Flores *et al*. (11, 56) have found higher concentrations of larger particles related to the deposition of mineral dust in aerosol sampled in the Atlantic compared to the Pacific transect. Other studies report massive dust quantities crossing over the Atlantic Ocean (53), which may introduce bacteria into this environment. The ENVO annotation retrieved from genomic databases corroborate these findings and emphasize the vast contribution of terrestrial dust-borne bacteria into the Atlantic Ocean. Additionally, we detected significantly higher DNA biomass in the Atlantic air filters compared to the Pacific (average of 639.6 ± 468.2 and 128.4 ± 54.4 pg m^-3^, respectively, *t*-test, *p*-value < 0.0001), implying a higher concentration of microbial cells per air volume in this region. It has been reported that the imprint of dust transport and human activity on the Pacific Ocean is relatively small (57), and that this area is considered a pristine and remote environment, while the Atlantic Ocean experiences high loads of dust, and anthropogenic impact, with a fast pace of change (58). Although with lower rates, the terrestrial annotations observed in the Pacific environment emphasize the long-range transport and dispersion of bacteria in this remote environment, implying that it cannot be considered a pristine environment, as observed in the Southern Ocean of Antarctica (59).

Environmental ontology analysis was further conducted for the shared and unique ASVs of the Atlantic and Pacific AMBLs (Fig. 4C-E). The unique ASVs detected in the Pacific samples were mainly annotated as marine (37%) and terrestrial (34%) (1 703 ASVs in total; Fig. 4C). The airborne Pacific-Atlantic shared bacterial ASVs were also composed mostly of marine (47%) and terrestrial (32%) annotated bacteria (220 ASVs in total; Fig. 4D). A large percentage of unique Atlantic airborne bacteria (62%) were annotated as terrestrial, whereas anthropogenic-(16%) and marine-(15%) annotated bacteria were less abundant (1 511 ASVs in total; Fig. 4E). The ubiquity of shared species found only in the atmospheric samples of the Atlantic and Pacific oceans suggest a potentially higher pool of air-resident bacteria with efficient long-range transport in the atmosphere (7).

It seems that boundaries might be drawn between the atmosphere and hydrosphere, allowing a nonrandom distribution of species between them. One case is the underrepresentation of marine bacteria in the Atlantic air, and others are airborne taxa (*e*.*g*., Firmicutes species) not detected in the water samples. Thus, we propose another constraint to the hypothesis, referring to the geo-distribution of bacteria depending not only on distance and time but also on the chemical and physical differences between these ecosystems, dictating a selective transport of different bacteria.

## Discussion

This study illustrates microbial landscape in the interface between the ocean and the atmospheric boundary layer and their spatial diversity and suggests mechanisms of dispersal at both local and global scales (Fig. 5). We observed a clear difference between the atmospheric and oceanic microbiomes, across thousands of kilometers in the North Atlantic and western Pacific oceans, with a highly variable airborne microbial composition, even with air masses spending more than 120 hours over the open ocean (Fig. 5A), compared to a more stable, and homogeneous composition of surface-water microbiome, even across latitudes and different oceans (Fig. 5B). This contrast can be attributed to the orders of magnitude differences in the characteristic advection and mixing scales in the two media, with higher stability and longer mixing cycles in the ocean and rapid changes of days to weeks in the atmosphere.

**Figure 5.**
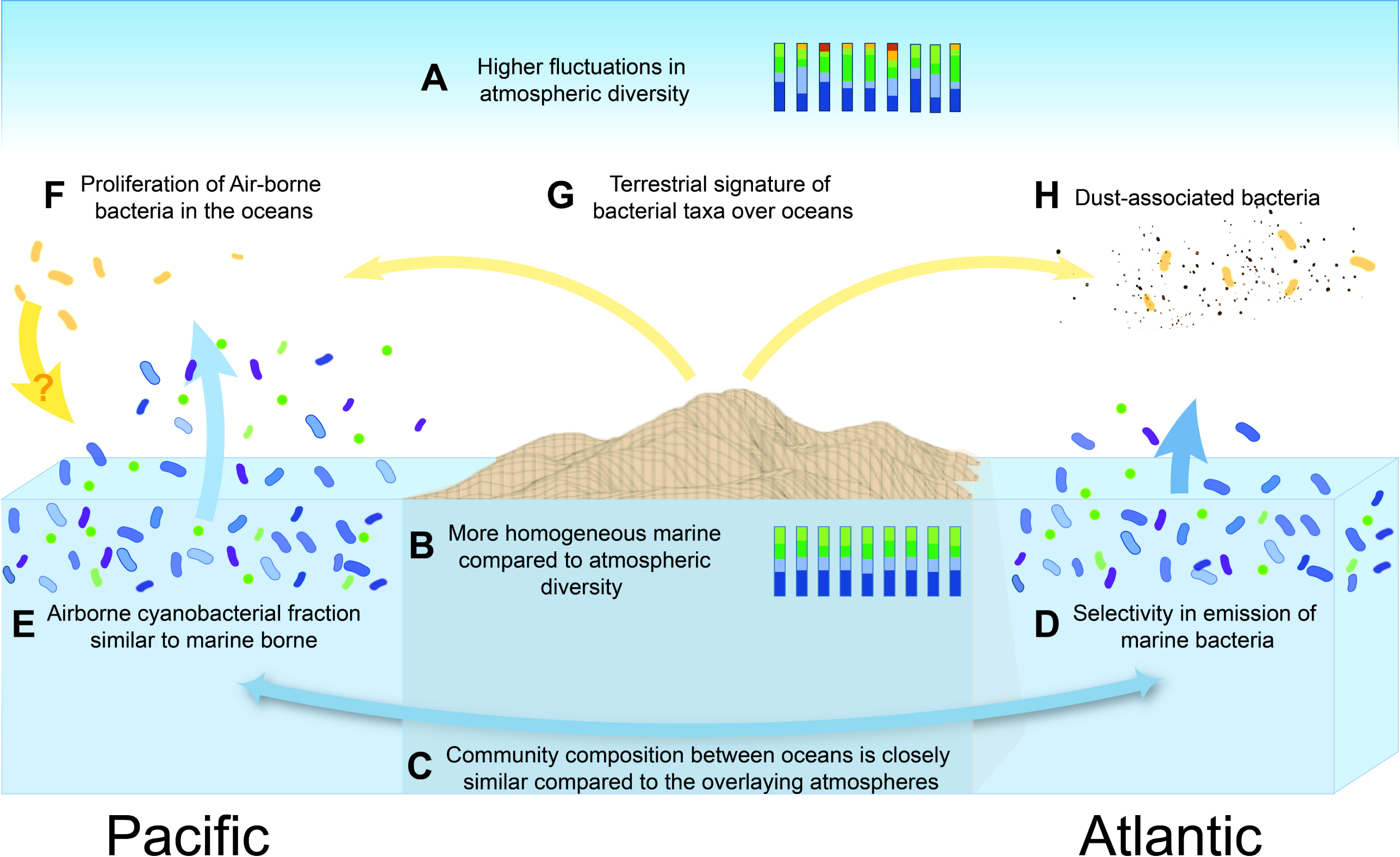
Trends and transitions in bacterial abundance and richness between marine and atmospheric environments. The main findings of this study are presented as a conceptual model: Residence time induces fluctuations in microbial composition rather than physical mixing (A, and B). The microbial compositions of similar environments (*e*.*g*., Atlantic and Pacific Oceans) are more similar than geographical relations (*e*.*g*., Pacific atmosphere and surface water; C). The marine bacterial aerosolization efficiency is different for different oceans (D), and aerosolized autotrophs kept a higher ratio *vs*. heterotrophs in the Pacific compared to the Atlantic, similarly as in the water (E). The atmospheric contribution to marine ecology was not detected in the framework of this study (F), but it cannot be excluded that difference in atmospheric microbial composition could impact deposition and surface water composition and function. A terrestrial signature was detected in a remote oceanic environment (G), with a significant contribution of dust-associated bacteria in the Atlantic atmospheric marine boundary layer (H).

Furthermore, the Atlantic and Pacific water samples showed greater resemblance to one another than to the atmosphere atop, and thousands of kilometer distant atmospheric samples shared more common taxa than with the ocean beneath (Fig. 5C). This suggests that the proximity of the sampled biomes might be less influential compared to the type of sampled environment (*i*.*e*., water *vs*. air). We additionally detected differences in the relative abundance of marine bacteria in the Pacific compared to the Atlantic atmosphere, with a significantly reduced spatial coverage in the Atlantic’s

(Fig. 5D). This observation may be linked to the dilution effect in the Atlantic AMBL due to the relatively high terrestrial-associated bacteria in this environment. Nevertheless, the consistent low appearance of these taxa in the Atlantic air, together with the high sequencing coverage (Fig. S13) of the air samples, suggests an additional mechanism. Differences in the properties of these environments and of their SMLs could impact the aerosolization efficiency of local marine bacteria, and explain the biases in their atmospheric abundance, but this hypothesis requires further validation. Although reduced in the Atlantic air, the airborne autotrophic cyanobacteria maintained a similar ratio *vs*. heterotrophs as in the surface waters in both environments (Fig. 5E).

While we found bacterial taxa associated with terrestrial environment (*e*.*g*., Firmicutes) to be significantly present in both oceans’ AMBL, we did not detect them in the oceanic surface waters, suggesting no preferential proliferation there (Fig. 5F). Since prokaryote concentrations in the atmosphere are orders of magnitude smaller compared to the ocean (∼ 10^3^-10^4^ m^-3^ in the atmosphere (4, 54) *vs*. ∼ 10^8^-10^12^ l^-1^ in the surface waters (21, 54)), sedimentation of terrestrial-originated bacteria to the ocean are not expected to induce significant change in the water composition, unless proliferating. Nevertheless, it cannot be excluded that difference in atmospheric microbial composition could impact surface water composition and function.

In addition, this study suggests that the local marine bacterial community emitted into the AMBL is enriched by long-range transported bacteria, associated with particle transport from terrestrial and anthropogenic environments (Fig. 5G). Thus, even remote locations such as the atmospheric environment of the western Pacific Ocean cannot be defined as pristine.

Finally, we associated the microbial composition of Atlantic aerosols with dust sources, making these bacteria suitable biomarkers to trace dust transport (Fig. 5H).

This study depicts a high-resolution spatial diversity of airborne bacteria and their shifts between the ocean and the atmosphere, at both local and global scales. This interplay between the ocean surface and atmospheric feedback provides new opportunities for future studies to further explore the selective properties of marine microbes within both environments and how they in turn may affect biogeochemical cycles.

## Supporting information

Spplemental file

## Acknowledgments

This research was supported by a research grant from Scott Jordan and Gina Valdez, the De Botton for Marine Science, the Yeda-Sela Center for Basic Research, and the Sustainability and Energy Research Initiative (SAERI). We are grateful to the following institutions for their financial and scientific support in Tara Pacific expedition: CNRS, PSL, CSM, EPHE, Genoscope/CEA, and France Génomique funding (ANR-10-INBS-09-08), Inserm, Université Côte d’Azur, ANR, agnès b., UNESCO-IOC, the Veolia Environment Foundation, Région Bretagne, Serge Ferrari, Billerudkorsnas, Amerisource Bergen Company, Lorient Agglomeration, Oceans by Disney, the Prince Albert II de Monaco Foundation, L’Oréal, Biotherm, France Collectivités, Kankyo Station, Fonds Français pour l’Environnement Mondial (FFEM), Etienne BOURGOIS, and the Tara Foundation teams and crew. The authors particularly thank Dr. Serge Planes, Prof. Denis Allemand, and the Tara Pacific consortium for their management and coordination. The authors gratefully acknowledging Prof. Daniel Sher for providing Prochlorococcus cultures for qPCR quantification, and Dr. Flora Vincent for her helpful, critical feedback on the manuscript. NLY acknowledges support from the Women Bridging position and the Sustainability and Energy Research Initiative (SAERI), Weizmann Institute of Science. YR acknowledges support from the Israel Science Foundation (grant #236/16). SSu acknowledges support from the ETH and Helmut Horten Foundation. This is publication number # 15 of the Tara Pacific Consortium.

## References

1. Bar-On YM, Phillips R, Milo R. The biomass distribution on Earth. Proc Natl Acad Sci U S A. 2018;115(25):6506–11.

2. Sunagawa S, Coelho LP, Chaffron S, Kultima JR, Labadie K, Salazar G, et al. Structure and function of the global ocean microbiome. Science. 2015;348(6237):1261359.

3. Ruiz-Gil T, Acuña JJ, Fujiyoshi S, Tanaka D, Noda J, Maruyama F, et al. Airborne bacterial communities of outdoor environments and their associated influencing factors. Environment International. 2020;145:106156.

4. Fröhlich-Nowoisky J, Kampf CJ, Weber B, Huffman JA, Pöhlker C, Andreae MO, et al. Bioaerosols in the Earth system: Climate, health, and ecosystem interactions. Atmos Res. 2016;182:346–76.

5. Burrows SM, Elbert W, Lawrence MG, Pöschl U. Bacteria in the global atmosphere – Part 1: Review and synthesis of literature data for different ecosystems. Atmos Chem Phys. 2009;9(23):9263–80.

6. Baas Becking LGM. Geobiologie of inleiding tot de milieukunde. 1934.

7. Martiny JBH, Bohannan BJM, Brown JH, Colwell RK, Fuhrman JA, Green JL, et al. Microbial biogeography: putting microorganisms on the map. Nat Rev Microbiol. 2006;4(2):102–12.

8. Martin K, Schmidt K, Toseland A, Boulton CA, Barry K, Beszteri B, et al. The biogeographic differentiation of algal microbiomes in the upper ocean from pole to pole. Nature Com. 2021;12(1):5483.

9. Rahav E, Paytan A, Mescioglu E, Galletti Y, Rosenfeld S, Raveh O, et al. Airborne Microbes Contribute to N2 Fixation in Surface Water of the Northern Red Sea. Geophys Res Lett. 2018;45(12):6186–94.

10. Tang W, Llort J, Weis J, Perron MMG, Basart S, Li Z, et al. Widespread phytoplankton blooms triggered by 2019–2020 Australian wildfires. Nature. 2021;597(7876):370–5.

11. Flores JM, Bourdin G, Altaratz O, Trainic M, Lang-Yona N, Dzimban E, et al. Tara Pacific expedition’s atmospheric measurements. Marine aerosols across the Atlantic and Pacific Oceans Overview and Preliminary results. Bull Am Meteorol Soc. 2020;101(5):536–54.

12. Gorsky G, Bourdin G, Lombard F, Pedrotti ML, Audrain S, Bin N, et al. Expanding Tara Oceans Protocols for Underway, Ecosystemic Sampling of the Ocean-Atmosphere Interface During Tara Pacific Expedition (2016–2018). Front Mar Sci. 2019;6(750).

13. Stein AF, Draxler RR, Rolph GD, Stunder BJB, Cohen MD, Ngan F. NOAA’s HYSPLIT Atmospheric Transport and Dispersion Modeling System. Bull Am Meteorol Soc. 2016;96(12):2059–77.

14. Alberti A, Poulain J, Engelen S, Labadie K, Romac S, Ferrera I, et al. Viral to metazoan marine plankton nucleotide sequences from the Tara Oceans expedition. Sci Dat. 2017;4(1):170093.

15. Parada AE, Needham DM, Fuhrman JA. Every base matters: assessing small subunit rRNA primers for marine microbiomes with mock communities, time series and global field samples. Environ Microbiol. 2016;18(5):1403–14.

16. Edgar RC. Search and clustering orders of magnitude faster than BLAST. Bioinformatics. 2010;26(19):2460–1.

17. Callahan BJ, McMurdie PJ, Rosen MJ, Han AW, Johnson AJA, Holmes SP. DADA2: High-resolution sample inference from Illumina amplicon data. Nat Methods. 2016;13(7):581–3.

18. Eisenhofer R, Minich JJ, Marotz C, Cooper A, Knight R, Weyrich LS. Contamination in Low Microbial Biomass Microbiome Studies: Issues and Recommendations. Trends Microbiol. 2019;27(2):105–17.

19. Ijaz AZ, Jeffries TC, Ijaz UZ, Hamonts K, Singh BK. Extending SEQenv: a taxa-centric approach to environmental annotations of 16S rDNA sequences. PeerJ. 2017;5:e3827–e.

20. DeLong EF, Preston CM, Mincer T, Rich V, Hallam SJ, Frigaard N-U, et al. Community Genomics Among Stratified Microbial Assemblages in the Ocean’s Interior. Science. 2006;311(5760):496.

21. Rodrigues TB, Silva AET. Molecular Diversity of Environmental Prokaryotes: CRC Press; 2016.

22. Sul WJ, Oliver TA, Ducklow HW, Amaral-Zettler LA, Sogin ML. Marine bacteria exhibit a bipolar distribution. Proc Natl Acad Sci U S A. 2013;110(6):2342–7.

23. Gat D, Mazar Y, Cytryn E, Rudich Y. Origin-Dependent Variations in the Atmospheric Microbiome Community in Eastern Mediterranean Dust Storms. Environ Sci Technol. 2017;51(12):6709–18.

24. Rahav E, Belkin N, Paytan A, Herut B. The Relationship between Air-Mass Trajectories and the Abundance of Dust-Borne Prokaryotes at the SE Mediterranean Sea. Atmosphere. 2019;10(5):280.

25. Lang-Yona N, Öztürk F, Gat D, Aktürk M, Dikmen E, Zarmpas P, et al. Links between airborne microbiome, meteorology, and chemical composition in northwestern Turkey. Sci Total Environ. 2020;725:138227.

26. Archer SDJ, Lee KC, Caruso T, King-Miaow K, Harvey M, Huang D, et al. Air mass source determines airborne microbial diversity at the ocean–atmosphere interface of the Great Barrier Reef marine ecosystem. Isme j. 2019;14:871–6.

27. Maki T, Lee KC, Kawai K, Onishi K, Hong CS, Kurosaki Y, et al. Aeolian Dispersal of Bacteria Associated With Desert Dust and Anthropogenic Particles Over Continental and Oceanic Surfaces. J Geophys Res-Atmos. 2019;124(10):5579–88.

28. Amato P, Besaury L, Joly M, Penaud B, Deguillaume L, Delort A-M. Metatranscriptomic exploration of microbial functioning in clouds. Sci Rep. 2019;9(1):4383.

29. Quaiser A, Ochsenreiter T, Lanz C, Schuster SC, Treusch AH, Eck J, et al. Acidobacteria form a coherent but highly diverse group within the bacterial domain: evidence from environmental genomics. Mol Microbiol. 2003;50(2):563–75.

30. Archer SDJ, Lee KC, Caruso T, Maki T, Lee CK, Cary SC, et al. Airborne microbial transport limitation to isolated Antarctic soil habitats. Nature Microbiol. 2019;4(6):925–32.

31. Caliz J, Triado-Margarit X, Camarero L, Casamayor EO. A long-term survey unveils strong seasonal patterns in the airborne microbiome coupled to general and regional atmospheric circulations. Proc Natl Acad Sci U S A. 2018;115(48):12229–34.

32. Oh KH, Jeong DH, Shin SH, Cho YC. Simultaneous quantification of cyanobacteria and Microcystis spp. using real-time PCR. J Microbiol Biotechnol. 2012;22(2):248–55.

33. Flombaum P, Gallegos JL, Gordillo RA, Rincón J, Zabala LL, Jiao N, et al. Present and future global distributions of the marine Cyanobacteria Prochlorococcus and Synechococcus. Proc Natl Acad Sci U S A. 2013;110(24):9824–9.

34. DeVries T, Primeau F. Dynamically and Observationally Constrained Estimates of Water-Mass Distributions and Ages in the Global Ocean. J Phys Oceanogr. 2011;41(12):2381–401.

35. Seinfeld JH, Pandis SN. Atmospheric chemistry and physics : from air pollution to climate change 2016.

36. Siller H, Rainey FA, Stackebrandt E, Winter J. Isolation and characterization of a new gram-negative, acetone-degrading, nitrate-reducing bacterium from soil, Paracoccus solventivorans sp. nov. Int J Syst Bacteriol. 1996;46(4):1125–30.

37. Lee JH, Kim YS, Choi TJ, Lee WJ, Kim YT. Paracoccus haeundaensis sp. nov., a Gram-negative, halophilic, astaxanthin-producing bacterium. Int J Syst Evol Microbiol. 2004;54(Pt 5):1699–702.

38. Liu XY, Wang BJ, Jiang CY, Liu SJ. Paracoccus sulfuroxidans sp.nov., a sulfur oxidizer from activated sludge. Int J Syst Evol Microbiol. 2006;56(Pt 11):2693–5.

39. DeLeon-Rodriguez N, Lathem TL, Rodriguez-R LM, Barazesh JM, Anderson BE, Beyersdorf AJ, et al. Microbiome of the upper troposphere: Species composition and prevalence, effects of tropical storms, and atmospheric implications. Proc Natl Acad Sci U S A. 2013;110(7):2575–80.

40. Mazar Y, Cytryn E, Erel Y, Rudich Y. Effect of Dust Storms on the Atmospheric Microbiome in the Eastern Mediterranean. Environ Sci Technol. 2016;50(8):4194–202.

41. Ivanova EP, Vysotskii MV, Svetashev VI, Nedashkovskaya OI, Gorshkova NM, Mikhailov VV, et al. Characterization of Bacillus strains of marine origin. Int Microbiol. 1999;2(4):267–71.

42. Mondol MAM, Shin HJ, Islam MT. Diversity of secondary metabolites from marine Bacillus species: chemistry and biological activity. Mar Drugs. 2013;11(8):2846–72.

43. Fierer N, Bradford MA, Jackson RB. Toward an ecological classification of soil bacteria. Ecology. 2007;88(6):1354–64.

44. Pootakham W, Mhuantong W, Yoocha T, Putchim L, Sonthirod C, Naktang C, et al. High resolution profiling of coral-associated bacterial communities using full-length 16S rRNA sequence data from PacBio SMRT sequencing system. Sci Rep. 2017;7(1):2774.

45. Mescioglu E, Rahav E, Belkin N, Xian P, Eizenga JM, Vichik A, et al. Aerosol Microbiome over the Mediterranean Sea Diversity and Abundance. Atmosphere. 2019;10(8):440.

46. Chisholm SW, Olson RJ, Zettler ER, Goericke R, Waterbury JB, Welschmeyer NA. A novel free-living prochlorophyte abundant in the oceanic euphotic zone. Nature. 1988;334(6180):340–3.

47. Chisholm SW, Frankel SL, Goericke R, Olson RJ, Palenik B, Waterbury JB, et al. Prochlorococcus marinus nov. gen. nov. sp.: an oxyphototrophic marine prokaryote containing divinyl chlorophyll a and b. Arch Microbiol. 1992;157(3):297–300.

48. Morris RM, Rappe MS, Connon SA, Vergin KL, Siebold WA, Carlson CA, et al. SAR11 clade dominates ocean surface bacterioplankton communities. Nature. 2002;420(6917):806–10.

49. Fahlgren C, Gomez-Consarnau L, Zabori J, Lindh MV, Krejci R, Martensson EM, et al. Seawater mesocosm experiments in the Arctic uncover differential transfer of marine bacteria to aerosols. Environ Microbiol Rep. 2015;7(3):460–70.

50. Wurl O, Ekau W, Landing W, Zappa C. Sea surface microlayer in a changing ocean – A perspective. Elementa-Sci Anthrop. 2017;5.

51. Michaud JM, Thompson LR, Kaul D, Espinoza JL, Richter RA, Xu ZZ, et al. Taxon-specific aerosolization of bacteria and viruses in an experimental ocean-atmosphere mesocosm. Nature Com. 2018;9(1):2017.

52. Buttigieg PL, Morrison N, Smith B, Mungall CJ, Lewis SE, the EC. The environment ontology: contextualising biological and biomedical entities. J Biomed Semant. 2013;4(1):43.

53. Kaufman YJ, Koren I, Remer LA, Tanré D, Ginoux P, Fan S. Dust transport and deposition observed from the Terra-Moderate Resolution Imaging Spectroradiometer (MODIS) spacecraft over the Atlantic Ocean. J Geophys Res-Atmos. 2005;110(D10).

54. Mayol E, Arrieta JM, Jiménez MA, Martínez-Asensio A, Garcias-Bonet N, Dachs J, et al. Long-range transport of airborne microbes over the global tropical and subtropical ocean. Nature Com. 2017;8(1):201.

55. Huang S, Hu W, Chen J, Wu Z, Zhang D, Fu P. Overview of biological ice nucleating particles in the atmosphere. Environment International. 2021;146:106197.

56. Flores JM, Bourdin G, Kostinski A, Altaratz O, Guy D, Lombard F, et al. Diel cycle of sea spray aerosol concentration over vast areas of the tropical Pacific Ocean and the Caribbean Sea. Nature Com. Accepted.

57. Jones KR, Klein CJ, Halpern BS, Venter O, Grantham H, Kuempel CD, et al. The Location and Protection Status of Earth’s Diminishing Marine Wilderness. Curr Biol. 2018;28(15):2506-12.e3.

58. Halpern BS, Frazier M, Afflerbach J, Lowndes JS, Micheli F, O’Hara C, et al. Recent pace of change in human impact on the world’s ocean. Sci Rep. 2019;9(1):11609.

59. Uetake J, Hill TCJ, Moore KA, DeMott PJ, Protat A, Kreidenweis SM. Airborne bacteria confirm the pristine nature of the Southern Ocean boundary layer. Proc Natl Acad Sci U S A. 2020.202000134.

60. de Leeuw G, Guieu Cc, Arneth A, Bellouin N, Bopp L, Boyd PW, et al. Ocean–Atmosphere Interactions of Particles. Ocean-Atmosphere Interactions of Gases and Particles. Berlin, Heidelberg: Springer Earth System Sciences; 2014. p. 171–246.

