## Supplementary material for "A cross-biomes bacterial diversity shed light on ocean-atmosphere microbial transmission": Spplemental file

### Airborne bacteria over oceans shed light on global biogeodiversity patterns

**This PDF file includes:**

Supplementary Text
Figs. S1 to S14
Tables S1 to S8
References (1 to 14)

**Supplementary Text**

Validation of DNA extraction from air filters

The yield of DNA extraction was optimized, by comparing different extraction methods, based on our previous studies, and estimating the sensitivity of bacterial detection determining the minimal detected levels (*1-3*). We verified no biases in sequencing coverages by extracting DNA from synthetic bacterial community-spiked filters. As can be seen in Figure S7, we reached a detection limit of down to 100 cells of bacteria per filter.

Validations for no boat-originated contamination

The possible local emission from the R/V Tara vessel and crew was tested and excluded through various approaches.

To validate that our population is not contaminated with human or ship-emitted bacteria, we compared day to night samples, assuming that if contamination affects the airborne bacterial community, this fraction will be higher during the daytime, as water sampling and other activities take place during this time. We, therefore, also expect higher aerosolization of dust from the deck. For ENVO analysis, there is no significant difference ($p$-$\text{Value}_{\text{Atlantic}}$ = 0.94, and 0.06, and $p$-$\text{Value}_{\text{Pacific}}$ = 0.79, and 0.82 for Terrestrial and Anthropogenic environments, with Mann-Whitney two-sample test) during the day and night samples (Fig. S9).

To make sure no artificial sea spray was generated by the boat, we initially checked the concentration of sea spray with wind direction relative to the boat. In Figure S10 below, a wind-rose plot of particle concentrations *vs*. wind direction and speed from the Atlantic transect is presented. As can be seen, no biases toward the front of the boat (where we expect to see the highest contribution of artificial particles generated by the boat) are observed.

Decontamination of the atmospheric samples' amplicon sequencing data

The analysis of microbial population was performed only after a thorough decontamination procedure, to mitigate putative contamination in sequence libraries from the air samples with ultra-low DNA biomass as follows:

Suspiciously frequent and/or prevalent ASV were identified using the [isContaminant] function of the R package *decontam* (*4*) and removed from the dataset if identified by either frequency or prevalence statistical threshold (0.1, and 0.5, respectively).

Although the blank bacterial population was very different from the airborne population (See Figure S8), all ASV encountered in any of the sampling blanks was removed.

Putative contaminants that may originate from human activity, based on previous studies of low biomass samples,(*5, 6*) were subtracted at the genus level (even if not found in our control samples). Removed bacterial genera included: *Bacteroides*, *Bifidobacterium*, *Corynebacterium*, *Cutibacterium*, *Escherichia*/*Shigella*, *Faecalibacterium*, *Haemophilus*, *Klebsiella Lactobacillus*, *Listeria*, *Moraxella*, *Neisseria*, *Porphyromonas*, *Prevotella*, *Salmonella*, *Staphylococcus*, *Streptococcus* and *Veillonella*. Other ASVs affiliated with potential reagent contaminant genera (*Acidovorax*, *Acinetobacter*, *Afipia*, *Aquabacterium*, *Bacillus*, *Bradyrhizobium*, *Burkholderia*, *Clostridium*, *Dysgonomonas*, *Enterobacter*, *Kocuria*, *Methylobacterium*, *Pseudomonas*, *Ralstonia*, *Renibacterium*, *Rhizobacter*, *Romboutsia*, *Sphingobium*, *Sphingomonas*, *Tardiphaga*, *Turicibacter*, *Variovorax*) were manually inspected and removed if showed strong inverse correlation to DNA concentration in samples (as those genera are also found in different environments). Overall, the multi-step decontamination process identified 1,286 bacterial ASVs as suspected contaminants that were removed from the dataset.

Correcting biomass for particle losses due to sampling tubing

Within the atmospheric marine boundary layer (AMBL), Lewis & Schwartz (2013) showed that the dependence of aerosol concentrations above 10 m with height for diameters < 10 µm is less than 5% for winds as low as 5 m $s^{-1}$ (and even less than that for higher winds) up to the first ~30 m above the sea surface (*7*) . Additionally, it is also shown that the AMBL typically mixes within an hour (*7*) . A characteristic time is in the order of 20 min for a 500 m AMBL. This implies that the 15m difference in the inlet height will not affect the composition of particles being measured, as the particles are homogeneously distributed in the sampled air.
Nevertheless, the length of the sampling tubing could impact the particle concentration sampled from the air as the long tubing leads to losses of particles. To account for the losses in both setups, we calculated the particles losses as a function of size using the Particle loss calculator (*8*). Figure S11 (adapted from Flores et al. 2020, Appendix A) (*9*) shows the particle losses through both inlet setups. For particles ~1 µm, the losses in the Pacific are calculated to be about 60% and in the Atlantic, about 30%.
It is important to note that although the biomass is reduced, the abundance of bacterial population should not be different, as the particles are homogeneously distributed in the sampled air.
This led to a reduced, but still significant difference between the sampled biomass in the Atlantic compared to the Pacific, with a factor of 4.98 ± 4.21 less in the Pacific (*t*-test, *p*-value < 0.0001).
We also calculated the aerosol concentration for diameter between 1 - 3 µm using online measurements of the aerosol concentrations (*9*), and as can be seen in Figure S12.
These two separate measurements are consistent with each other, and show that the higher mass we measured in the Atlantic is not due to neither the inlet height difference nor possible contamination from the deck of *Tara*.

Correcting biomass for particle losses due to sampling tubing

The bacterial genome concentration obtained from Qubit fluorescence reads was converted to DNA biomass per $m^3$ as follows equation S1:

$$\text{(S1)} \quad C_{DNA}\left[\frac{pg}{m^3}\right] = \frac{C_{DNA}\left[\frac{pg}{\mu l}\right]\cdot V_{DNA}[\mu l]}{T[h]\cdot 60[\frac{min}{h}]\cdot V_{Air}[\frac{l}{min}]\cdot 0.001[\frac{m^3}{l}]} \cdot Tubing\ factor$$

Where $C_{DNA}\left[\frac{pg}{\mu l}\right]$ is the values obtained by Qubit reads, $V_{DNA}[\mu l]$ is the volume of DNA solution extracted, $T[h]$ is the air sampling time, and $V_{Air}[\frac{l}{min}] \cdot 0.001[\frac{m^3}{l}]$ is the air sampling volume. The tubing factor is the particle lose rates for particles ~1 µm, in a given tubing length, calculated using the Particle loss Calculator (*8*).

Comparison of microbial composition from filters with different sampling duration

Although DNA molecules are very stable under dry conditions,(*10, 11*) we examined the possible under-representation of certain bacteria from filters sampled over long time periods by comparing the community composition of 12 hrs. filter samples (the majority of the samples) and >20 hrs. filter samples (Table S1, for filter information). We did not find significant difference in community composition between the two groups (ANOSIM, *p*-value = 0.145), as shown in Fig. S14.

Detailed description of the Environmental Ontology analysis and the SEQenv pipeline

Environmental descriptive terms were extracted from the closest matches (97% identity) using the SEQenv pipeline for Python (version 1.3.0) with default parameters(*12*, *13*) and ENVO terms.(*14*) The input data included FASTA files of sequences after quality control check and removal of blank contaminants per each sample, to be compared to highly similar sequences from public repositories (such as SILVA and GenBank, using the NCBI nucleotide data base). Subsequently, from each of those records, text fields carrying environmental context information have been extracted. Existing links to PubMed abstracts were also followed and the relevant abstracts collected. The SEQenv pipeline is capable of annotating genetic sequences based on environment descriptive terms occurring within their records and/or in relevant literature. Followingly, after all relevant pieces of text for each matching sequence have been gathered, they were processed by a text mining module capable of identifying any Environment Ontology (ENVO) environment descriptive terms mentioned therein. The identified ENVO terms, and their mention frequency were then subjected to clustering analysis and multivariate statistics. As a result, tag-clouds and heatmaps of environment descriptive terms characterizing different set of sequences (e.g., originating from different samples) were generated.

In the current study, the terms were further clustered into five main groups: marine, terrestrial, fresh water, anthropogenic, and unclassified, as detailed in Table S7.

While the ENVO analysis is capable of comparing our identified taxa to the current reported datasets, it may potentially exclude unreported environments, where a specific bacterium may potentially be found.

**Fig. S1.**

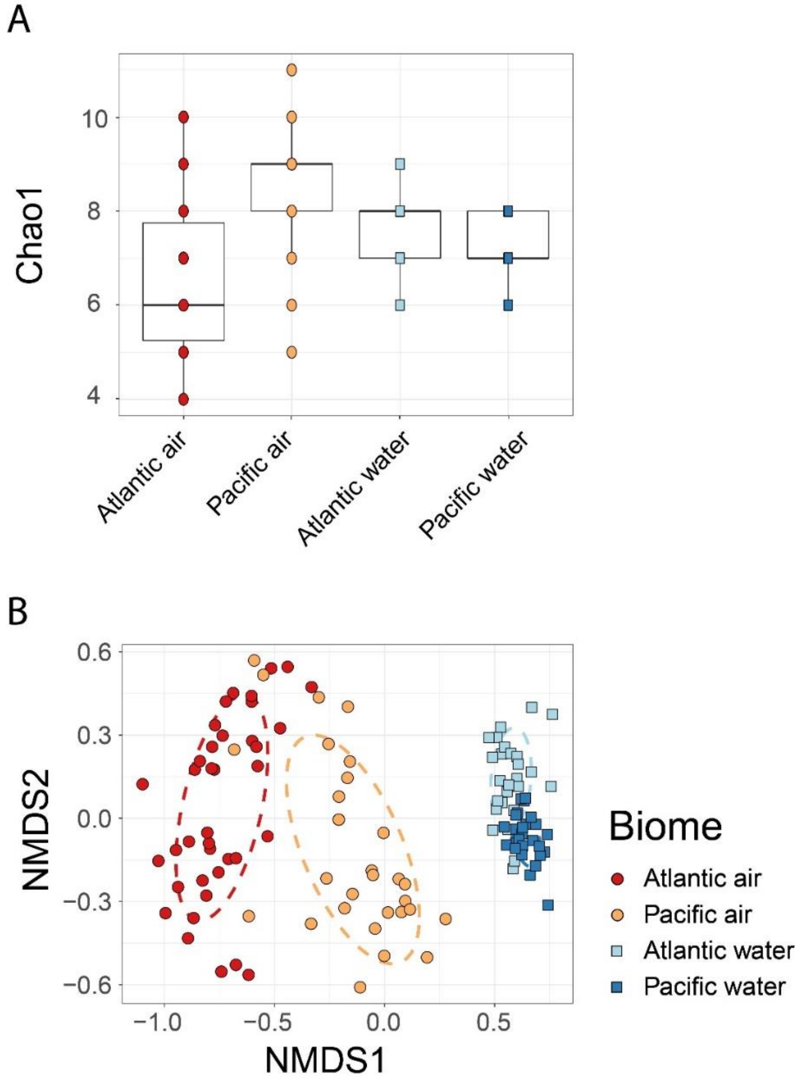


**Fig. S1. Descriptive statistics for bacterial phyla in the Pacific and Atlantic oceans and air**. The bacterial phyla richness (based on the Chao 1 estimator) in the four environments (A) and nonmetric multidimensional scaling (NMDS) ordination, with Bray-Curtis dissimilarity metrics (B) with 95% confidence ellipses.

**Fig. S2.**

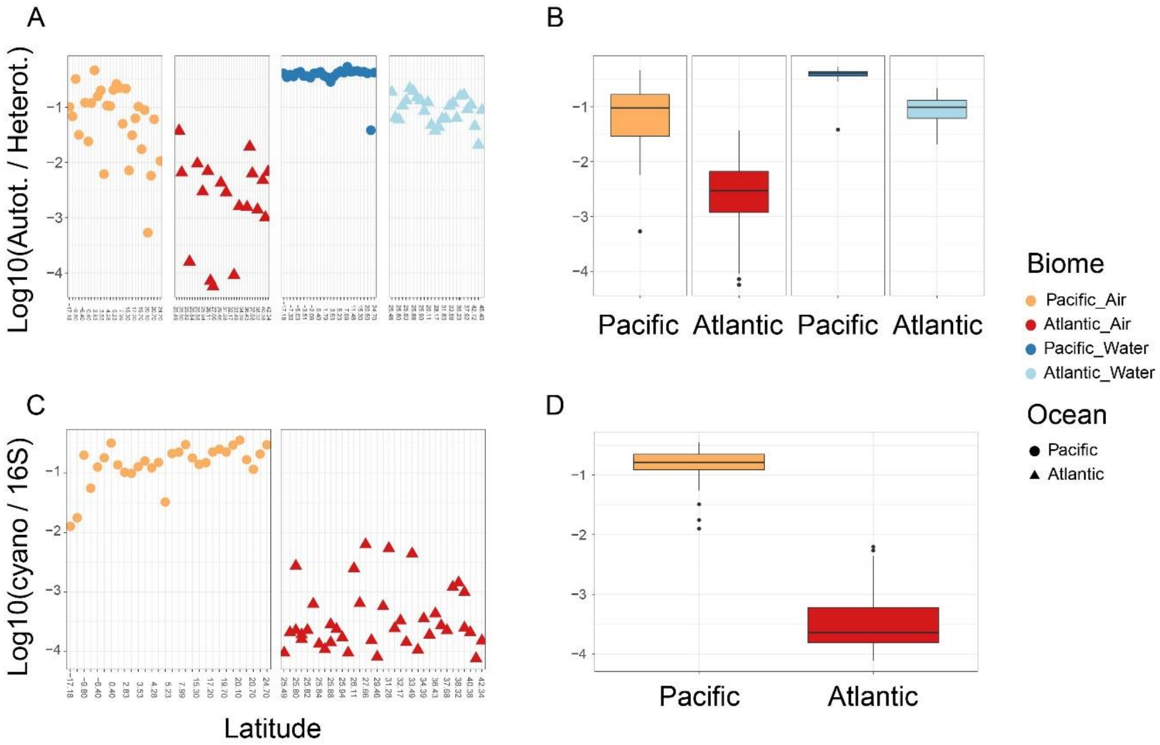


**Fig. S2. Aerial signature of marine productivity.** The ratio between autotroph ASVs to heterotrophs (A) with latitudinal change and the average values (B) for the Atlantic air, Pacific Air, Atlantic water and Pacific water samples. The scaled cyanobacterial gene copy number to universal bacteria 16S with latitudinal change (C) and the average values (D) for the Pacific and Atlantic transects. Missing datapoints in (A) are due to the absence of identified autotrophs in those samples.

**Fig. S3.**

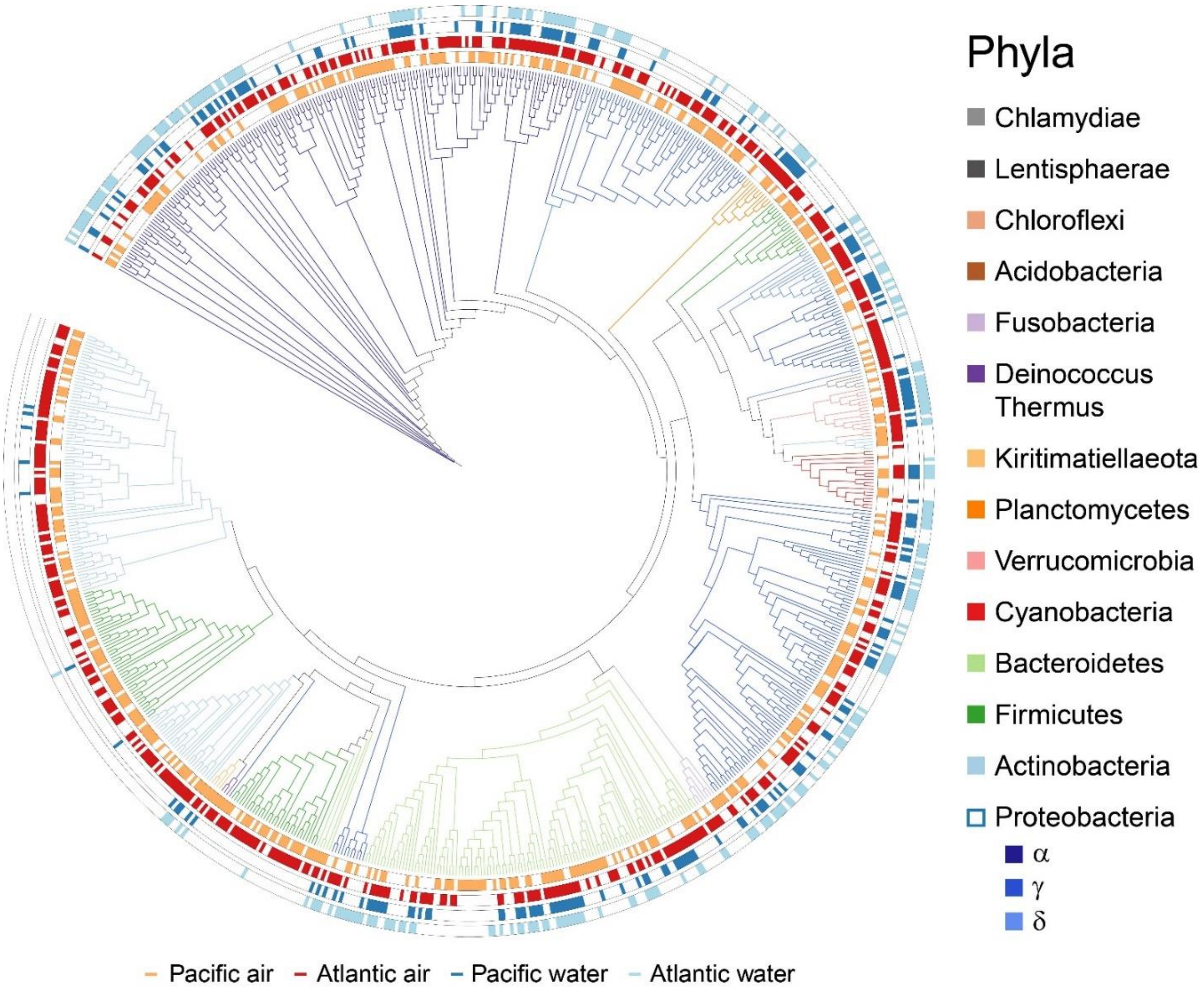


**Fig. S3: Phylogenetic overview of the ocean-atmosphere bacterial community.** Most likelihood phylogenetic tree constructed based on bacterial taxa observed in more than one sample. Colored bars at the outer circles indicate occurrences of specific taxa in the different biomes: Pacific air (Orange), Atlantic air (red), Pacific water (dark blue), and Atlantic water (light blue).

**Fig. S4.**

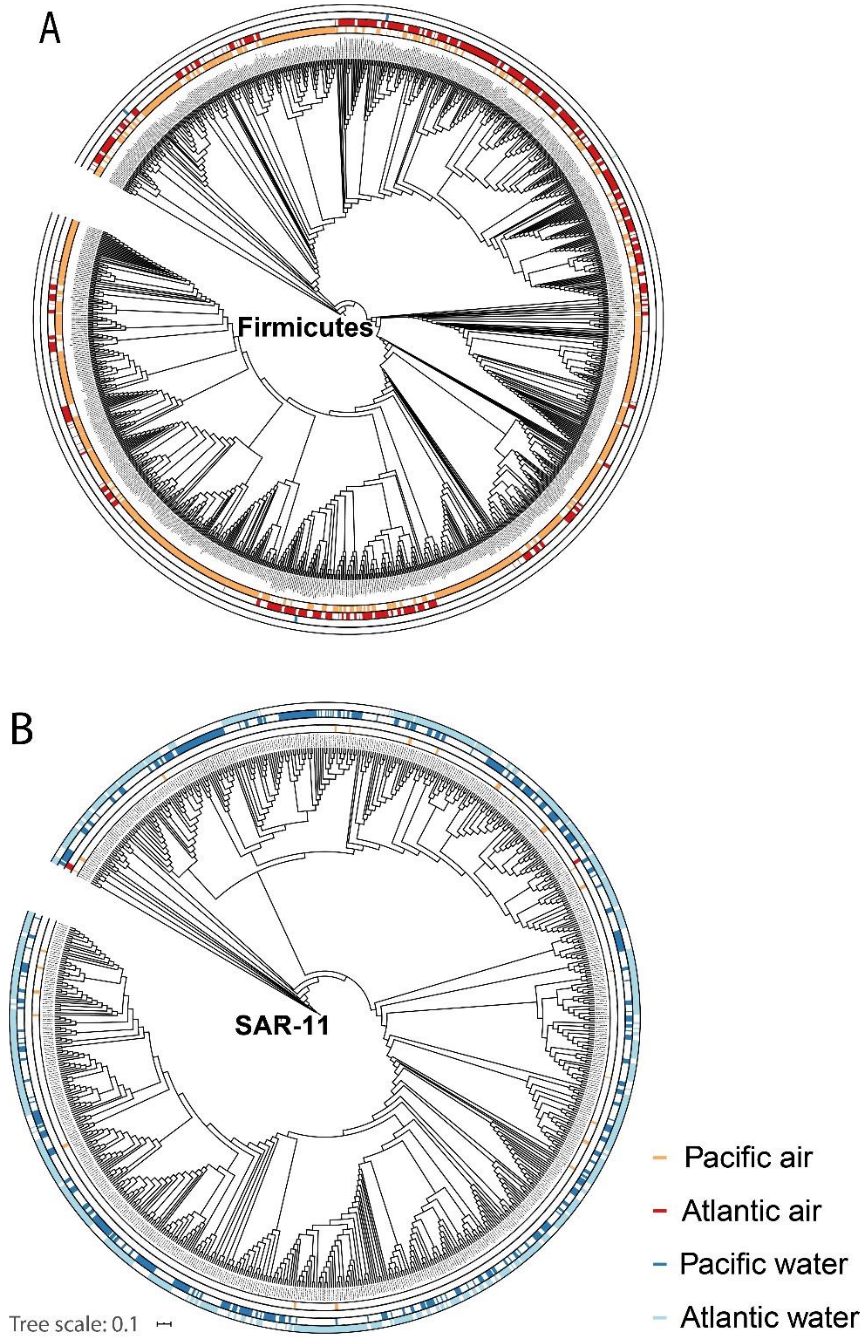


**Fig. S4. Phylogenetic overview of targeted bacterial community.** Most likelihood phylogenetic tree constructed based on Firmicutes (A) and SAR-11 clade (B) ASVs. Colored bars at the outer circles indicate occurrences of specific taxa in the different biomes: Pacific air (Orange), Atlantic air (red), Pacific water (dark blue), and Atlantic water (light blue).

**Fig. S5.**

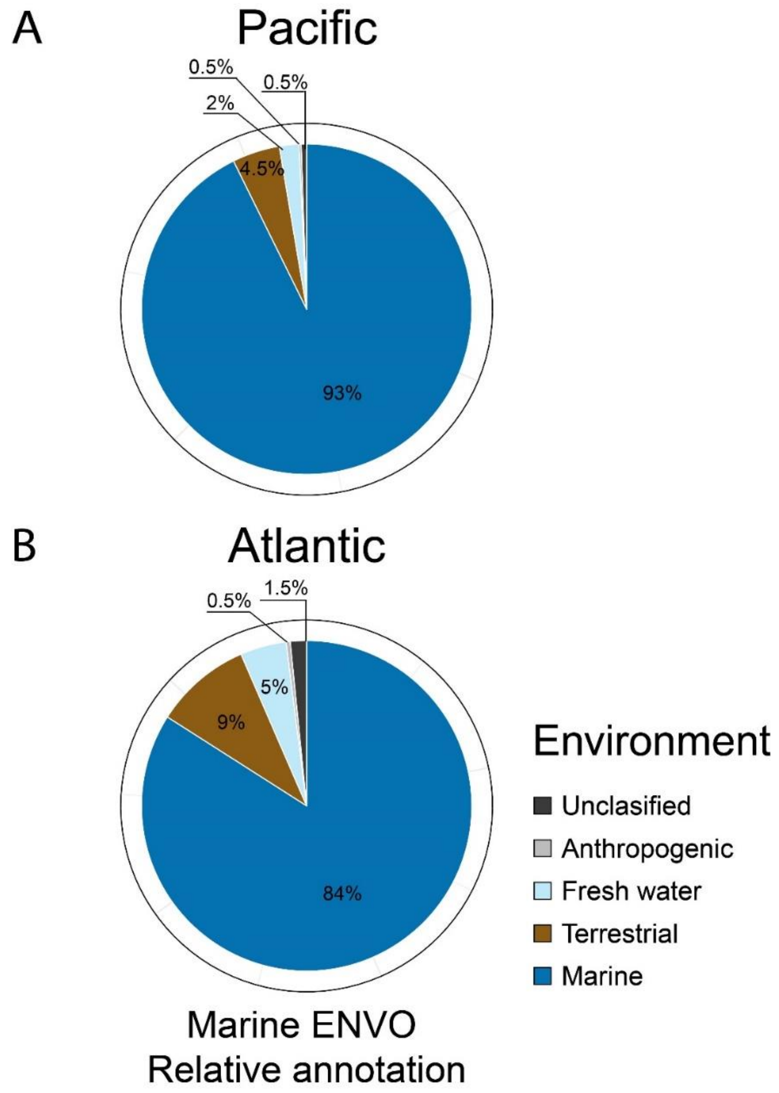


**Fig. S5. Environmental distribution of surface water-borne bacterial communities.** The average environmental ontology (ENVO) distributions of the surface water sample ASVs are presented in A and B for the Pacific and Atlantic environments, respectively.

**Fig. S6.**

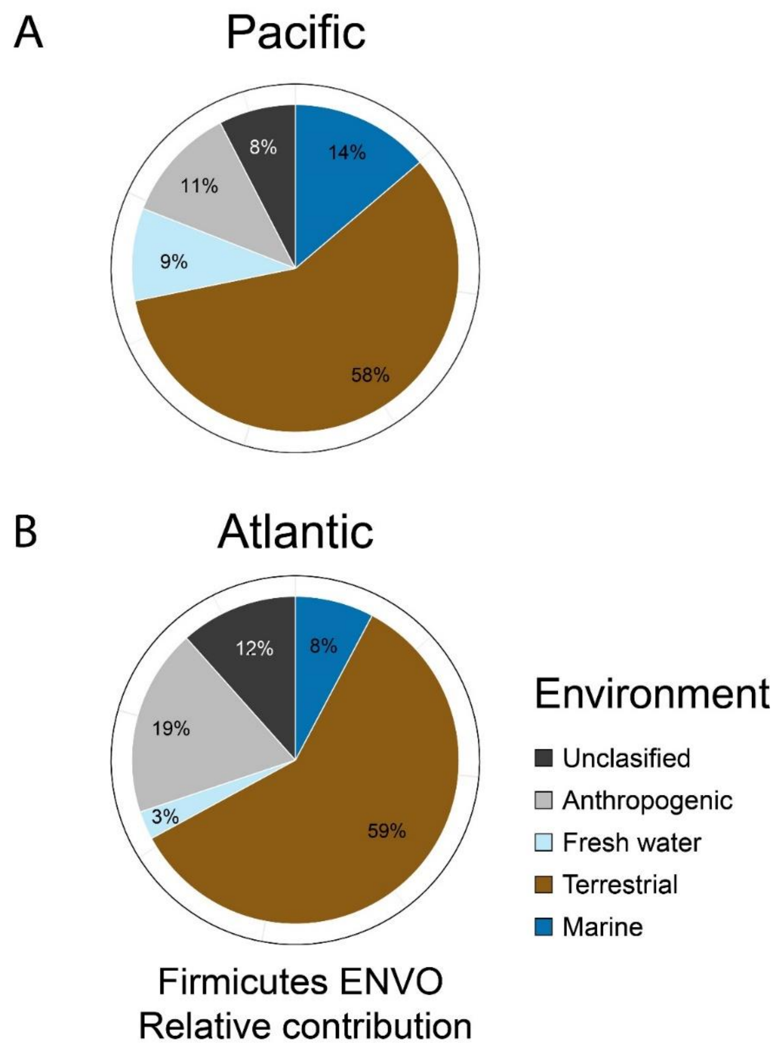


**Fig. S6. The environmental distribution of targeted bacterial communities.** Environmental ontology (ENVO) relative contribution of airborne Firmicutes ASVs clustered into five main groups (detailed terms are listed in Table S6) for the Pacific (A) and Atlantic (B) based on the average abundance in each transect.

Fig. S7.

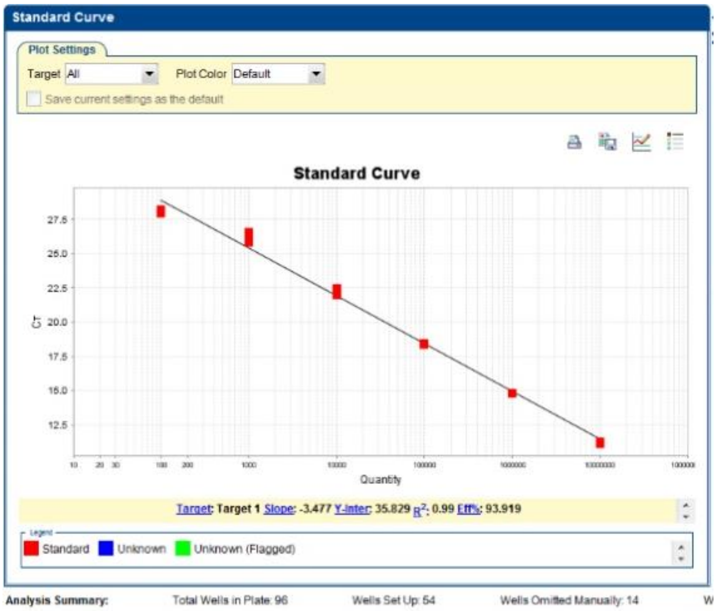


**Fig. S7. Bacterial 16S gene calibration curve.** The sensitivity (100 bacterial cells) and amplification efficiency (93.9%) of 16S region of bacterial genome extracted from a dilution series of filters with known amounts of bacterial cells, obtained using qPCR analysis. Dilutions are in triplicates.

**Fig. S8.**

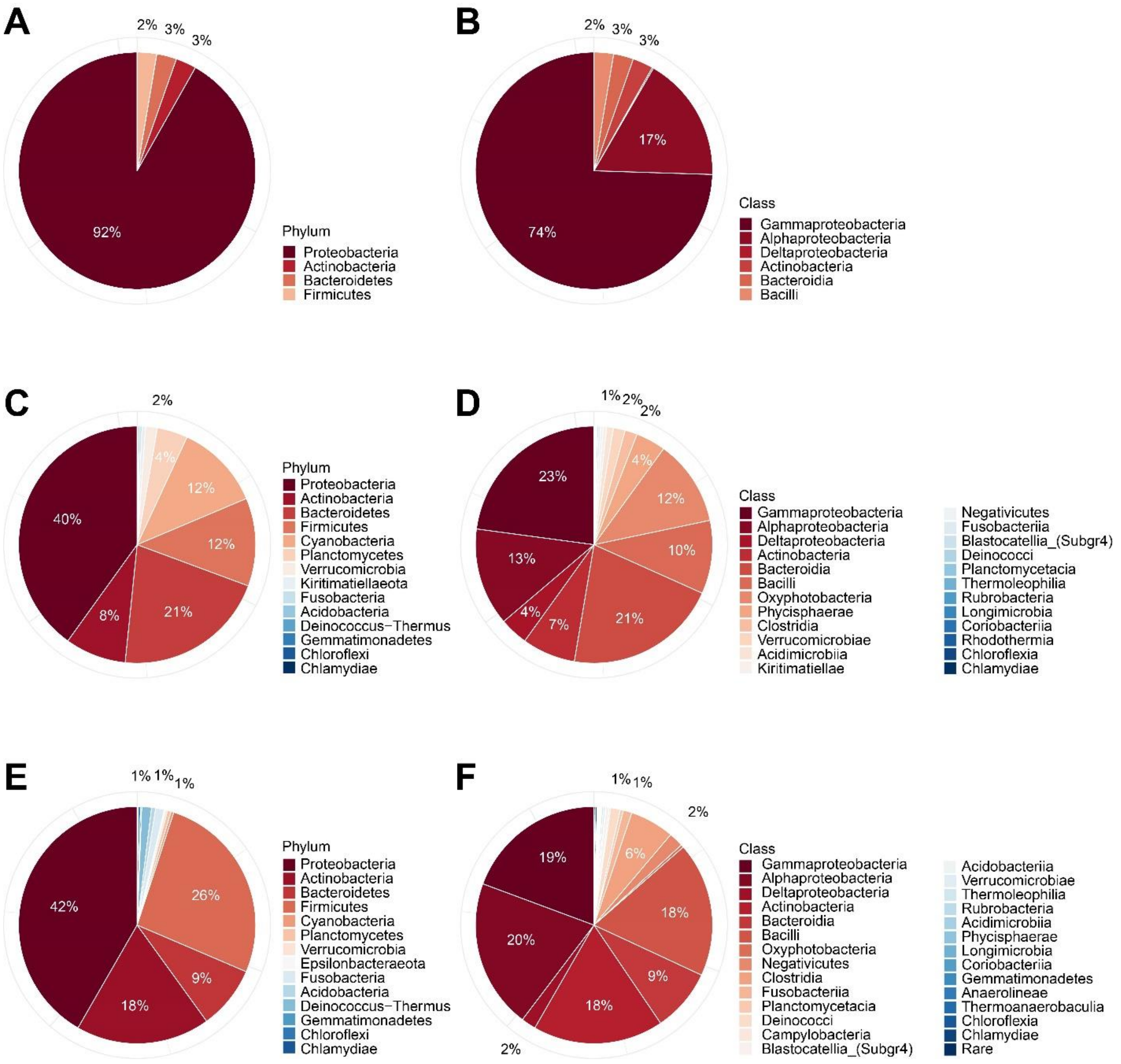


**Fig. S8. Blank filters bacterial community compared to the air filter community.** The relative distribution of blank filters phylum (A) and Class (B), compared to the Pacific (C, and D, respectively) and Atlantic (E, and F, respectively) air filter communities.

**Fig. S9.**

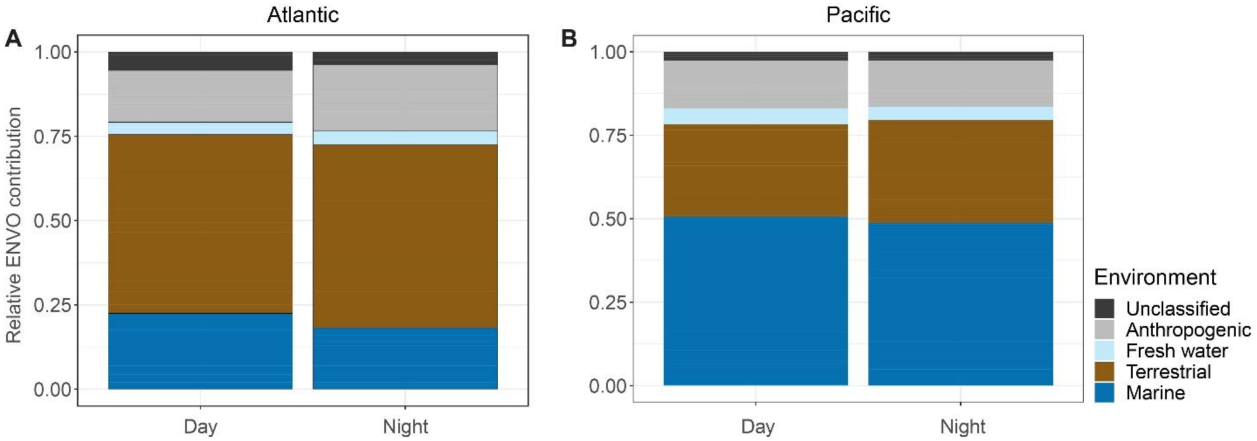


**Fig. S9. Similarity in day and night environmental ontology analysis for air samples.** Day- and night-sampled filters in the Atlantic (A) and Pacific (B) clustered into environmental ontology environments, to show no significant difference between terrestrial and anthropogenic contribution from the boat to the aerosolized fraction.

**Fig. S10.**

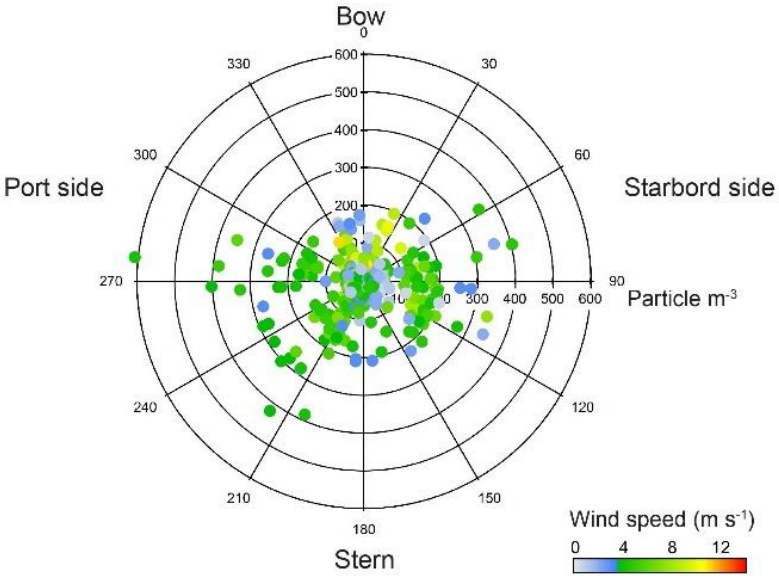


**Fig. 10. Boat-originate pollution does not impact on air samples due to wind direction in the Atlantic transect.** Aerosol concentration *vs.* angle of wind relative to the R/V Tara (particle size range 0.25 < D < 32 µm). Wind speed is denoted by the color scale.

**Fig. S11.**

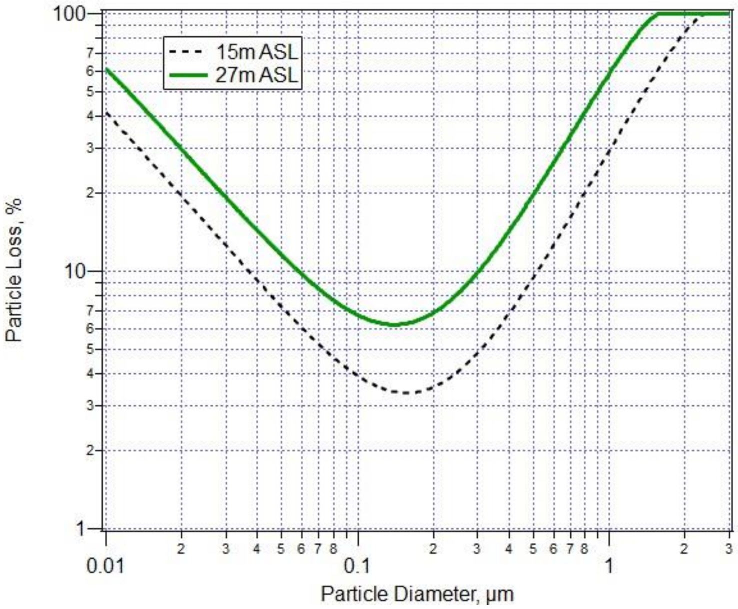


**Fig. S11. Theoretical particle loss through the inlets installed on the back stay of Tara.** The calculations were done using the Particle loss Calculator.(*8*) The curves represent the losses at the two inlet heights above sea levels (ASL).

**Fig. S12.**

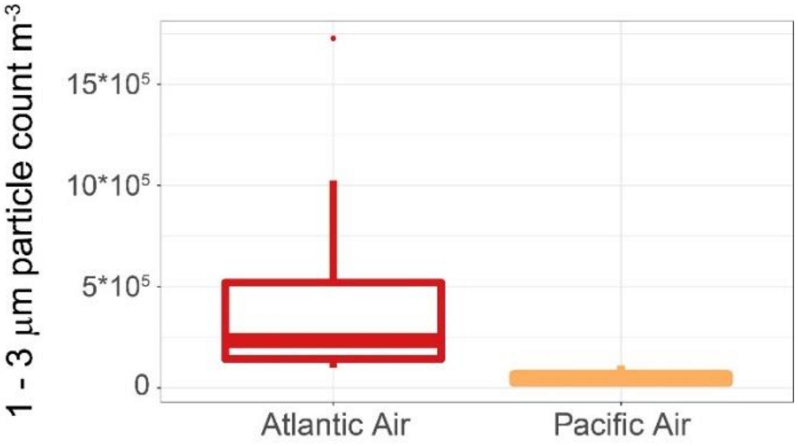


**Fig. S12.** Box plot of the Atlantic and Pacific aerosol (1 – 3 µm) concentrations after correction for particle losses due to inlet tube length.

**Fig. S13.**

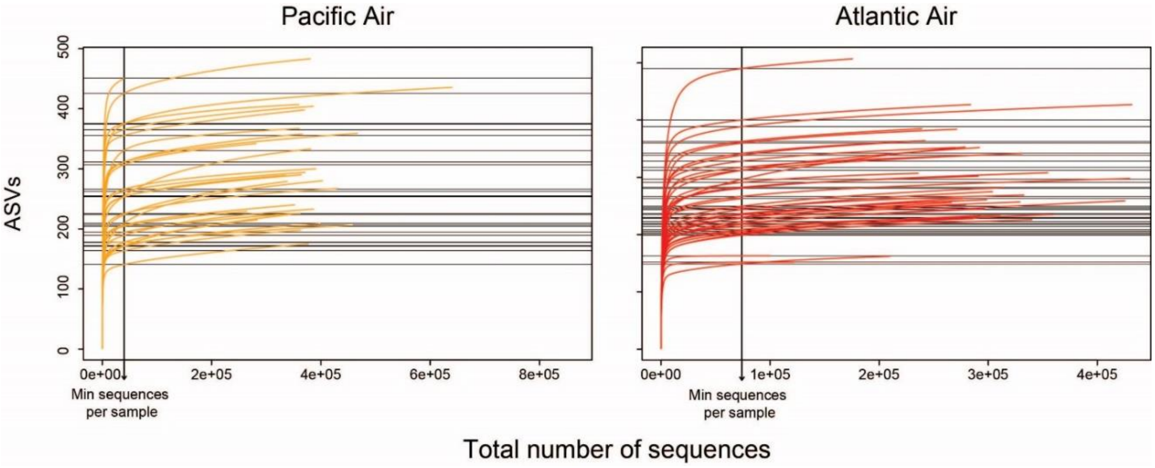


**Fig. S13. Airborne microbiome rarefaction curves.** The rarified ASVs per number of sequences in each sample for the Pacific (Orange) and the Atlantic (red) air. The minimum number of sequences per sample is denoted.

**Fig. S14.**

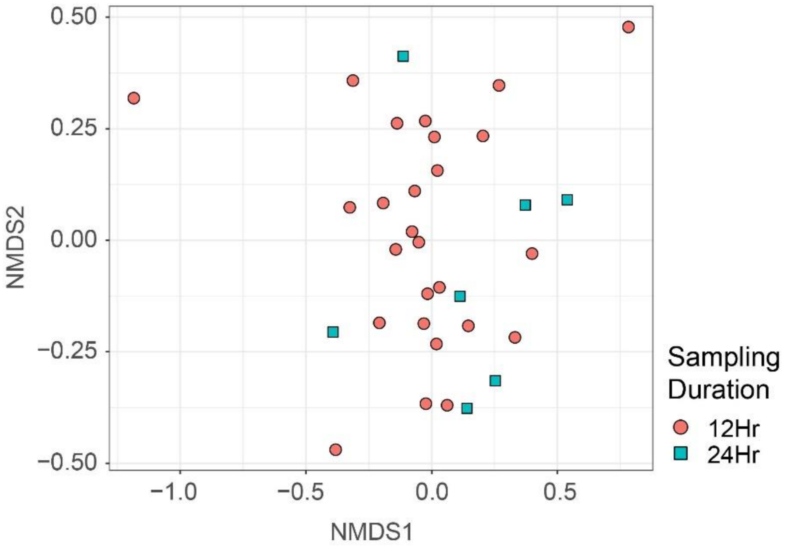


**Fig. S14. No impact of sampling duration on the airborne microbiome in the Atlantic transect**. Nonmetric multidimensional scaling distance matrix (NMDS) comparing microbial composition of 12 hrs. (pink circles) and 24 hrs. (green squares) sampling duration.

**Table S1. Atmosphere and ocean surface water filter samples.** The metadata of filter samples including ID, barcode, geographic region, Environment, sampling date, time and mean latitude

| Sample ID | Filter Barcode | Geographic region | Environment | Date, air sampling duration (hh:mm) | Lat |
|---|---|---|---|---|---|
| TARA_G.0000028_METAB.16S | G-0000028 | Atlantic | Air | 20160531, 10:39 | 43.20 |
| TARA_G.0000031_METAB.16S | G-0000031 | Atlantic | Air | 20160601, 12:47 | 42.34 |
| TARA_G.0000032_METAB.16S | G-0000032 | Atlantic | Air | 20160601, 12:31 | 41.36 |
| TARA_G.0000034_METAB.16S | G-0000034 | Atlantic | Air | 20160602, 11:48 | 40.38 |
| TARA_G.0000036_METAB.16S | G-0000036 | Atlantic | Air | 20160602, 11:10 | 39.37 |
| TARA_G.0000038_METAB.16S | G-0000038 | Atlantic | Air | 20160603, 22:14 | 38.32 |
| TARA_G.0000039_METAB.16S | G-0000039 | Atlantic | Air | 20160604, 11:43 | 37.84 |
| TARA_G.0000060_METAB.16S | G-0000060 | Atlantic | Air | 20160604, 11:34 | 37.68 |
| TARA_G.0000061_METAB.16S | G-0000061 | Atlantic | Air | 20160605, 11:10 | 37.29 |
| TARA_G.0000068_METAB.16S | G-0000068 | Atlantic | Air | 20160607, 12:55 | 36.43 |
| TARA_G.0000069_METAB.16S | G-0000069 | Atlantic | Air | 20160607, 12:34 | 35.36 |
| TARA_G.0000092_METAB.16S | G-0000092 | Atlantic | Air | 20160608, 23:21 | 34.39 |
| TARA_G.0000094_METAB.16S | G-0000094 | Atlantic | Air | 20160609, 11:34 | 33.49 |
| TARA_G.0000096_METAB.16S | G-0000096 | Atlantic | Air | 20160609, 11:13 | 33.65 |
| TARA_G.0000098_METAB.16S | G-0000098 | Atlantic | Air | 20160610, 12:49 | 33.10 |
| TARA_G.0000100_METAB.16S | G-0000100 | Atlantic | Air | 20160610, 11:10 | 32.17 |
| TARA_G.0000102_METAB.16S | G-0000102 | Atlantic | Air | 20160611, 10:56 | 31.70 |
| TARA_G.0000104_METAB.16S | G-0000104 | Atlantic | Air | 20160611, 12:24 | 31.28 |
| TARA_G.0000106_METAB.16S | G-0000106 | Atlantic | Air | 20160612, 22:37 | 30.39 |
| TARA_G.0000122_METAB.16S | G-0000122 | Atlantic | Air | 20160613, 13:14 | 29.46 |
| TARA_G.0000124_METAB.16S | G-0000124 | Atlantic | Air | 20160613, 11:20 | 28.59 |
| TARA_G.0000126_METAB.16S | G-0000126 | Atlantic | Air | 20160614, 23:39 | 27.66 |
| TARA_G.0000128_METAB.16S | G-0000128 | Atlantic | Air | 20160615, 24:07 | 26.48 |
| TARA_G.0000130_METAB.16S | G-0000130 | Atlantic | Air | 20160616, 24:00 | 25.83 |
| TARA_G.0000131_METAB.16S | G-0000131 | Atlantic | Air | 20160617, 23:41 | 25.49 |
| TARA_G.0000159_METAB.16S | G-0000159 | Atlantic | Air | 20160619, 24:29 | 25.88 |
| TARA_G.0000166_METAB.16S | G-0000166 | Atlantic | Air | 20160621, 11:44 | 25.79 |
| TARA_G.0000175_METAB.16S | G-0000175 | Atlantic | Air | 20160620, 11:01 | 25.85 |
| TARA_G.0000178_METAB.16S | G-0000178 | Atlantic | Air | 20160621, 12:36 | 25.80 |
| TARA_G.0000185_METAB.16S | G-0000185 | Atlantic | Air | 20160622, 12:41 | 25.83 |
| TARA_G.0000209_METAB.16S | G-0000209 | Atlantic | Air | 20160625, 11:38 | 25.95 |
| TARA_G.0000210_METAB.16S | G-0000210 | Atlantic | Air | 20160625, 10:50 | 25.90 |
| TARA_G.0000212_METAB.16S | G-0000212 | Atlantic | Air | 20160626, 11:12 | 25.81 |
| TARA_G.0000219_METAB.16S | G-0000219 | Atlantic | Air | 20160626, 11:48 | 25.94 |
| TARA_G.0000220_METAB.16S | G-0000220 | Atlantic | Air | 20160627, 11:17 | 26.11 |
| TARA_G.0000227_METAB.16S | G-0000227 | Atlantic | Air | 20160627, 12:42 | 25.82 |
| TARA_G.0000228_METAB.16S | G-0000228 | Atlantic | Air | 20160628, 11:16 | 25.80 |
| TARA_OA.0000025_METAB.16S | OA-0000025 | Atlantic | Water | 20160530 | 45.40 |
| TARA_OA.0000037_METAB.16S | OA-0000037 | Atlantic | Water | 20160531 | 43.88 |
| TARA_OA.0000046_METAB.16S | OA-0000046 | Atlantic | Water | 20160601 | 42.72 |
| TARA_OA.0000056_METAB.16S | OA-0000056 | Atlantic | Water | 20160602 | 40.91 |

| | | | | | |
|---|---|---|---|---|---|
| TARA_OA.0000067_METAB.16S | OA-0000067 | Atlantic | Water | 20160604 | 37.92 |
| TARA_OA.0000084_METAB.16S | OA-0000084 | Atlantic | Water | 20160606 | 36.27 |
| TARA_OA.0000094_METAB.16S | OA-0000094 | Atlantic | Water | 20160607 | 36.23 |
| TARA_OA.0000106_METAB.16S | OA-0000106 | Atlantic | Water | 20160608 | 34.94 |
| TARA_OA.0000116_METAB.16S | OA-0000116 | Atlantic | Water | 20160609 | 33.58 |
| TARA_OA.0000139_METAB.16S | OA-0000139 | Atlantic | Water | 20160610 | 33.54 |
| TARA_OA.0000150_METAB.16S | OA-0000150 | Atlantic | Water | 20160611 | 31.83 |
| TARA_OA.0000156_METAB.16S | OA-0000156 | Atlantic | Water | 20160613 | 29.52 |
| TARA_OA.0000170_METAB.16S | OA-0000170 | Atlantic | Water | 20161614 | 28.17 |
| TARA_OA.0000176_METAB.16S | OA-0000176 | Atlantic | Water | 20160615 | 26.69 |
| TARA_OA.0000191_METAB.16S | OA-0000191 | Atlantic | Water | 20160616 | 26.11 |
| TARA_OA.0000206_METAB.16S | OA-0000206 | Atlantic | Water | 20161617 | 25.48 |
| TARA_OA.0000216_METAB.16S | OA-0000216 | Atlantic | Water | 20160618 | 25.60 |
| TARA_OA.0000228_METAB.16S | OA-0000228 | Atlantic | Water | 20160620 | 25.90 |
| TARA_OA.0000242_METAB.16S | OA-0000242 | Atlantic | Water | 20160621 | 25.80 |
| TARA_OA.0000259_METAB.16S | OA-0000259 | Atlantic | Water | 20160622 | 25.81 |
| TARA_OA.0000266_METAB.16S | OA-0000266 | Atlantic | Water | 20160623 | 25.94 |
| TARA_OA.0000278_METAB.16S | OA-0000278 | Atlantic | Water | 20160624 | 25.89 |
| TARA_OA.0000292_METAB.16S | OA-0000292 | Atlantic | Water | 20160625 | 25.88 |
| TARA_OA.0000301_METAB.16S | OA-0000301 | Atlantic | Water | 20160626 | 25.83 |
| TARA_OA.0000315_METAB.16S | OA-0000315 | Atlantic | Water | 20160627 | 25.87 |
| TARA_OA.0000329_METAB.16S | OA-0000329 | Atlantic | Water | 20160706 | 24.26 |
| TARA_G.0001068_METAB.16S | G-0001068 | Pacific | Air | 20170430, 12:28 | 24.70 |
| TARA_G.0001099_METAB.16S | G-0001099 | Pacific | Air | 20170503, 12:15 | 21.10 |
| TARA_G.0001100_METAB.16S | G-0001100 | Pacific | Air | 20170503, 11:52 | 20.70 |
| TARA_G.0001117_METAB.16S | G-0001117 | Pacific | Air | 20170513, 11:06 | 4.28 |
| TARA_G.0001118_METAB.16S | G-0001118 | Pacific | Air | 20170504, 11:37 | 20.50 |
| TARA_G.0001121_METAB.16S | G-0001121 | Pacific | Air | 20170504, 10:55 | 20.10 |
| TARA_G.0001123_METAB.16S | G-0001123 | Pacific | Air | 20170504, 11:35 | 19.70 |
| TARA_G.0001143_METAB.16S | G-0001143 | Pacific | Air | 20170506, 12:08 | 19.00 |
| TARA_G.0001150_METAB.16S | G-0001150 | Pacific | Air | 20170506, 12:08 | 18.10 |
| TARA_G.0001161_METAB.16S | G-0001161 | Pacific | Air | 20170507, 11:00 | 17.20 |
| TARA_G.0001162_METAB.16S | G-0001162 | Pacific | Air | 20170507, 11:14 | 16.40 |
| TARA_G.0001189_METAB.16S | G-0001189 | Pacific | Air | 20170507, 12:15 | 15.30 |
| TARA_G.0001206_METAB.16S | G-0001206 | Pacific | Air | 20170509, 11:59 | 11.48 |
| TARA_G.0001211_METAB.16S | G-0001211 | Pacific | Air | 20170510, 12:02 | 10.35 |
| TARA_G.0001230_METAB.16S | G-0001230 | Pacific | Air | 20170511, 11:47 | 7.99 |
| TARA_G.0001279_METAB.16S | G-0001279 | Pacific | Air | 20170512, 12:00 | 5.23 |
| TARA_G.0001280_METAB.16S | G-0001280 | Pacific | Air | 20170513, 12:27 | 4.68 |
| TARA_G.0001285_METAB.16S | G-0001285 | Pacific | Air | 20170514, 11:26 | 3.98 |
| TARA_G.0001306_METAB.16S | G-0001306 | Pacific | Air | 20170514, 12:16 | 3.53 |
| TARA_G.0001307_METAB.16S | G-0001307 | Pacific | Air | 20170515, 11:45 | 3.22 |
| TARA_G.0001314_METAB.16S | G-0001314 | Pacific | Air | 20170515, 11:56 | 2.83 |
| TARA_G.0001325_METAB.16S | G-0001325 | Pacific | Air | 20170516, 11:28 | 1.79 |

| | | | | | |
|---|---|---|---|---|---|
| TARA_G.0001356_METAB.16S | G-0001356 | Pacific | Air | 20170517, 12:14 | 1.28 |
| TARA_G.0001358_METAB.16S | G-0001358 | Pacific | Air | 20170518, 11:52 | 0.40 |
| TARA_G.0001447_METAB.16S | G-0001447 | Pacific | Air | 20170525, 10:56 | -6.40 |
| TARA_G.0001448_METAB.16S | G-0001448 | Pacific | Air | 20170526, 12:08 | -7.30 |
| TARA_G.0001471_METAB.16S | G-0001471 | Pacific | Air | 20170527, 10:35 | -9.80 |
| TARA_G.0001511_METAB.16S | G-0001511 | Pacific | Air | 20170528, 11:46 | -13.52 |
| TARA_G.0001528_METAB.16S | G-0001528 | Pacific | Air | 20170529, 11:51 | -17.18 |
| TARA_OA-0000762_METAB.16S | OA-0000762 | Pacific | Water | 20170430 | 24.70 |
| TARA_OA-0000870_METAB.16S | OA-0000870 | Pacific | Water | 20170511 | 7.99 |
| TARA_OA-0000945_METAB.16S | OA-0000945 | Pacific | Water | 20170517 | 0.86 |
| TARA_OA-0001019_METAB.16S | OA-0001019 | Pacific | Water | 20170524 | -5.03 |
| TARA_OA-0000891_METAB.16S | OA-0000891 | Pacific | Water | 20170512 | 5.23 |
| TARA_OA-0000962_METAB.16S | OA-0000962 | Pacific | Water | 20170518 | 0.40 |
| TARA_OA-0001033_METAB.16S | OA-0001033 | Pacific | Water | 20170528 | -13.52 |
| TARA_OA-0000780_METAB.16S | OA-0000780 | Pacific | Water | 20170504 | 20.50 |
| TARA_OA-0002975_METAB.16S | OA-0002975 | Pacific | Water | 20170509 | 11.48 |
| TARA_OA-0000968_METAB.16S | OA-0000968 | Pacific | Water | 20170519 | -0.99 |
| TARA_OA-0001045_METAB.16S | OA-0001045 | Pacific | Water | 20170526 | -7.30 |
| TARA_OA-0000793_METAB.16S | OA-0000793 | Pacific | Water | 20170503 | 21.10 |
| TARA_OA-0002985_METAB.16S | OA-0002985 | Pacific | Water | 20170508 | 13.15 |
| TARA_OA-0000983_METAB.16S | OA-0000983 | Pacific | Water | 20170520 | -2.06 |
| TARA_OA-0001069_METAB.16S | OA-0001069 | Pacific | Water | 20170530 | -17.18 |
| TARA_OA-0000807_METAB.16S | OA-0000807 | Pacific | Water | 20170504 | 19.70 |
| TARA_OA-0000897_METAB.16S | OA-0000897 | Pacific | Water | 20170513 | 4.28 |
| TARA_OA-0000646_METAB.16S | OA-0000646 | Pacific | Water | 20170522 | -3.51 |
| TARA_OA-0000813_METAB.16S | OA-0000813 | Pacific | Water | 20170512 | 5.98 |
| TARA_OA-0000914_METAB.16S | OA-0000914 | Pacific | Water | 20170514 | 3.53 |
| TARA_OA-0000653_METAB.16S | OA-0000653 | Pacific | Water | 20170523 | -4.27 |
| TARA_OA-0000840_METAB.16S | OA-0000840 | Pacific | Water | 20170507 | 15.30 |
| TARA_OA-0000919_METAB.16S | OA-0000919 | Pacific | Water | 20170515 | 2.83 |
| TARA_OA-0000989_METAB.16S | OA-0000989 | Pacific | Water | 20170521 | -2.79 |
| TARA_OA-0000864_METAB.16S | OA-0000864 | Pacific | Water | 20170510 | 9.19 |
| TARA_OA-0000931_METAB.16S | OA-0000931 | Pacific | Water | 20170516 | 1.79 |
| TARA_OA-0001013_METAB.16S | OA-0001013 | Pacific | Water | 20170525 | -6.40 |

**Table S2. Bacterial taxa detected in blank filter samples.** The taxonomic Assignation of bacteria detected in the blank air filters.

| Blank Taxon ID | Phylum | Class | Order | Family | Genus | Species |
|---|---|---|---|---|---|---|
| B001 | Proteobacteria | Alphaproteobacteria | Caulobacterales | Caulobacteraceae | Brevundimonas | NA |
| B002 | Proteobacteria | Alphaproteobacteria | Caulobacterales | Caulobacteraceae | NA | NA |
| B003 | Proteobacteria | Alphaproteobacteria | Rhizobiales | Beijerinckiaceae | Bosea | NA |
| B004 | Proteobacteria | Alphaproteobacteria | Rhizobiales | Beijerinckiaceae | Methylobacterium | NA |
| B005 | Proteobacteria | Alphaproteobacteria | Rhizobiales | Beijerinckiaceae | Methylobacterium | radiotolerans |
| B006 | Proteobacteria | Alphaproteobacteria | Rhizobiales | Beijerinckiaceae | Methylobacterium | adhaesivum |
| B007 | Proteobacteria | Alphaproteobacteria | Rhizobiales | Hyphomicrobiaceae | Pedomicrobium | ferrugineum |
| B008 | Proteobacteria | Alphaproteobacteria | Rhizobiales | Rhizobiaceae | Allorhizobium-Neorhizobium-Pararhizobium-Rhizobium | NA |
| B009 | Proteobacteria | Alphaproteobacteria | Rhizobiales | Rhizobiaceae | Mesorhizobium | loti |
| B010 | Proteobacteria | Alphaproteobacteria | Rhizobiales | Rhizobiaceae | NA | NA |
| B011 | Proteobacteria | Alphaproteobacteria | Rhizobiales | Rhizobiaceae | Ochrobactrum | NA |
| B012 | Proteobacteria | Alphaproteobacteria | Rhizobiales | Xanthobacteraceae | Afipia | NA |
| B013 | Proteobacteria | Alphaproteobacteria | Rhizobiales | Xanthobacteraceae | Bradyrhizobium | NA |
| B014 | Proteobacteria | Alphaproteobacteria | Rhizobiales | Xanthobacteraceae | Rhodopseudomonas | boonkerdii |
| B015 | Proteobacteria | Alphaproteobacteria | Rhodobacterales | Rhodobacteraceae | Paracoccus | NA |
| B016 | Proteobacteria | Alphaproteobacteria | Sphingomonadales | Sphingomonadaceae | Blastomonas | NA |
| B017 | Proteobacteria | Alphaproteobacteria | Sphingomonadales | Sphingomonadaceae | Erythrobacter | NA |
| B018 | Proteobacteria | Alphaproteobacteria | Sphingomonadales | Sphingomonadaceae | Sphingobium | NA |
| B019 | Proteobacteria | Alphaproteobacteria | Sphingomonadales | Sphingomonadaceae | Sphingobium | amiense |
| B020 | Proteobacteria | Alphaproteobacteria | Sphingomonadales | Sphingomonadaceae | Sphingomonas | NA |
| B021 | Proteobacteria | Deltaproteobacteria | Myxococcales | Polyangiaceae | Pajaroellobacter | NA |
| B022 | Proteobacteria | Gammaproteobacteria | Alteromonadales | Shewanellaceae | Shewanella | NA |
| B023 | Proteobacteria | Gammaproteobacteria | Betaproteobacteriales | Burkholderiaceae | Alicycliphilus | NA |
| B024 | Proteobacteria | Gammaproteobacteria | Betaproteobacteriales | Burkholderiaceae | Aquabacterium | NA |
| B025 | Proteobacteria | Gammaproteobacteria | Betaproteobacteriales | Burkholderiaceae | Aquabacterium | citratiphilum |
| B026 | Proteobacteria | Gammaproteobacteria | Betaproteobacteriales | Burkholderiaceae | Comamonas | denitrificans |
| B027 | Proteobacteria | Gammaproteobacteria | Betaproteobacteriales | Burkholderiaceae | Comamonas | NA |
| B028 | Proteobacteria | Gammaproteobacteria | Betaproteobacteriales | Burkholderiaceae | Curvibacter | NA |
| B029 | Proteobacteria | Gammaproteobacteria | Betaproteobacteriales | Burkholderiaceae | Massilia | NA |
| B030 | Proteobacteria | Gammaproteobacteria | Betaproteobacteriales | Burkholderiaceae | NA | NA |
| B031 | Proteobacteria | Gammaproteobacteria | Betaproteobacteriales | Burkholderiaceae | Ralstonia | NA |
| B032 | Proteobacteria | Gammaproteobacteria | Betaproteobacteriales | Burkholderiaceae | Tepidimonas | NA |
| B033 | Proteobacteria | Gammaproteobacteria | Betaproteobacteriales | Methylophilaceae | Methylobacillus | NA |
| B034 | Proteobacteria | Gammaproteobacteria | Betaproteobacteriales | Neisseriaceae | Neisseria | NA |
| B035 | Proteobacteria | Gammaproteobacteria | Enterobacteriales | Enterobacteriaceae | Klebsiella | NA |
| B036 | Proteobacteria | Gammaproteobacteria | Enterobacteriales | Enterobacteriaceae | NA | NA |
| B037 | Proteobacteria | Gammaproteobacteria | Enterobacteriales | Enterobacteriaceae | Salmonella | NA |
| B038 | Proteobacteria | Gammaproteobacteria | Gammaproteobacteria | Unknown_Family | Acidibacter | NA |
| B039 | Proteobacteria | Gammaproteobacteria | Oceanospirillales | Alcanivoracaceae | Alcanivorax | venustensis |
| B040 | Proteobacteria | Gammaproteobacteria | Oceanospirillales | Halomonadaceae | Chromohalobacter | NA |
| B041 | Proteobacteria | Gammaproteobacteria | Oceanospirillales | Halomonadaceae | Halomonas | phoceae |
| B042 | Proteobacteria | Gammaproteobacteria | Oceanospirillales | Halomonadaceae | Halomonas | NA |
| B043 | Proteobacteria | Gammaproteobacteria | Oceanospirillales | Halomonadaceae | Halomonas | aquamarina |
| B044 | Proteobacteria | Gammaproteobacteria | Pseudomonadales | Moraxellaceae | Acinetobacter | NA |
| B045 | Proteobacteria | Gammaproteobacteria | Pseudomonadales | Moraxellaceae | Acinetobacter | baylyi |
| B046 | Proteobacteria | Gammaproteobacteria | Pseudomonadales | Moraxellaceae | Enhydrobacter | aerosaccus |
| B047 | Proteobacteria | Gammaproteobacteria | Pseudomonadales | Pseudomonadaceae | Pseudomonas | NA |
| B048 | Proteobacteria | Gammaproteobacteria | Salinisphaerales | Salinisphaeraceae | Salinisphaera | japonica |
| B049 | Proteobacteria | Gammaproteobacteria | Salinisphaerales | Salinisphaeraceae | Salinisphaera | NA |
| B050 | Proteobacteria | Gammaproteobacteria | Vibrionales | Vibrionaceae | Vibrio | NA |
| B051 | Proteobacteria | Gammaproteobacteria | Xanthomonadales | Xanthomonadaceae | Stenotrophomonas | NA |
| B052 | Acidobacteria | Acidobacteriia | Solibacterales | Solibacteraceae (Subgroup_3) | Bryobacter | NA |
| B053 | Actinobacteria | Actinobacteria | Corynebacteriales | Corynebacteriaceae | Corynebacterium_1 | appendicis |
| B054 | Actinobacteria | Actinobacteria | Corynebacteriales | Corynebacteriaceae | Corynebacterium_1 | NA |
| B055 | Actinobacteria | Actinobacteria | Corynebacteriales | Corynebacteriaceae | Lawsonella | NA |
| B056 | Actinobacteria | Actinobacteria | Corynebacteriales | Dietziaceae | Dietzia | NA |
| B057 | Actinobacteria | Actinobacteria | Corynebacteriales | Nocardiaceae | Gordonia | NA |
| B058 | Actinobacteria | Actinobacteria | Micrococcales | Dermacoccaceae | Dermacoccus | NA |
| B059 | Actinobacteria | Actinobacteria | Micrococcales | Micrococcaceae | Kocuria | NA |
| B060 | Actinobacteria | Actinobacteria | Micrococcales | Micrococcaceae | Micrococcus | NA |
| B061 | Actinobacteria | Actinobacteria | Micrococcales | Promicromonosporaceae | Cellulosimicrobium | NA |
| B062 | Actinobacteria | Actinobacteria | Propionibacteriales | Propionibacteriaceae | Cutibacterium | NA |
| B063 | Actinobacteria | Actinobacteria | Propionibacteriales | Propionibacteriaceae | Cutibacterium | acnes |

| | | | | | | |
|---|---|---|---|---|---|---|
| B064 | Actinobacteria | Actinobacteria | Propionibacteriales | Propionibacteriaceae | Cutibacterium | granulosum |
| B065 | Actinobacteria | Actinobacteria | Pseudonocardiales | Pseudonocardiaceae | Saccharopolyspora | NA |
| B066 | Firmicutes | Bacilli | Bacillales | Bacillaceae | Bacillus | rigiliprofundi |
| B067 | Firmicutes | Bacilli | Bacillales | Bacillaceae | Bacillus | NA |
| B068 | Firmicutes | Bacilli | Bacillales | Bacillaceae | Bacillus | cihuensis |
| B069 | Firmicutes | Bacilli | Bacillales | Bacillaceae | NA | NA |
| B070 | Firmicutes | Bacilli | Bacillales | Family_XI | Gemella | NA |
| B071 | Firmicutes | Bacilli | Bacillales | Listeriaceae | Listeria | NA |
| B072 | Firmicutes | Bacilli | Bacillales | Staphylococcaceae | Salinicoccus | NA |
| B073 | Firmicutes | Bacilli | Bacillales | Staphylococcaceae | Staphylococcus | NA |
| B074 | Firmicutes | Bacilli | Lactobacillales | Enterococcaceae | Enterococcus | NA |
| B075 | Firmicutes | Bacilli | Lactobacillales | Enterococcaceae | Enterococcus | cecorum |
| B076 | Firmicutes | Bacilli | Lactobacillales | Lactobacillaceae | Lactobacillus | NA |
| B077 | Firmicutes | Bacilli | Lactobacillales | Lactobacillaceae | Lactobacillus | iners |
| B078 | Firmicutes | Bacilli | Lactobacillales | Streptococcaceae | Streptococcus | NA |
| B079 | Bacteroidetes | Bacteroidia | Bacteroidales | Prevotellaceae | Prevotella_7 | NA |
| B080 | Bacteroidetes | Bacteroidia | Chitinophagales | Chitinophagaceae | Sediminibacterium | NA |
| B081 | Bacteroidetes | Bacteroidia | Flavobacteriales | Flavobacteriaceae | Marixanthomonas | NA |
| B082 | Bacteroidetes | Bacteroidia | Flavobacteriales | Weeksellaceae | Chryseobacterium | NA |
| B083 | Bacteroidetes | Bacteroidia | Flavobacteriales | Weeksellaceae | Cloacibacterium | NA |
| B084 | Cyanobacteria | Oxyphotobacteria | Phormidesmiales | Nodosilineaceae | Nodosilinea_PCC-7104 | NA |
| B085 | Chloroflexi | Ktedonobacteria | Ktedonobacterales | Ktedonobacteraceae | JG30a-KF-32 | NA |

**Table S3. Cyanobacteria and Total bacteria concentrations in the Atlantic and Pacific transects.** The calculated concentrations of Cyanobacteria and Total bacteria, obtained from qPCR calibration curves, using DNA extracted from a known number of cells of *Prochlorococcus marinus*, and *Sulfitobacter* D7 cultures, respectively.

| **qPCR Assay** | **Atlantic (Avg. Cell $m^{-3}$)** | **Pacific (Avg. Cell $m^{-3}$)** |
|---|---|---|
| Total bacteria (16S) | 4064.1 ± 6348.1 | 3036.2 ± 1678.9 |
| Cyanobacteria (16S) | 32.5 ± 13.1 | 335.8 ± 129.3 |
| Ratio | 0.0080 ± 0.0129 | 0.1106 ± 0.0745 |

**Table S4. Frequency of appearance of Atmospheric bacteria.** The coverage percentile of bacterial taxa > 5% found only in atmospheric samples.

| Taxa | Atmospheric coverage (%) |
|---|---|
| Proteobacteria_Alphaproteobacteria_Rhodobacterales_Rhodobacteraceae_Paracoccus_NA | 94.2 |
| Proteobacteria_Alphaproteobacteria_Caulobacterales_Caulobacteraceae_NA_NA | 91.3 |
| Proteobacteria_Alphaproteobacteria_Rhizobiales_Rhizobiaceae_Mesorhizobium_NA | 79.7 |
| Proteobacteria_Gammaproteobacteria_Gammaproteobacteria_Acidibacter_NA | 76.8 |
| Proteobacteria_Alphaproteobacteria_Caulobacterales_Caulobacteraceae_Brevundimonas_NA | 76.8 |
| Proteobacteria_Deltaproteobacteria_Myxococcales_Polyangiaceae_Pajaroellobacter_NA | 63.8 |
| Proteobacteria_Alphaproteobacteria_Rhizobiales_Rhizobiaceae_Allorhizobium-Neorhizobium-Pararhizobium-Rhizobium_NA | 62.3 |
| Proteobacteria_Gammaproteobacteria_Xanthomonadales_Xanthomonadaceae_Stenotrophomonas_NA | 60.9 |
| Proteobacteria_Gammaproteobacteria_Betaproteobacteriales_Burkholderiaceae_Massilia_NA | 59.4 |
| Proteobacteria_Deltaproteobacteria_Myxococcales_Polyangiaceae_Minicystis_NA | 58.0 |
| Proteobacteria_Gammaproteobacteria_Enterobacteriales_Enterobacteriaceae_Enterobacter_NA | 52.2 |
| Proteobacteria_Gammaproteobacteria_Betaproteobacteriales_Methylophilaceae_Methylobacillus_NA | 49.3 |
| Proteobacteria_Gammaproteobacteria_Betaproteobacteriales_Burkholderiaceae_Pelomonas_NA | 47.8 |
| Proteobacteria_Gammaproteobacteria_Pseudomonadales_Moraxellaceae_Enhydrobacter_aerosaccus | 43.5 |
| Proteobacteria_Gammaproteobacteria_Betaproteobacteriales_Burkholderiaceae_Tepidimonas_NA | 42.0 |
| Proteobacteria_Alphaproteobacteria_Rhizobiales_Rhizobiaceae_Mesorhizobium_loti | 39.1 |
| Proteobacteria_Alphaproteobacteria_Rhizobiales_Xanthobacteraceae_Afipia_broomeae | 37.7 |
| Proteobacteria_Gammaproteobacteria_Xanthomonadales_Xanthomonadaceae_Silanimonas_NA | 37.7 |
| Proteobacteria_Gammaproteobacteria_Betaproteobacteriales_Burkholderiaceae_Delftia_NA | 34.8 |
| Proteobacteria_Alphaproteobacteria_Rhodobacterales_Rhodobacteraceae_Paracoccus_caeni | 34.8 |
| Proteobacteria_Gammaproteobacteria_Betaproteobacteriales_Methylophilaceae_Methylophilus_NA | 33.3 |
| Proteobacteria_Alphaproteobacteria_Rhizobiales_Beijerinckiaceae_Methylobacterium_radiotolerans | 33.3 |
| Proteobacteria_Gammaproteobacteria_Betaproteobacteriales_Burkholderiaceae_Comamonas_NA | 31.9 |
| Proteobacteria_Alphaproteobacteria_Rhizobiales_Rhizobiaceae_Ochrobactrum_NA | 31.9 |
| Proteobacteria_Alphaproteobacteria_Rhizobiales_Beijerinckiaceae_Bosea_NA | 31.9 |
| Proteobacteria_Alphaproteobacteria_Rhizobiales_Beijerinckiaceae_Methylobacterium_adhaesivum | 30.4 |
| Proteobacteria_Gammaproteobacteria_Pseudomonadales_Moraxellaceae_Acinetobacter_soli | 27.5 |
| Proteobacteria_Gammaproteobacteria_Salinisphaerales_Salinisphaeraceae_Salinisphaera_japonica | 26.1 |
| Proteobacteria_Alphaproteobacteria_Sphingomonadales_Sphingomonadaceae_Blastomonas_NA | 26.1 |
| Proteobacteria_Gammaproteobacteria_Betaproteobacteriales_Burkholderiaceae_Noviherbaspirillum_suwonense | 26.1 |
| Proteobacteria_Gammaproteobacteria_Betaproteobacteriales_Rhodocyclaceae_Methyloversatilis_NA | 26.1 |
| Proteobacteria_Gammaproteobacteria_Betaproteobacteriales_Burkholderiaceae_AAP99_NA | 24.6 |
| Proteobacteria_Gammaproteobacteria_Pseudomonadales_Moraxellaceae_Enhydrobacter_NA | 23.2 |
| Proteobacteria_Gammaproteobacteria_Pseudomonadales_Pseudomonadaceae_Azorhizophilus_NA | 23.2 |
| Proteobacteria_Gammaproteobacteria_Alteromonadales_Alteromonadaceae_Alishewanella_agri | 23.2 |
| Proteobacteria_Alphaproteobacteria_Rhodobacterales_Rhodobacteraceae_Rubellimicrobium_NA | 23.2 |
| Proteobacteria_Alphaproteobacteria_Rhizobiales_Rhizobiaceae_Shinella_NA | 21.7 |
| Proteobacteria_Alphaproteobacteria_Sphingomonadales_Sphingomonadaceae_Sphingobium_yanoikuyae | 21.7 |
| Proteobacteria_Gammaproteobacteria_Betaproteobacteriales_Burkholderiaceae_Curvibacter_NA | 21.7 |
| Proteobacteria_Gammaproteobacteria_Enterobacteriales_Enterobacteriaceae_Atlantibacter_NA | 21.7 |
| Proteobacteria_Gammaproteobacteria_Enterobacteriales_Enterobacteriaceae_Yersinia_NA | 21.7 |
| Proteobacteria_Alphaproteobacteria_Acetobacterales_Acetobacteraceae_Roseomonas_NA | 21.7 |
| Proteobacteria_Gammaproteobacteria_Oceanospirillales_Halomonadaceae_Chromohalobacter_NA | 21.7 |
| Proteobacteria_Alphaproteobacteria_Rhizobiales_Xanthobacteraceae_NA_NA | 20.3 |
| Proteobacteria_Gammaproteobacteria_Betaproteobacteriales_Burkholderiaceae_Variovorax_paradoxus | 20.3 |
| Proteobacteria_Gammaproteobacteria_Pasteurellales_Pasteurellaceae_Actinobacillus_NA | 20.3 |
| Proteobacteria_Gammaproteobacteria_Xanthomonadales_Xanthomonadaceae_SN8_NA | 20.3 |
| Proteobacteria_Alphaproteobacteria_Sphingomonadales_Sphingomonadaceae_Porphyrobacter_NA | 20.3 |
| Proteobacteria_Alphaproteobacteria_Rhizobiales_Beijerinckiaceae_Methylobacterium_komagatae | 20.3 |
| Proteobacteria_Alphaproteobacteria_Rhodobacterales_Rhodobacteraceae_Paracoccus_chinensis | 18.8 |
| Proteobacteria_Alphaproteobacteria_Rhodobacterales_Rhodobacteraceae_Paracoccus_sanguinis | 18.8 |
| Proteobacteria_Gammaproteobacteria_Betaproteobacteriales_Burkholderiaceae_Alicycliphilus_NA | 17.4 |
| Proteobacteria_Gammaproteobacteria_Betaproteobacteriales_Burkholderiaceae_Lautropia_mirabilis | 17.4 |
| Proteobacteria_Gammaproteobacteria_Enterobacteriales_Enterobacteriaceae_Raoultella_NA | 17.4 |
| Proteobacteria_Gammaproteobacteria_Betaproteobacteriales_Burkholderiaceae_Zhizhongheella_caldifontis | 17.4 |
| Proteobacteria_Alphaproteobacteria_Rhizobiales_Rhizobiaceae_Pseudaminobacter_NA | 15.9 |
| Proteobacteria_Alphaproteobacteria_Rhizobiales_Beijerinckiaceae_NA_NA | 15.9 |
| Proteobacteria_Gammaproteobacteria_Betaproteobacteriales_Burkholderiaceae_Acidovorax_temperans | 15.9 |
| Proteobacteria_Alphaproteobacteria_Rhizobiales_Hyphomicrobiaceae_Pedomicrobium_ferrugineum | 14.5 |
| Proteobacteria_Gammaproteobacteria_Alteromonadales_Idiomarinaceae_Idiomarina_fontislapidosi | 14.5 |
| Proteobacteria_Gammaproteobacteria_Alteromonadales_Alteromonadaceae_Alishewanella_NA | 14.5 |
| Proteobacteria_Gammaproteobacteria_Betaproteobacteriales_Burkholderiaceae_Cupriavidus_respiraculi | 14.5 |
| Proteobacteria_Alphaproteobacteria_Micropepsales_Micropepsaceae_NA_NA | 14.5 |

| | |
|---|---|
| Proteobacteria_Gammaproteobacteria_Pasteurellales_Pasteurellaceae_Aggregatibacter_segnis | 13.0 |
| Proteobacteria_Alphaproteobacteria_Rhizobiales_Rhizobiaceae_Aminobacter_NA | 13.0 |
| Proteobacteria_Gammaproteobacteria_Betaproteobacteriales_Burkholderiaceae_Variovorax_NA | 13.0 |
| Proteobacteria_Alphaproteobacteria_Sphingomonadales_Sphingomonadaceae_Novosphingobium_aromaticivorans | 13.0 |
| Proteobacteria_Alphaproteobacteria_Rhizobiales_Rhizobiaceae_Aurantimonas_NA | 13.0 |
| Proteobacteria_Alphaproteobacteria_Rhizobiales_Beijerinckiaceae_Microvirga_NA | 13.0 |
| Proteobacteria_Gammaproteobacteria_Betaproteobacteriales_Rhodocyclaceae_Dechlorosoma_NA | 11.6 |
| Proteobacteria_Gammaproteobacteria_Enterobacteriales_Enterobacteriaceae_Lelliottia_NA | 11.6 |
| Proteobacteria_Alphaproteobacteria_Sphingomonadales_Sphingomonadaceae_Sphingopyxis_baekryungensis | 11.6 |
| Proteobacteria_Gammaproteobacteria_Vibrionales_Vibrionaceae_Photobacterium_damselae | 11.6 |
| Proteobacteria_Gammaproteobacteria_Pseudomonadales_Pseudomonadaceae_Azomonas_NA | 11.6 |
| Proteobacteria_Gammaproteobacteria_Betaproteobacteriales_Rhodocyclaceae_Dechloromonas_agitata | 11.6 |
| Proteobacteria_Gammaproteobacteria_Xanthomonadales_Xanthomonadaceae_Pseudoxanthomonas_NA | 11.6 |
| Proteobacteria_Gammaproteobacteria_Enterobacteriales_Enterobacteriaceae_Pantoea_NA | 10.1 |
| Proteobacteria_Gammaproteobacteria_Pseudomonadales_Moraxellaceae_Alkanindiges_NA | 10.1 |
| Proteobacteria_Gammaproteobacteria_Pseudomonadales_Pseudomonadaceae_Pseudomonas_alcaligenes | 10.1 |
| Proteobacteria_Gammaproteobacteria_Aeromonadales_Aeromonadaceae_Aeromonas_NA | 10.1 |
| Proteobacteria_Alphaproteobacteria_Azospirillales_Azospirillaceae_Skermanella_NA | 10.1 |
| Proteobacteria_Alphaproteobacteria_Reyranellales_Reyranellaceae_Reyranella_massiliensis | 10.1 |
| Proteobacteria_Gammaproteobacteria_Pseudomonadales_Moraxellaceae_Psychrobacter_marincola | 10.1 |
| Proteobacteria_Gammaproteobacteria_Betaproteobacteriales_Burkholderiaceae_Kinneretia_NA | 8.7 |
| Proteobacteria_Gammaproteobacteria_Betaproteobacteriales_Burkholderiaceae_Burkholderia-Caballeronia-Paraburkholderia_NA | 8.7 |
| Proteobacteria_Gammaproteobacteria_Pasteurellales_Pasteurellaceae_NA_NA | 8.7 |
| Proteobacteria_Alphaproteobacteria_Rhizobiales_Xanthobacteraceae_Rhodopseudomonas_boonkerdii | 8.7 |
| Proteobacteria_Gammaproteobacteria_Betaproteobacteriales_Burkholderiaceae_Hydrogenophaga_caeni | 8.7 |
| Proteobacteria_Gammaproteobacteria_Pasteurellales_Pasteurellaceae_Aggregatibacter_aphrophilus | 8.7 |
| Proteobacteria_Gammaproteobacteria_Xanthomonadales_Xanthomonadaceae_Lysobacter_NA | 8.7 |
| Proteobacteria_Gammaproteobacteria_Betaproteobacteriales_Burkholderiaceae_Duganella_NA | 8.7 |
| Proteobacteria_Alphaproteobacteria_Reyranellales_Reyranellaceae_Reyranella_NA | 8.7 |
| Proteobacteria_Gammaproteobacteria_Betaproteobacteriales_Burkholderiaceae_Sphaerotilus_NA | 8.7 |
| Proteobacteria_Alphaproteobacteria_Rhizobiales_Xanthobacteraceae_Xanthobacter_autotrophicus | 8.7 |
| Proteobacteria_Alphaproteobacteria_Rhodobacterales_Rhodobacteraceae_Rhodobacter_NA | 8.7 |
| Proteobacteria_Alphaproteobacteria_Rhodobacterales_Rhodobacteraceae_Roseivivax_NA | 7.2 |
| Proteobacteria_Gammaproteobacteria_Betaproteobacteriales_Burkholderiaceae_Lautropia_NA | 7.2 |
| Proteobacteria_Gammaproteobacteria_Betaproteobacteriales_Burkholderiaceae_Cupriavidus_metallidurans | 7.2 |
| Proteobacteria_Gammaproteobacteria_Betaproteobacteriales_Burkholderiaceae_Hydrogenophaga_intermedia | 7.2 |
| Proteobacteria_Gammaproteobacteria_Xanthomonadales_Xanthomonadaceae_Xanthomonas_NA | 7.2 |
| Proteobacteria_Alphaproteobacteria_Sphingomonadales_Sphingomonadaceae_Qipengyuania_NA | 7.2 |
| Proteobacteria_Alphaproteobacteria_Acetobacterales_Acetobacteraceae_NA_NA | 7.2 |
| Proteobacteria_Gammaproteobacteria_Pseudomonadales_Moraxellaceae_Acinetobacter_schindleri | 7.2 |
| Proteobacteria_Alphaproteobacteria_Rhizobiales_Xanthobacteraceae_Oligotropha_NA | 5.8 |
| Proteobacteria_Alphaproteobacteria_Sphingomonadales_Sphingomonadaceae_Croceicoccus_NA | 5.8 |
| Proteobacteria_Gammaproteobacteria_Betaproteobacteriales_Burkholderiaceae_Polynucleobacter_NA | 5.8 |
| Proteobacteria_Gammaproteobacteria_Pasteurellales_Pasteurellaceae_Aggregatibacter_NA | 5.8 |
| Proteobacteria_Gammaproteobacteria_Enterobacteriales_Enterobacteriaceae_Yokenella_NA | 5.8 |
| Proteobacteria_Gammaproteobacteria_Enterobacteriales_Enterobacteriaceae_Franconibacter_NA | 5.8 |
| Proteobacteria_Gammaproteobacteria_Betaproteobacteriales_Neisseriaceae_Kingella_NA | 5.8 |
| Proteobacteria_Alphaproteobacteria_Sphingomonadales_Sphingomonadaceae_Ellin6055_NA | 5.8 |
| Proteobacteria_Gammaproteobacteria_Betaproteobacteriales_Burkholderiaceae_Limnohabitans_NA | 5.8 |
| Proteobacteria_Alphaproteobacteria_Rhizobiales_Beijerinckiaceae_Methylobacterium_jeotgali | 5.8 |
| Proteobacteria_Gammaproteobacteria_Betaproteobacteriales_Burkholderiaceae_Janthinobacterium_NA | 5.8 |
| Proteobacteria_Gammaproteobacteria_Cellvibrionales_Cellvibrionaceae_Cellvibrio_NA | 5.8 |
| Proteobacteria_Alphaproteobacteria_Sphingomonadales_Sphingomonadaceae_Rhizorhapis_NA | 5.8 |
| Proteobacteria_Gammaproteobacteria_Betaproteobacteriales_Burkholderiaceae_Alcaligenes_NA | 5.8 |
| Proteobacteria_Deltaproteobacteria_Myxococcales_Haliangiaceae_Haliangium_NA | 5.8 |
| Proteobacteria_Gammaproteobacteria_Alteromonadales_Marinobacteraceae_Marinobacter_oulmenensis | 5.8 |
| Proteobacteria_Alphaproteobacteria_Sphingomonadales_Sphingomonadaceae_Sphingomonas_hunanensis | 5.8 |
| Proteobacteria_Gammaproteobacteria_Xanthomonadales_Xanthomonadaceae_Luteimonas_aestuarii | 5.8 |
| Proteobacteria_Gammaproteobacteria_Xanthomonadales_Xanthomonadaceae_Pseudoxanthomonas_mexicana | 5.8 |
| Proteobacteria_Deltaproteobacteria_Myxococcales_Myxococcaceae_Myxococcus_NA | 5.8 |
| Actinobacteria_Actinobacteria_Micrococcales_Micrococcaceae_Micrococcus_NA | 75.4 |
| Actinobacteria_Actinobacteria_Corynebacteriales_Corynebacteriaceae_Lawsonella_NA | 71.0 |
| Actinobacteria_Actinobacteria_Micrococcales_Micrococcaceae_Kocuria_NA | 59.4 |
| Actinobacteria_Actinobacteria_Micrococcales_Brevibacteriaceae_Brevibacterium_NA | 56.5 |
| Actinobacteria_Actinobacteria_Corynebacteriales_Corynebacteriaceae_Turicella_otitidis | 53.6 |
| Actinobacteria_Actinobacteria_Actinomycetales_Actinomycetaceae_Actinomyces_NA | 49.3 |
| Actinobacteria_Actinobacteria_Micrococcales_Promicromonosporaceae_Cellulosimicrobium_NA | 46.4 |
| Actinobacteria_Actinobacteria_Actinomycetales_Actinomycetaceae_Actinomyces_odontolyticus | 36.2 |
| Actinobacteria_Actinobacteria_Corynebacteriales_Dietziaceae_Dietzia_NA | 34.8 |
| Actinobacteria_Actinobacteria_Micrococcales_Dermabacteraceae_Brachybacterium_NA | 30.4 |

| | |
|---|---|
| Actinobacteria_Actinobacteria_Pseudonocardiales_Pseudonocardiaceae_Pseudonocardia_NA | 30.4 |
| Actinobacteria_Actinobacteria_Frankiales_Geodermatophilaceae_Blastococcus_NA | 29.0 |
| Actinobacteria_Rubrobacteria_Rubrobacterales_Rubrobacteriaceae_Rubrobacter_NA | 27.5 |
| Actinobacteria_Actinobacteria_Micrococcales_Dermabacteraceae_Brachybacterium_muris | 27.5 |
| Actinobacteria_Actinobacteria_Micrococcales_Micrococcaceae_Rothia_NA | 26.1 |
| Actinobacteria_Actinobacteria_Micrococcales_Micrococcaceae_Rothia_mucilaginosa | 21.7 |
| Actinobacteria_Actinobacteria_Pseudonocardiales_Pseudonocardiaceae_Saccharopolyspora_NA | 21.7 |
| Actinobacteria_Actinobacteria_Micrococcales_Intrasporangiaceae_Janibacter_NA | 18.8 |
| Actinobacteria_Actinobacteria_Corynebacteriales_Nocardiaceae_Rhodococcus_NA | 18.8 |
| Actinobacteria_Actinobacteria_Micrococcales_Micrococcaceae_NA_NA | 18.8 |
| Actinobacteria_Actinobacteria_Corynebacteriales_Corynebacteriaceae_Turicella_NA | 17.4 |
| Actinobacteria_Actinobacteria_Micrococcales_Dermabacteraceae_Dermabacter_NA | 17.4 |
| Actinobacteria_Actinobacteria_Micrococcales_Micrococcaceae_Rothia_dentocariosa | 15.9 |
| Actinobacteria_Actinobacteria_Propionibacteriales_Nocardioidaceae_Marmoricola_NA | 15.9 |
| Actinobacteria_Actinobacteria_Frankiales_Geodermatophilaceae_Geodermatophilus_NA | 15.9 |
| Actinobacteria_Actinobacteria_Micrococcales_Microbacteriaceae_Agrococcus_NA | 15.9 |
| Actinobacteria_Actinobacteria_Micrococcales_Intrasporangiaceae_Ornithinimicrobium_NA | 13.0 |
| Actinobacteria_Actinobacteria_Micrococcales_Microbacteriaceae_Curtobacterium_NA | 13.0 |
| Actinobacteria_Actinobacteria_Micrococcales_Microbacteriaceae_NA_NA | 13.0 |
| Actinobacteria_Actinobacteria_Streptomycetales_Streptomycetaceae_Streptomyces_NA | 13.0 |
| Actinobacteria_Actinobacteria_Micrococcales_Cellulomonadaceae_Cellulomonas_NA | 11.6 |
| Actinobacteria_Actinobacteria_Micrococcales_Dermacoccaceae_Dermacoccus_NA | 11.6 |
| Actinobacteria_Actinobacteria_Frankiales_Geodermatophilaceae_Modestobacter_NA | 11.6 |
| Actinobacteria_Coriobacteriia_Coriobacteriales_Atopobiaceae_Atopobium_NA | 10.1 |
| Actinobacteria_Thermoleophilia_Solirubrobacterales_67-14_NA_NA | 10.1 |
| Actinobacteria_Actinobacteria_Micrococcales_Intrasporangiaceae_Aquipuribacter_NA | 10.1 |
| Actinobacteria_Actinobacteria_Actinomycetales_Actinomycetaceae_Actinomyces_graevenitzii | 10.1 |
| Actinobacteria_Thermoleophilia_Solirubrobacterales_Solirubrobacteraceae_NA_NA | 8.7 |
| Actinobacteria_Actinobacteria_Micrococcales_Brevibacteriaceae_Brevibacterium_yomogidense | 8.7 |
| Actinobacteria_Actinobacteria_Micrococcales_Micrococcaceae_Arthrobacter_NA | 8.7 |
| Actinobacteria_Actinobacteria_Actinomycetales_Actinomycetaceae_Actinomyces_massiliensis | 7.2 |
| Actinobacteria_Actinobacteria_Micrococcales_Dermacoccaceae_Kytococcus_sedentarius | 7.2 |
| Actinobacteria_Actinobacteria_Micrococcales_Dermacoccaceae_Dermacoccus_nishinomiyaensis | 7.2 |
| Actinobacteria_Actinobacteria_Actinomycetales_Actinomycetaceae_Actinotignum_NA | 7.2 |
| Actinobacteria_Actinobacteria_Corynebacteriales_Tsukamurellaceae_Tsukamurella_NA | 7.2 |
| Actinobacteria_Actinobacteria_Frankiales_Frankiaceae_Jatrophihabitans_NA | 7.2 |
| Actinobacteria_Actinobacteria_Streptosporangiales_Thermomonosporaceae_Actinomadura_NA | 7.2 |
| Actinobacteria_Actinobacteria_Micrococcales_Brevibacteriaceae_Brevibacterium_pityocampae | 7.2 |
| Actinobacteria_Actinobacteria_Micrococcales_Micrococcaceae_Pseudarthrobacter_NA | 7.2 |
| Actinobacteria_Actinobacteria_Micromonosporales_Micromonosporaceae_NA_NA | 7.2 |
| Actinobacteria_Actinobacteria_Micrococcales_Microbacteriaceae_Microbacterium_sediminis | 5.8 |
| Actinobacteria_Actinobacteria_Micrococcales_Micrococcaceae_Glutamicibacter_NA | 5.8 |
| Actinobacteria_Actinobacteria_Bifidobacteriales_Bifidobacteriaceae_Gardnerella_vaginalis | 5.8 |
| Actinobacteria_Thermoleophilia_Gaiellales_Gaiellaceae_Gaiella_NA | 5.8 |
| Actinobacteria_Actinobacteria_Propionibacteriales_Propionibacteriaceae_Friedmanniella_NA | 5.8 |
| Actinobacteria_Thermoleophilia_Solirubrobacterales_Solirubrobacteraceae_Patulibacter_NA | 5.8 |
| Firmicutes_Bacilli_Bacillales_Bacillaceae_Bacillus_NA | 81.2 |
| Firmicutes_Clostridia_Clostridiales_Family_XI_Anaerococcus_NA | 72.5 |
| Firmicutes_Bacilli_Bacillales_Staphylococcaceae_NA_NA | 49.3 |
| Firmicutes_Bacilli_Lactobacillales_Streptococcaceae_Lactococcus_NA | 49.3 |
| Firmicutes_Bacilli_Bacillales_Family_XI_Gemella_NA | 46.4 |
| Firmicutes_Bacilli_Lactobacillales_Carnobacteriaceae_Granulicatella_NA | 46.4 |
| Firmicutes_Bacilli_Lactobacillales_Carnobacteriaceae_Alloiococcus_otitis | 44.9 |
| Firmicutes_Bacilli_Bacillales_Bacillaceae_Marinococcus_NA | 40.6 |
| Firmicutes_Bacilli_Bacillales_Bacillaceae_NA_NA | 40.6 |
| Firmicutes_Bacilli_Bacillales_Alicyclobacillaceae_Tumebacillus_NA | 37.7 |
| Firmicutes_Clostridia_Clostridiales_Family_XI_Peptoniphilus_NA | 36.2 |
| Firmicutes_Clostridia_Clostridiales_Family_XI_Finegoldia_NA | 34.8 |
| Firmicutes_Clostridia_Clostridiales_Clostridiaceae_1_Clostridium_sensu_stricto_11_NA | 33.3 |
| Firmicutes_Clostridia_Clostridiales_Clostridiaceae_1_Clostridium_sensu_stricto_1_saccharobutylicum | 33.3 |
| Firmicutes_Bacilli_Lactobacillales_Carnobacteriaceae_NA_NA | 29.0 |
| Firmicutes_Clostridia_Clostridiales_Family_XI_Ezakiella_NA | 23.2 |
| Firmicutes_Bacilli_Lactobacillales_Enterococcaceae_Melissococcus_NA | 23.2 |
| Firmicutes_Bacilli_Bacillales_Staphylococcaceae_Jeotgalicoccus_NA | 23.2 |
| Firmicutes_Bacilli_Lactobacillales_Carnobacteriaceae_Alkalibacterium_NA | 21.7 |
| Firmicutes_Bacilli_Lactobacillales_Enterococcaceae_NA_NA | 21.7 |
| Firmicutes_Bacilli_Lactobacillales_Aerococcaceae_Eremococcus_NA | 21.7 |
| Firmicutes_Clostridia_Clostridiales_Clostridiaceae_1_Clostridium_sensu_stricto_1_NA | 21.7 |
| Firmicutes_Bacilli_Bacillales_Family_XII_Exiguobacterium_NA | 20.3 |
| Firmicutes_Bacilli_Lactobacillales_Carnobacteriaceae_Granulicatella_elegans | 20.3 |

| | |
|---|---|
| Firmicutes_Clostridia_Clostridiales_Family_XIII_NA_NA | 20.3 |
| Firmicutes_Bacilli_Lactobacillales_Enterococcaceae_Enterococcus_NA | 18.8 |
| Firmicutes_Bacilli_Bacillales_Bacillaceae_Bacillus_rigiliprofundi | 18.8 |
| Firmicutes_Bacilli_Bacillales_Staphylococcaceae_Salinicoccus_NA | 17.4 |
| Firmicutes_Bacilli_Lactobacillales_Carnobacteriaceae_Alloiococcus_NA | 14.5 |
| Firmicutes_Bacilli_Lactobacillales_Carnobacteriaceae_Isobaculum_NA | 14.5 |
| Firmicutes_Bacilli_Lactobacillales_Aerococcaceae_Abiotrophia_defectiva | 13.0 |
| Firmicutes_Bacilli_Lactobacillales_Leuconostocaceae_Leuconostoc_NA | 13.0 |
| Firmicutes_Bacilli_Bacillales_Bacillaceae_Bacillus_cihuensis | 13.0 |
| Firmicutes_Clostridia_Clostridiales_Peptostreptococcaceae_Peptostreptococcus_stomatis | 11.6 |
| Firmicutes_Bacilli_Lactobacillales_Carnobacteriaceae_Carnobacterium_NA | 11.6 |
| Firmicutes_Bacilli_Lactobacillales_Aerococcaceae_Aerococcus_NA | 11.6 |
| Firmicutes_Bacilli_Lactobacillales_Carnobacteriaceae_Atopostipes_NA | 10.1 |
| Firmicutes_Bacilli_Bacillales_Staphylococcaceae_S31_NA | 10.1 |
| Firmicutes_Bacilli_Lactobacillales_Carnobacteriaceae_Desemzia_incerta | 10.1 |
| Firmicutes_Bacilli_Bacillales_Planococcaceae_Lysinibacillus_NA | 10.1 |
| Firmicutes_Clostridia_Clostridiales_Lachnospiraceae_Lachnoanaerobaculum_NA | 10.1 |
| Firmicutes_Bacilli_Lactobacillales_Carnobacteriaceae_Dolosigranulum_pigrum | 10.1 |
| Firmicutes_Clostridia_Clostridiales_Family_XI_Parvimonas_NA | 8.7 |
| Firmicutes_Negativicutes_Selenomonadales_Veillonellaceae_Dialister_propionicifaciens | 8.7 |
| Firmicutes_Bacilli_Bacillales_Planococcaceae_Sporosarcina_NA | 8.7 |
| Firmicutes_Clostridia_Clostridiales_Peptostreptococcaceae_Peptostreptococcus_NA | 8.7 |
| Firmicutes_Clostridia_Clostridiales_Lachnospiraceae_Oribacterium_NA | 8.7 |
| Firmicutes_Clostridia_Clostridiales_Ruminococcaceae_Ruminococcaceae_UCG-014_NA | 8.7 |
| Firmicutes_Clostridia_Clostridiales_Lachnospiraceae_Stomatobaculum_NA | 8.7 |
| Firmicutes_Erysipelotrichia_Erysipelotrichales_Erysipelotrichaceae_Solobacterium_NA | 7.2 |
| Firmicutes_Bacilli_Lactobacillales_Enterococcaceae_Enterococcus_faecalis | 7.2 |
| Firmicutes_Bacilli_Lactobacillales_Aerococcaceae_Facklamia_NA | 7.2 |
| Firmicutes_Clostridia_Clostridiales_Clostridiaceae_1_Clostridium_sensu_stricto_12_NA | 7.2 |
| Firmicutes_Negativicutes_Selenomonadales_Veillonellaceae_Megasphaera_micronuciformis | 7.2 |
| Firmicutes_Clostridia_Clostridiales_Lachnospiraceae_Oribacterium_sinus | 7.2 |
| Firmicutes_Clostridia_Clostridiales_Family_XIII_Mogibacterium_NA | 7.2 |
| Firmicutes_Bacilli_Bacillales_Paenibacillaceae_Brevibacillus_NA | 7.2 |
| Firmicutes_Bacilli_Lactobacillales_Leuconostocaceae_Weissella_NA | 7.2 |
| Firmicutes_Bacilli_Bacillales_Listeriaceae_Brochothrix_thermosphacta | 7.2 |
| Firmicutes_Bacilli_Bacillales_Bacillaceae_Anoxybacillus_NA | 7.2 |
| Firmicutes_Bacilli_Bacillales_Planococcaceae_NA_NA | 5.8 |
| Firmicutes_Bacilli_Bacillales_Listeriaceae_NA_NA | 5.8 |
| Firmicutes_Clostridia_Clostridiales_Lachnospiraceae_Stomatobaculum_longum | 5.8 |
| Bacteroidetes_Bacteroidia_Flavobacteriales_Weeksellaceae_Chryseobacterium_NA | 63.8 |
| Bacteroidetes_Bacteroidia_Bacteroidales_Prevotellaceae_Alloprevotella_NA | 37.7 |
| Bacteroidetes_Bacteroidia_Chitinophagales_Chitinophagaceae_Chitinophaga_NA | 29.0 |
| Bacteroidetes_Bacteroidia_Sphingobacteriales_Sphingobacteriaceae_Sphingobacterium_NA | 24.6 |
| Bacteroidetes_Bacteroidia_Flavobacteriales_Flavobacteriaceae_Zunongwangia_NA | 24.6 |
| Bacteroidetes_Bacteroidia_Flavobacteriales_Flavobacteriaceae_Capnocytophaga_NA | 23.2 |
| Bacteroidetes_Bacteroidia_Cytophagales_Hymenobacteraceae_Hymenobacter_NA | 23.2 |
| Bacteroidetes_Bacteroidia_Flavobacteriales_Weeksellaceae_Cloacibacterium_NA | 21.7 |
| Bacteroidetes_Bacteroidia_Flavobacteriales_Weeksellaceae_Bergeyella_NA | 20.3 |
| Bacteroidetes_Bacteroidia_Cytophagales_Microscillaceae_Siphonobacter_NA | 18.8 |
| Bacteroidetes_Bacteroidia_Flavobacteriales_Flavobacteriaceae_Mesonia_NA | 17.4 |
| Bacteroidetes_Bacteroidia_Chitinophagales_Chitinophagaceae_Taibaiella_NA | 17.4 |
| Bacteroidetes_Bacteroidia_Sphingobacteriales_Sphingobacteriaceae_Pedobacter_NA | 17.4 |
| Bacteroidetes_Bacteroidia_Flavobacteriales_Flavobacteriaceae_Mesonia_mobilis | 17.4 |
| Bacteroidetes_Bacteroidia_Flavobacteriales_Weeksellaceae_Empedobacter_NA | 17.4 |
| Bacteroidetes_Bacteroidia_Flavobacteriales_Weeksellaceae_Elizabethkingia_NA | 14.5 |
| Bacteroidetes_Bacteroidia_Flavobacteriales_Flavobacteriaceae_Salegentibacter_echinorum | 14.5 |
| Bacteroidetes_Bacteroidia_Cytophagales_Spirosomaceae_Fibrella_NA | 13.0 |
| Bacteroidetes_Bacteroidia_Chitinophagales_Chitinophagaceae_Vibrionimonas_NA | 10.1 |
| Bacteroidetes_Bacteroidia_Flavobacteriales_Flavobacteriaceae_Gramella_NA | 10.1 |
| Bacteroidetes_Bacteroidia_Flavobacteriales_Flavobacteriaceae_Zunongwangia_atlantica | 10.1 |
| Bacteroidetes_Bacteroidia_Cytophagales_Hymenobacteraceae_Adhaeribacter_NA | 7.2 |
| Bacteroidetes_Bacteroidia_Chitinophagales_Chitinophagaceae_Flavisolibacter_NA | 7.2 |
| Bacteroidetes_Bacteroidia_Chitinophagales_Chitinophagaceae_Ferruginibacter_NA | 7.2 |
| Bacteroidetes_Bacteroidia_Flavobacteriales_Flavobacteriaceae_Salegentibacter_NA | 7.2 |
| Bacteroidetes_Bacteroidia_Chitinophagales_Chitinophagaceae_Segetibacter_NA | 5.8 |
| Bacteroidetes_Bacteroidia_Chitinophagales_Chitinophagaceae_Flavitalea_NA | 5.8 |
| Bacteroidetes_Bacteroidia_Cytophagales_Spirosomaceae_Pseudarcicella_NA | 5.8 |
| Bacteroidetes_Bacteroidia_Flavobacteriales_Weeksellaceae_Chishuiella_NA | 5.8 |
| Bacteroidetes_Bacteroidia_Cytophagales_Spirosomaceae_Dyadobacter_NA | 5.8 |
| Cyanobacteria_Oxyphotobacteria_Nostocales_Chroococcidiopsaceae_Aliterella_CENA595_NA | 15.9 |

| | |
|---|---|
| Cyanobacteria_Oxyphotobacteria_Phormidesmiales_Nodosilineaceae_Nodosilinea_PCC-7104_NA | 14.5 |
| Cyanobacteria_Oxyphotobacteria_Nostocales_Chroococcidiopsaceae_NA_NA | 10.1 |
| Verrucomicrobia_Verrucomicrobiae_Chthoniobacterales_Chthoniobacteraceae_Chthoniobacter_NA | 8.7 |
| Planctomycetes_Planctomycetacia_Isosphaerales_Isosphaeraceae_NA_NA | 7.2 |
| Planctomycetes_Planctomycetacia_Isosphaerales_Isosphaeraceae_Singulisphaera_NA | 5.8 |
| Deinococcus-Thermus_Deinococci_Deinococcales_Trueperaceae_Truepera_NA | 5.8 |
| Deinococcus-Thermus_Deinococci_Thermales_Thermaceae_Thermus_NA | 8.7 |
| Fusobacteria_Fusobacteriia_Fusobacteriales_Fusobacteriaceae_Fusobacterium_NA | 42.0 |
| Fusobacteria_Fusobacteriia_Fusobacteriales_Leptotrichiaceae_Leptotrichia_NA | 26.1 |
| Fusobacteria_Fusobacteriia_Fusobacteriales_Leptotrichiaceae_Leptotrichia_hongkongensis | 8.7 |
| Acidobacteria_Acidobacteriia_Solibacterales_Solibacteraceae_(Subgroup_3)_Bryobacter_NA | 21.7 |
| Chloroflexi_Chloroflexia_Thermomicrobiales_JG30-KF-CM45_NA_NA | 20.3 |
| Chloroflexi_Ktedonobacteria_Ktedonobacterales_Ktedonobacteraceae_NA_NA | 7.2 |

**Table S5. Frequency of appearance of bacteria detected in both Atmospheric and Oceanic samples.** The coverage percentile of different bacteria > 5% found in atmospheric, and oceanic samples.

| Taxa | Atmospheric coverage (%) | Marine coverage (%) |
|---|---|---|
| Proteobacteria_Alphaproteobacteria_SAR11_clade_Clade_I_Clade_Ia_NA | 37.7 | 100.0 |
| Proteobacteria_Alphaproteobacteria_SAR11_clade_Clade_I_Clade_Ib_NA | 24.6 | 100.0 |
| Proteobacteria_Alphaproteobacteria_Rhodospirillales_AEGEAN-169_marine_group_NA_NA | 17.4 | 100.0 |
| Proteobacteria_Alphaproteobacteria_SAR11_clade_Clade_II_NA_NA | 13.0 | 100.0 |
| Proteobacteria_Alphaproteobacteria_Puniceispirillales_SAR116_clade_NA_NA | 29.0 | 100.0 |
| Proteobacteria_Alphaproteobacteria_Rhodobacterales_Rhodobacteraceae_NA_NA | 43.5 | 100.0 |
| Proteobacteria_Alphaproteobacteria_Rickettsiales_S25-593_NA_NA | 8.7 | 100.0 |
| Proteobacteria_Gammaproteobacteria_Alteromonadales_Alteromonadaceae_Alteromonas_NA | 42.0 | 100.0 |
| Proteobacteria_Gammaproteobacteria_Cellvibrionales_Porticoccaceae_SAR92_clade_NA | 17.4 | 100.0 |
| Proteobacteria_Gammaproteobacteria_Cellvibrionales_Halieaceae_OM60(NOR5)_clade_NA | 4.3 | 100.0 |
| Proteobacteria_Alphaproteobacteria_SAR11_clade_Clade_III_NA_NA | 1.4 | 100.0 |
| Proteobacteria_Alphaproteobacteria_Parvibaculales_Parvibaculaceae_NA_NA | 2.9 | 100.0 |
| Proteobacteria_Alphaproteobacteria_SAR11_clade_Clade_IV_NA_NA | 17.4 | 98.1 |
| Proteobacteria_Alphaproteobacteria_Rhodobacterales_Rhodobacteraceae_Roseovarius_NA | 1.4 | 98.1 |
| Proteobacteria_Gammaproteobacteria_Thiotrichales_Thiotrichaceae_NA_NA | 1.4 | 98.1 |
| Proteobacteria_Deltaproteobacteria_Bdellovibrionales_Bdellovibrionaceae_OM27_clade_NA | 46.4 | 98.1 |
| Proteobacteria_Alphaproteobacteria_Parvibaculales_PS1_clade_NA_NA | 18.8 | 98.1 |
| Proteobacteria_Gammaproteobacteria_Betaproteobacteriales_Burkholderiaceae_MWH-UniP1_aquatic_group_NA | 2.9 | 98.1 |
| Proteobacteria_Gammaproteobacteria_Coxiellales_Coxiellaceae_Coxiella_NA | 13.0 | 98.1 |
| Proteobacteria_Gammaproteobacteria_Oceanospirillales_Pseudohongiellaceae_Pseudohongiella_NA | 1.4 | 98.1 |
| Proteobacteria_Alphaproteobacteria_Thalassobaculales_Nisaeaceae_OM75_clade_NA | 4.3 | 96.2 |
| Proteobacteria_Alphaproteobacteria_Sphingomonadales_Sphingomonadaceae_Erythrobacter_NA | 37.7 | 94.3 |
| Proteobacteria_Deltaproteobacteria_Bdellovibrionales_Bacteriovoracaceae_Peredibacter_NA | 1.4 | 94.3 |
| Proteobacteria_Deltaproteobacteria_Myxococcales_P3OB-42_NA_NA | 13.0 | 92.5 |
| Proteobacteria_Alphaproteobacteria_Puniceispirillales_SAR116_clade_Candidatus_Puniceispirillum_NA | 1.4 | 90.6 |
| Proteobacteria_Deltaproteobacteria_Bdellovibrionales_Bacteriovoracaceae_NA_NA | 34.8 | 90.6 |
| Proteobacteria_Gammaproteobacteria_Ectothiorhodospirales_Ectothiorhodospiraceae_NA_NA | 4.3 | 88.7 |
| Proteobacteria_Alphaproteobacteria_SAR11_clade_Clade_I_NA_NA | 1.4 | 86.8 |
| Proteobacteria_Alphaproteobacteria_Micavibrionales_Micavibrionaceae_NA_NA | 2.9 | 84.9 |
| Proteobacteria_Gammaproteobacteria_Oceanospirillales_Litoricolaceae_Litoricola_NA | 1.4 | 81.1 |
| Proteobacteria_Gammaproteobacteria_Alteromonadales_Marinobacteraceae_Marinobacter_NA | 27.5 | 81.1 |
| Proteobacteria_Alphaproteobacteria_Parvibaculales_OCS116_clade_NA_NA | 5.8 | 79.2 |
| Proteobacteria_Gammaproteobacteria_Oceanospirillales_Halomonadaceae_Halomonas_NA | 62.3 | 79.2 |
| Proteobacteria_Alphaproteobacteria_Rhodospirillales_Magnetospiraceae_NA_NA | 2.9 | 77.4 |
| Proteobacteria_Alphaproteobacteria_Rhodobacterales_Rhodobacteraceae_Ascidiaceihabitans_NA | 2.9 | 73.6 |
| Proteobacteria_Alphaproteobacteria_Rickettsiales_Midichloriaceae_MD3-55_NA | 7.2 | 69.8 |
| Proteobacteria_Gammaproteobacteria_Francisellales_Francisellaceae_NA_NA | 4.3 | 66.0 |
| Proteobacteria_Gammaproteobacteria_Piscirickettsiales_Piscirickettsiaceae_Candidatus_Endoecteinascidia_NA | 8.7 | 64.2 |
| Proteobacteria_Gammaproteobacteria_Vibrionales_Vibrionaceae_Vibrio_NA | 20.3 | 62.3 |
| Proteobacteria_Gammaproteobacteria_Pseudomonadales_Pseudomonadaceae_Pseudomonas_NA | 98.6 | 62.3 |
| Proteobacteria_Gammaproteobacteria_Alteromonadales_Pseudoalteromonadaceae_Pseudoalteromonas_NA | 33.3 | 60.4 |
| Proteobacteria_Deltaproteobacteria_Oligoflexales_Oligoflexaceae_NA_NA | 2.9 | 54.7 |
| Proteobacteria_Gammaproteobacteria_Alteromonadales_Idiomarinaceae_Idiomarina_NA | 50.7 | 52.8 |
| Proteobacteria_Alphaproteobacteria_Rhizobiales_Beijerinckiaceae_Methylobacterium_NA | 81.2 | 50.9 |
| Proteobacteria_Gammaproteobacteria_Alteromonadales_Alteromonadaceae_Alteromonas_australica | 1.4 | 49.1 |
| Proteobacteria_Gammaproteobacteria_Alteromonadales_Alteromonadaceae_Alteromonas_genovensis | 1.4 | 47.2 |
| Proteobacteria_Gammaproteobacteria_Oceanospirillales_Saccharospirillaceae_Oleibacter_NA | 2.9 | 47.2 |
| Proteobacteria_Gammaproteobacteria_Salinisphaerales_Salinisphaeraceae_Salinisphaera_NA | 24.6 | 47.2 |
| Proteobacteria_Gammaproteobacteria_Alteromonadales_Alteromonadaceae_Alteromonas_mediterranea | 1.4 | 45.3 |
| Proteobacteria_Gammaproteobacteria_Legionellales_Legionellaceae_NA_NA | 2.9 | 45.3 |
| Proteobacteria_Alphaproteobacteria_Rhodobacterales_Rhodobacteraceae_Ruegeria_NA | 1.4 | 43.4 |
| Proteobacteria_Deltaproteobacteria_Myxococcales_Nannocystaceae_NA_NA | 1.4 | 43.4 |
| Proteobacteria_Alphaproteobacteria_Rhodobacterales_Rhodobacteraceae_Sulfitobacter_pontiacus | 2.9 | 41.5 |
| Proteobacteria_Alphaproteobacteria_Caulobacterales_Hyphomonadaceae_NA_NA | 2.9 | 41.5 |
| Proteobacteria_Gammaproteobacteria_Oceanospirillales_Alcanivoracaceae_Alcanivorax_borkumensis | 2.9 | 39.6 |
| Proteobacteria_Gammaproteobacteria_Alteromonadales_Shewanellaceae_Shewanella_NA | 37.7 | 39.6 |
| Proteobacteria_Gammaproteobacteria_Alteromonadales_Alteromonadaceae_NA_NA | 1.4 | 35.8 |
| Proteobacteria_Gammaproteobacteria_Vibrionales_Vibrionaceae_NA_NA | 5.8 | 35.8 |
| Proteobacteria_Alphaproteobacteria_Rhodobacterales_Rhodobacteraceae_Roseobacter_NA | 2.9 | 35.8 |
| Proteobacteria_Gammaproteobacteria_Cellvibrionales_Spongiibacteraceae_BD1-7_clade_NA | 13.0 | 34.0 |
| Proteobacteria_Gammaproteobacteria_Pseudomonadales_Moraxellaceae_Psychrobacter_NA | 27.5 | 34.0 |
| Proteobacteria_Gammaproteobacteria_Oceanospirillales_Halomonadaceae_Halomonas_phoceae | 2.9 | 34.0 |

| | | |
|---|---|---|
| Proteobacteria_Alphaproteobacteria_Rickettsiales_Midichloriaceae_NA_NA | 2.9 | 28.3 |
| Proteobacteria_Gammaproteobacteria_Oceanospirillales_Halomonadaceae_Cobetia_NA | 7.2 | 28.3 |
| Proteobacteria_Gammaproteobacteria_Betaproteobacteriales_Burkholderiaceae_Caenimonas_NA | 1.4 | 26.4 |
| Proteobacteria_Gammaproteobacteria_Betaproteobacteriales_Burkholderiaceae_NA_NA | 73.9 | 26.4 |
| Proteobacteria_Gammaproteobacteria_Oceanospirillales_Halomonadaceae_Halomonas_aquamarina | 5.8 | 24.5 |
| Proteobacteria_Gammaproteobacteria_Vibrionales_Vibrionaceae_Photobacterium_NA | 1.4 | 24.5 |
| Proteobacteria_Gammaproteobacteria_Alteromonadales_Marinobacteraceae_Marinobacter_manganoxydans | 2.9 | 22.6 |
| Proteobacteria_Deltaproteobacteria_Bdellovibrionales_Bdellovibrionaceae_Bdellovibrio_NA | 13.0 | 22.6 |
| Proteobacteria_Gammaproteobacteria_Betaproteobacteriales_Burkholderiaceae_Aquabacterium_NA | 39.1 | 22.6 |
| Proteobacteria_Gammaproteobacteria_Oceanospirillales_Alcanivoracaceae_Alcanivorax_NA | 10.1 | 22.6 |
| Proteobacteria_Alphaproteobacteria_Sphingomonadales_Sphingomonadaceae_Novosphingobium_NA | 24.6 | 22.6 |
| Proteobacteria_Alphaproteobacteria_Paracaedibacterales_Paracaedibacteraceae_NA_NA | 1.4 | 22.6 |
| Proteobacteria_Alphaproteobacteria_Rhodobacterales_Rhodobacteraceae_Pseudooceanicola_NA | 1.4 | 20.8 |
| Proteobacteria_Gammaproteobacteria_Betaproteobacteriales_Burkholderiaceae_Acidovorax_NA | 8.7 | 20.8 |
| Proteobacteria_Gammaproteobacteria_Alteromonadales_Alteromonadaceae_Aestuariibacter_aggregatus | 1.4 | 18.9 |
| Proteobacteria_Gammaproteobacteria_Enterobacteriales_Enterobacteriaceae_NA_NA | 71.0 | 18.9 |
| Proteobacteria_Gammaproteobacteria_Steroidobacterales_Woeseiaceae_Woeseia_NA | 1.4 | 18.9 |
| Proteobacteria_Alphaproteobacteria_Rhodospirillales_Terasakiellaceae_NA_NA | 1.4 | 18.9 |
| Proteobacteria_Alphaproteobacteria_Rhodobacterales_Rhodobacteraceae_Sulfitobacter_NA | 7.2 | 18.9 |
| Proteobacteria_Gammaproteobacteria_Oceanospirillales_Halomonadaceae_Salinicola_NA | 36.2 | 18.9 |
| Proteobacteria_Gammaproteobacteria_Oceanospirillales_Marinomonadaceae_Marinomonas_NA | 4.3 | 17.0 |
| Proteobacteria_Gammaproteobacteria_Betaproteobacteriales_Nitrosomonadaceae_mle1-7_NA | 1.4 | 17.0 |
| Proteobacteria_Alphaproteobacteria_Rhodobacterales_Rhodobacteraceae_Aliiroseovarius_NA | 4.3 | 17.0 |
| Proteobacteria_Alphaproteobacteria_Sphingomonadales_Sphingomonadaceae_Sphingobium_NA | 24.6 | 17.0 |
| Proteobacteria_Gammaproteobacteria_Vibrionales_Vibrionaceae_Aliivibrio_NA | 2.9 | 15.1 |
| Proteobacteria_Alphaproteobacteria_Rhodobacterales_Rhodobacteraceae_Sulfitobacter_dubius | 34.8 | 15.1 |
| Proteobacteria_Alphaproteobacteria_Caulobacterales_Hyphomonadaceae_Maricaulis_virginensis | 1.4 | 13.2 |
| Proteobacteria_Alphaproteobacteria_Rhodobacterales_Rhodobacteraceae_Marinibacterium_NA | 2.9 | 13.2 |
| Proteobacteria_Gammaproteobacteria_Betaproteobacteriales_Burkholderiaceae_Limnobacter_thiooxidans | 4.3 | 13.2 |
| Proteobacteria_Gammaproteobacteria_Gammaproteobacteria_Incertae_Sedis_Unknown_Family_Marinicella_NA | 2.9 | 11.3 |
| Proteobacteria_Alphaproteobacteria_Rhodobacterales_Rhodobacteraceae_Roseobacter_clade_NAC11-7_lineage_NA | 1.4 | 11.3 |
| Proteobacteria_Alphaproteobacteria_Caulobacterales_Caulobacteraceae_Caulobacter_NA | 21.7 | 11.3 |
| Proteobacteria_Gammaproteobacteria_Legionellales_Legionellaceae_Legionella_NA | 14.5 | 11.3 |
| Proteobacteria_Gammaproteobacteria_Oceanospirillales_SS1-B-06-26_NA_NA | 1.4 | 9.4 |
| Proteobacteria_Gammaproteobacteria_Alteromonadales_Alteromonadaceae_Salinimonas_NA | 1.4 | 9.4 |
| Proteobacteria_Gammaproteobacteria_Vibrionales_Vibrionaceae_Vibrio_caribbeanicus | 5.8 | 9.4 |
| Proteobacteria_Gammaproteobacteria_Betaproteobacteriales_Burkholderiaceae_Achromobacter_NA | 5.8 | 9.4 |
| Proteobacteria_Gammaproteobacteria_Betaproteobacteriales_Burkholderiaceae_Paucibacter_NA | 15.9 | 9.4 |
| Proteobacteria_Alphaproteobacteria_Caulobacterales_Hyphomonadaceae_Henriciella_NA | 1.4 | 7.5 |
| Proteobacteria_Gammaproteobacteria_Pseudomonadales_Moraxellaceae_Acinetobacter_baylyi | 24.6 | 7.5 |
| Proteobacteria_Alphaproteobacteria_Rhodospirillales_Thalassospiraceae_Thalassospira_NA | 2.9 | 7.5 |
| Proteobacteria_Deltaproteobacteria_Myxococcales_BIrii41_NA_NA | 2.9 | 7.5 |
| Proteobacteria_Gammaproteobacteria_Oceanospirillales_Alcanivoracaceae_Alcanivorax_jadensis | 2.9 | 5.7 |
| Proteobacteria_Deltaproteobacteria_Myxococcales_Sandaracinaceae_Sandaracinus_NA | 5.8 | 5.7 |
| Proteobacteria_Gammaproteobacteria_Pseudomonadales_Pseudomonadaceae_NA_NA | 11.6 | 5.7 |
| Proteobacteria_Alphaproteobacteria_Rhizobiales_Rhizobiaceae_NA_NA | 27.5 | 5.7 |
| Proteobacteria_Deltaproteobacteria_Oligoflexales_053A03-B-DI-P58_NA_NA | 7.2 | 5.7 |
| Proteobacteria_Alphaproteobacteria_Rhodobacterales_Rhodobacteraceae_Oceanibulbus_indolifex | 10.1 | 5.7 |
| Proteobacteria_Alphaproteobacteria_Sphingomonadales_Sphingomonadaceae_Altererythrobacter_NA | 17.4 | 5.7 |
| Proteobacteria_Alphaproteobacteria_Sphingomonadales_Sphingomonadaceae_NA_NA | 7.2 | 3.8 |
| Proteobacteria_Alphaproteobacteria_Rhizobiales_Devosiaceae_Devosia_NA | 26.1 | 1.9 |
| Proteobacteria_Gammaproteobacteria_Betaproteobacteriales_Neisseriaceae_NA_NA | 55.1 | 1.9 |
| Actinobacteria_Acidimicrobiia_Actinomarinales_Actinomarinaceae_Candidatus_Actinomarina_NA | 31.9 | 100.0 |
| Actinobacteria_Acidimicrobiia_Microtrichales_Microtrichaceae_Sva0996_marine_group_NA | 14.5 | 54.7 |
| Actinobacteria_Acidimicrobiia_Microtrichales_Ilumatobacteraceae_Ilumatobacter_NA | 10.1 | 5.7 |
| Actinobacteria_Acidimicrobiia_Microtrichales_Ilumatobacteraceae_NA_NA | 7.2 | 3.8 |
| Actinobacteria_Actinobacteria_Propionibacteriales_Nocardioidaceae_Nocardioides_NA | 42.0 | 3.8 |
| Actinobacteria_Actinobacteria_Corynebacteriales_Mycobacteriaceae_Mycobacterium_NA | 21.7 | 1.9 |
| Actinobacteria_Actinobacteria_Micrococcales_Microbacteriaceae_Microbacterium_NA | 50.7 | 1.9 |
| Firmicutes_Bacilli_Bacillales_Paenibacillaceae_Paenibacillus_NA | 29.0 | 1.9 |
| Firmicutes_Clostridia_Clostridiales_Lachnospiraceae_NA_NA | 20.3 | 1.9 |
| Bacteroidetes_Bacteroidia_Flavobacteriales_Flavobacteriaceae_NS4_marine_group_NA | 21.7 | 100.0 |
| Bacteroidetes_Bacteroidia_Flavobacteriales_Flavobacteriaceae_NS5_marine_group_NA | 23.2 | 100.0 |
| Bacteroidetes_Bacteroidia_Flavobacteriales_Flavobacteriaceae_NS2b_marine_group_NA | 15.9 | 100.0 |
| Bacteroidetes_Bacteroidia_Flavobacteriales_NS9_marine_group_NA_NA | 46.4 | 100.0 |
| Bacteroidetes_Bacteroidia_Flavobacteriales_NS7_marine_group_NA_NA | 24.6 | 100.0 |
| Bacteroidetes_Bacteroidia_Flavobacteriales_Cryomorphaceae_NA_NA | 2.9 | 100.0 |
| Bacteroidetes_Bacteroidia_Cytophagales_Cyclobacteriaceae_Marinoscillum_NA | 4.3 | 98.1 |
| Bacteroidetes_Rhodothermia_Balneolales_Balneolaceae_Balneola_NA | 1.4 | 81.1 |

| | | |
|---|---|---|
| Bacteroidetes_Bacteroidia_Flavobacteriales_Flavobacteriaceae_NA_NA | 7.2 | 75.5 |
| Bacteroidetes_Bacteroidia_Flavobacteriales_Flavobacteriaceae_Muricauda_NA | 7.2 | 75.5 |
| Bacteroidetes_Bacteroidia_Flavobacteriales_Flavobacteriaceae_Formosa_NA | 4.3 | 73.6 |
| Bacteroidetes_Bacteroidia_Flavobacteriales_Flavobacteriaceae_Polaribacter_4_NA | 2.9 | 67.9 |
| Bacteroidetes_Bacteroidia_Flavobacteriales_Flavobacteriaceae_Winogradskyella_NA | 2.9 | 67.9 |
| Bacteroidetes_Bacteroidia_Chitinophagales_Saprospiraceae_Aureispira_NA | 5.8 | 60.4 |
| Bacteroidetes_Bacteroidia_Flavobacteriales_Crocinitomicaceae_Fluviicola_NA | 2.9 | 52.8 |
| Bacteroidetes_Bacteroidia_Chitinophagales_Saprospiraceae_NA_NA | 1.4 | 49.1 |
| Bacteroidetes_Bacteroidia_Flavobacteriales_Flavobacteriaceae_Dokdonia_NA | 1.4 | 37.7 |
| Bacteroidetes_Bacteroidia_Cytophagales_Cyclobacteriaceae_NA_NA | 1.4 | 37.7 |
| Bacteroidetes_Bacteroidia_Flavobacteriales_Flavobacteriaceae_Maribacter_NA | 4.3 | 35.8 |
| Bacteroidetes_Bacteroidia_Flavobacteriales_Flavobacteriaceae_Leeuwenhoekiella_blandensis | 4.3 | 34.0 |
| Bacteroidetes_Bacteroidia_Flavobacteriales_Flavobacteriaceae_Tenacibaculum_NA | 7.2 | 32.1 |
| Bacteroidetes_Bacteroidia_Flavobacteriales_Flavobacteriaceae_Leeuwenhoekiella_NA | 1.4 | 32.1 |
| Bacteroidetes_Bacteroidia_Chitinophagales_Saprospiraceae_Saprospira_NA | 10.1 | 30.2 |
| Bacteroidetes_Bacteroidia_Cytophagales_Cyclobacteriaceae_Fabibacter_NA | 11.6 | 24.5 |
| Bacteroidetes_Bacteroidia_Sphingobacteriales_NS11-12_marine_group_NA_NA | 4.3 | 20.8 |
| Bacteroidetes_Bacteroidia_Cytophagales_Cyclobacteriaceae_Algoriphagus_NA | 2.9 | 20.8 |
| Bacteroidetes_Bacteroidia_Flavobacteriales_Flavobacteriaceae_Kordia_NA | 11.6 | 17.0 |
| Bacteroidetes_Bacteroidia_Flavobacteriales_Cryomorphaceae_Owenweeksia_NA | 1.4 | 13.2 |
| Bacteroidetes_Bacteroidia_Flavobacteriales_Flavobacteriaceae_Mesoflavibacter_NA | 1.4 | 9.4 |
| Bacteroidetes_Bacteroidia_Flavobacteriales_Flavobacteriaceae_Muricauda_beolgyonensis | 1.4 | 5.7 |
| Bacteroidetes_Rhodothermia_Rhodothermales_Rhodothermaceae_Rubrivirga_NA | 5.8 | 5.7 |
| Bacteroidetes_Bacteroidia_Flavobacteriales_Flavobacteriaceae_Olleya_NA | 1.4 | 5.7 |
| Bacteroidetes_Bacteroidia_Flavobacteriales_Flavobacteriaceae_Marixanthomonas_NA | 21.7 | 3.8 |
| Bacteroidetes_Bacteroidia_Flavobacteriales_Crocinitomicaceae_NA_NA | 4.3 | 3.8 |
| Bacteroidetes_Bacteroidia_Chitinophagales_Chitinophagaceae_NA_NA | 13.0 | 1.9 |
| Bacteroidetes_Bacteroidia_Cytophagales_Spirosomaceae_Spirosoma_NA | 5.8 | 1.9 |
| Bacteroidetes_Bacteroidia_Flavobacteriales_Flavobacteriaceae_Flavobacterium_NA | 30.4 | 1.9 |
| Bacteroidetes_Bacteroidia_Chitinophagales_Chitinophagaceae_Sediminibacterium_NA | 58.0 | 1.9 |
| Cyanobacteria_Oxyphotobacteria_Synechococcales_Cyanobiaceae_Prochlorococcus_MIT9313_marinus | 40.6 | 100.0 |
| Cyanobacteria_Oxyphotobacteria_Synechococcales_Cyanobiaceae_Synechococcus_CC9902_NA | 7.2 | 100.0 |
| Cyanobacteria_Oxyphotobacteria_Synechococcales_Cyanobiaceae_Prochlorococcus_MIT9313_NA | 44.9 | 92.5 |
| Cyanobacteria_Oxyphotobacteria_Synechococcales_Cyanobiaceae_Synechococcus_MBIC10613_NA | 1.4 | 54.7 |
| Cyanobacteria_Oxyphotobacteria_Synechococcales_Cyanobiaceae_NA_NA | 1.4 | 47.2 |
| Cyanobacteria_Oxyphotobacteria_Nostocales_Microcystaceae_Atelocyanobacterium_(UCYN-A)_thalassa | 1.4 | 11.3 |
| Cyanobacteria_Oxyphotobacteria_Nostocales_Microcystaceae_Crocosphaera_WH_0003_(UCYN-B)_watsonii | 1.4 | 9.4 |
| Cyanobacteria_Oxyphotobacteria_Nostocales_Xenococcaceae_Pleurocapsa_PCC-7319_NA | 4.3 | 1.9 |
| Verrucomicrobia_Verrucomicrobiae_Opitutales_Puniceicoccaceae_Lentimonas_NA | 47.8 | 100.0 |
| Verrucomicrobia_Verrucomicrobiae_Opitutales_Puniceicoccaceae_MB11C04_marine_group_NA | 4.3 | 98.1 |
| Verrucomicrobia_Verrucomicrobiae_Opitutales_Puniceicoccaceae_Coraliomargarita_NA | 17.4 | 90.6 |
| Verrucomicrobia_Verrucomicrobiae_Verrucomicrobiales_DEV007_NA_NA | 8.7 | 62.3 |
| Verrucomicrobia_Verrucomicrobiae_Verrucomicrobiales_Rubritaleaceae_Roseibacillus_NA | 1.4 | 58.5 |
| Verrucomicrobia_Verrucomicrobiae_Opitutales_Puniceicoccaceae_Pelagicoccus_NA | 1.4 | 43.4 |
| Verrucomicrobia_Verrucomicrobiae_Verrucomicrobiales_Rubritaleaceae_Rubritalea_NA | 1.4 | 24.5 |
| Verrucomicrobia_Verrucomicrobiae_Opitutales_Puniceicoccaceae_Cerasicoccus_NA | 1.4 | 17.0 |
| Verrucomicrobia_Verrucomicrobiae_Pedosphaerales_Pedosphaeraceae_NA_NA | 4.3 | 3.8 |
| Planctomycetes_Planctomycetacia_Pirellulales_Pirellulaceae_NA_NA | 10.1 | 90.6 |
| Planctomycetes_Phycisphaerae_Phycisphaerales_Phycisphaeraceae_Urania-1B-19_marine_sediment_group_NA | 34.8 | 86.8 |
| Planctomycetes_Phycisphaerae_Phycisphaerales_Phycisphaeraceae_CL500-3_NA | 42.0 | 83.0 |
| Planctomycetes_Planctomycetacia_Pirellulales_Pirellulaceae_Pirellula_NA | 4.3 | 71.7 |
| Planctomycetes_Phycisphaerae_Phycisphaerales_Phycisphaeraceae_FS140-16B-02_marine_group_NA | 23.2 | 41.5 |
| Planctomycetes_Planctomycetacia_Pirellulales_Pirellulaceae_Rubripirellula_NA | 1.4 | 26.4 |
| Planctomycetes_Phycisphaerae_Phycisphaerales_Phycisphaeraceae_SM1A02_NA | 1.4 | 5.7 |
| Planctomycetes_Planctomycetacia_Pirellulales_Pirellulaceae_Rhodopirellula_NA | 4.3 | 3.8 |
| Kiritimatiellaeota_Kiritimatiellae_Kiritimatiellales_Kiritimatiellaceae_R76-B128_NA | 30.4 | 81.1 |
| Deinococcus-Thermus_Deinococci_Deinococcales_Deinococcaceae_Deinococcus_NA | 42.0 | 1.9 |
| Acidobacteria_Blastocatellia_(Subgroup_4)_Blastocatellales_Blastocatellaceae_Blastocatella_NA | 46.4 | 1.9 |

**Table S6. Frequency of appearance of marine bacteria.** The coverage percentile of bacterial taxa > 5% found only in oceanic samples.

| Taxa | Marine coverage (%) |
|---|---|
| Proteobacteria_Alphaproteobacteria_Caulobacterales_Hyphomonadaceae_Ponticaulis_NA | 49.1 |
| Proteobacteria_Alphaproteobacteria_Caulobacterales_Hyphomonadaceae_Hyphomonas_NA | 47.2 |
| Proteobacteria_Deltaproteobacteria_Bdellovibrionales_Bacteriovoracaceae_Halobacteriovorax_NA | 47.2 |
| Proteobacteria_Alphaproteobacteria_Caulobacterales_Hyphomonadaceae_Oceanicaulis_NA | 45.3 |
| Proteobacteria_Gammaproteobacteria_Nitrosococcales_Methylophagaceae_Methylophaga_nitratireducenticrescens | 43.4 |
| Proteobacteria_Alphaproteobacteria_Rhizobiales_Stappiaceae_Labrenzia_NA | 43.4 |
| Proteobacteria_Alphaproteobacteria_Rickettsiales_Rickettsiaceae_NA_NA | 41.5 |
| Proteobacteria_Alphaproteobacteria_Caulobacterales_Hyphomonadaceae_Maricaulis_maris | 41.5 |
| Proteobacteria_Gammaproteobacteria_Alteromonadales_Pseudoalteromonadaceae_Pseudoalteromonas_spongiae | 41.5 |
| Proteobacteria_Alphaproteobacteria_Rhodobacterales_Rhodobacteraceae_Pseudophaeobacter_NA | 41.5 |
| Proteobacteria_Alphaproteobacteria_Rhodobacterales_Rhodobacteraceae_Loktanella_NA | 39.6 |
| Proteobacteria_Alphaproteobacteria_Caulobacterales_Hyphomonadaceae_Oceanicaulis_alexandrii | 37.7 |
| Proteobacteria_Alphaproteobacteria_Caulobacterales_Hyphomonadaceae_Hirschia_baltica | 34.0 |
| Proteobacteria_Alphaproteobacteria_Rhizobiales_Hyphomicrobiaceae_Filomicrobium_NA | 34.0 |
| Proteobacteria_Deltaproteobacteria_Oligoflexales_Oligoflexaceae_Pseudobacteriovorax_NA | 32.1 |
| Proteobacteria_Alphaproteobacteria_Rhizobiales_Rhizobiales_Incertae_Sedis_Phreatobacter_NA | 32.1 |
| Proteobacteria_Alphaproteobacteria_Rickettsiales_Rickettsiaceae_Candidatus_Megaira_NA | 32.1 |
| Proteobacteria_Gammaproteobacteria_Alteromonadales_Alteromonadaceae_Aestuariibacter_NA | 32.1 |
| Proteobacteria_Gammaproteobacteria_Oceanospirillales_Kangiellaceae_Kangiella_NA | 30.2 |
| Proteobacteria_Gammaproteobacteria_Alteromonadales_Alteromonadaceae_Paraglaciecola_NA | 28.3 |
| Proteobacteria_Alphaproteobacteria_Rhodobacterales_Rhodobacteraceae_Nautella_italica | 28.3 |
| Proteobacteria_Alphaproteobacteria_Rickettsiales_Rickettsiaceae_Occidentia_NA | 26.4 |
| Proteobacteria_Gammaproteobacteria_Salinisphaerales_Solimonadaceae_Oceanococcus_NA | 26.4 |
| Proteobacteria_Alphaproteobacteria_Thalassobaculales_Nisaeaceae_Nisaea_NA | 24.5 |
| Proteobacteria_Gammaproteobacteria_Alteromonadales_Pseudoalteromonadaceae_Pseudoalteromonas_phenolica | 24.5 |
| Proteobacteria_Alphaproteobacteria_Caulobacterales_Hyphomonadaceae_Ponticaulis_koreensis | 24.5 |
| Proteobacteria_Gammaproteobacteria_Tenderiales_Tenderiaceae_Candidatus_Tenderia_NA | 24.5 |
| Proteobacteria_Gammaproteobacteria_Oceanospirillales_Saccharospirillaceae_Thalassolituus_oleivorans | 22.6 |
| Proteobacteria_Gammaproteobacteria_Alteromonadales_Alteromonadaceae_Glaciecola_NA | 22.6 |
| Proteobacteria_Alphaproteobacteria_Rhodovibrionales_Kiloniellaceae_Tistlia_NA | 22.6 |
| Proteobacteria_Gammaproteobacteria_Oceanospirillales_Nitrincolaceae_Neptuniibacter_pectenicola | 22.6 |
| Proteobacteria_Gammaproteobacteria_Betaproteobacteriales_Burkholderiaceae_Aquabacterium_commune | 20.8 |
| Proteobacteria_Gammaproteobacteria_Thiohalorhabdales_Thiohalorhabdaceae_NA_NA | 18.9 |
| Proteobacteria_Gammaproteobacteria_Oceanospirillales_Oleiphilaceae_Oleiphilus_NA | 18.9 |
| Proteobacteria_Alphaproteobacteria_Caulobacterales_Hyphomonadaceae_Litorimonas_NA | 17.0 |
| Proteobacteria_Gammaproteobacteria_Vibrionales_Vibrionaceae_Catenococcus_NA | 17.0 |
| Proteobacteria_Gammaproteobacteria_Betaproteobacteriales_Methylophilaceae_OM43_clade_NA | 17.0 |
| Proteobacteria_Gammaproteobacteria_Thiomicrospirales_Thioglobaceae_SUP05_cluster_NA | 13.2 |
| Proteobacteria_Alphaproteobacteria_Rhodobacterales_Rhodobacteraceae_Planktomarina_NA | 13.2 |
| Proteobacteria_Alphaproteobacteria_Rhodobacterales_Rhodobacteraceae_Amylibacter_NA | 13.2 |
| Proteobacteria_Gammaproteobacteria_Arenicellales_Arenicellaceae_Arenicella_NA | 11.3 |
| Proteobacteria_Gammaproteobacteria_Alteromonadales_Alteromonadaceae_Alteromonas_litorea | 11.3 |
| Proteobacteria_Alphaproteobacteria_Rhodobacterales_Rhodobacteraceae_Marinovum_NA | 11.3 |
| Proteobacteria_Alphaproteobacteria_Rhodobacterales_Rhodobacteraceae_Lentibacter_NA | 11.3 |
| Proteobacteria_Alphaproteobacteria_Holosporales_Holosporaceae_NA_NA | 9.4 |
| Proteobacteria_Gammaproteobacteria_Alteromonadales_Alteromonadaceae_Alteromonas_lipolytica | 9.4 |
| Proteobacteria_Gammaproteobacteria_Alteromonadales_Colwelliaceae_Thalassotalea_NA | 7.5 |
| Bacteroidetes_Bacteroidia_Cytophagales_Flammeovirgaceae_NA_NA | 35.8 |
| Bacteroidetes_Bacteroidia_Flavobacteriales_Flavobacteriaceae_Aurantivirga_NA | 34.0 |
| Bacteroidetes_Bacteroidia_Flavobacteriales_Flavobacteriaceae_Mesonia_algae | 28.3 |
| Bacteroidetes_Bacteroidia_Flavobacteriales_Flavobacteriaceae_Croceibacter_NA | 24.5 |
| Bacteroidetes_Bacteroidia_Flavobacteriales_Flavobacteriaceae_Tenacibaculum_mesophilum | 20.8 |
| Bacteroidetes_Bacteroidia_Flavobacteriales_Flavobacteriaceae_Nonlabens_NA | 18.9 |
| Bacteroidetes_Bacteroidia_Flavobacteriales_Flavobacteriaceae_Gilvibacter_NA | 17.0 |
| Bacteroidetes_Bacteroidia_Flavobacteriales_Flavobacteriaceae_Psychroserpens_NA | 15.1 |
| Bacteroidetes_Bacteroidia_Flavobacteriales_Flavobacteriaceae_Pseudofulvibacter_NA | 11.3 |
| Bacteroidetes_Bacteroidia_Flavobacteriales_Flavobacteriaceae_Aquibacter_NA | 11.3 |
| Bacteroidetes_Bacteroidia_Cytophagales_Cyclobacteriaceae_Ekhidna_NA | 11.3 |
| Bacteroidetes_Bacteroidia_Flavobacteriales_Flavobacteriaceae_Tenacibaculum_geojense | 9.4 |
| Bacteroidetes_Bacteroidia_Flavobacteriales_Cryomorphaceae_NS10_marine_group_NA | 9.4 |
| Bacteroidetes_Bacteroidia_Chitinophagales_Saprospiraceae_Lewinella_NA | 9.4 |
| Bacteroidetes_Bacteroidia_Flavobacteriales_Flavobacteriaceae_Flavicella_NA | 7.5 |
| Bacteroidetes_Bacteroidia_Flavobacteriales_Flavobacteriaceae_Gilvibacter_sediminis | 7.5 |

| | |
|---|---|
| Cyanobacteria_Oxyphotobacteria_Nostocales_Nostocales_Incertae_Sedis_Phormidium_SAG_81.79_NA | 37.7 |
| Cyanobacteria_Oxyphotobacteria_Synechococcales_Cyanobiaceae_Cyanobium_PCC-6307_NA | 34.0 |
| Cyanobacteria_Oxyphotobacteria_Nostocales_Phormidiaceae_Trichodesmium_IMS101_thiebautii | 28.3 |
| Verrucomicrobia_Verrucomicrobiae_Opitutales_Puniceicoccaceae_NA_NA | 28.3 |
| Verrucomicrobia_Verrucomicrobiae_Pedosphaerales_Pedosphaeraceae_SCGC_AAA164-E04_NA | 22.6 |
| Planctomycetes_Planctomycetacia_Pirellulales_Pirellulaceae_Blastopirellula_NA | 39.6 |
| Lentisphaerae_Lentisphaeria_Lentisphaerales_Lentisphaeraceae_Lentisphaera_NA | 22.6 |

**Table S7. Calcification of environmental ontology (ENVO) terms.** The ENVO terms were clustered into five main groups: marine, terrestrial, fresh water, anthropogenic, and unclassified, and their source internationalized resource identifier (IRI).

| Cluster | Term | IRI | Cluster | Term | IRI | Cluster | Term | IRI |
|---|---|---|---|---|---|---|---|---|
| Marine | abyssal plain | **ENVO_00000244 | Terrestrial | humus | **ENVO_01000000 | Fresh water | lake | **ENVO_00000020 |
| Marine | aphotic zone | **ENVO_00000210 | Terrestrial | hypersaline water | **ENVO_00002012 | Fresh water | marsh | **ENVO_00000035 |
| Marine | archipelago | **ENVO_00000220 | Terrestrial | ice sheet | **ENVO_00000132 | Fresh water | meromictic lake | **ENVO_00000199 |
| Marine | atoll | **ENVO_00000166 | Terrestrial | ice shelf | **ENVO_00000380 | Fresh water | pond | **ENVO_00000033 |
| Marine | Back-arc basin | **ENVO_00002277 | Terrestrial | iceberg | **ENVO_00000298 | Fresh water | pond water | **ENVO_00002228 |
| Marine | bathypelagic zone | **ENVO_00000211 | Terrestrial | impact crater | **ENVO_01001071 | Fresh water | reservoir | **ENVO_00000025 |
| Marine | bay | **ENVO_00000032 | Terrestrial | island | **ENVO_00000098 | Fresh water | river | **ENVO_00000022 |
| Marine | beach | **ENVO_00000091 | Terrestrial | karst | **ENVO_00000175 | Fresh water | river bed | **ENVO_00000384 |
| Marine | black smoker | **ENVO_00000218 | Terrestrial | lake bed | **ENVO_00000268 | Fresh water | spring | **ENVO_00000027 |
| Marine | blowhole | **ENVO_00000168 | Terrestrial | lake sediment | **ENVO_00000546 | Fresh water | stream | **ENVO_00000023 |
| Marine | brackish water | **ENVO_00002019 | Terrestrial | lake shore | **ENVO_00000382 | Fresh water | tap water | **ENVO_00003096 |
| Marine | channel | **ENVO_00000395 | Terrestrial | lava | **ENVO_01000231 | Fresh water | underground water | **ENVO_00005792 |
| Marine | coastal upwelling | **ENVO_01000006 | Terrestrial | lava cave | **ENVO_00000322 | Fresh water | water body | **ENVO_00000063 |
| Marine | coastal water | **ENVO_00002150 | Terrestrial | leachate | **ENVO_00002141 | Fresh water | water well | **ENVO_01000002 |
| Marine | coastal water body | **ENVO_02000049 | Terrestrial | limestone | **ENVO_00002053 | Fresh water | watercourse | **ENVO_00000029 |
| Marine | coastal wetland | **ENVO_00000230 | Terrestrial | loam | **ENVO_00002258 | Fresh water | waterfall | **ENVO_00000040 |
| Marine | cold seep | **ENVO_01000263 | Terrestrial | mangrove swamp | **ENVO_00000057 | Fresh water | watershed | **ENVO_00000291 |
| Marine | continental shelf | **ENVO_00000223 | Terrestrial | massif | **ENVO_00000381 | Fresh water | wetland | **ENVO_00000043 |
| Marine | continental slope | **ENVO_00000273 | Terrestrial | meadow | **ENVO_00000108 | Anthropogenic | acid mine drainage | **ENVO_00001997 |
| Marine | coral reef | **ENVO_00000150 | Terrestrial | meadow soil | **ENVO_00005761 | Anthropogenic | activated sludge | **ENVO_00002046 |
| Marine | marine bulk water | **ENVO_01000055 | Terrestrial | mediterranean | **ENVO_01000207 | Anthropogenic | administrative region | **ENVO_00000004 |
| Marine | marine habitat | **ENVO_00000569 | Terrestrial | metal contaminated soil | **ENVO_00003081 | Anthropogenic | agricultural feature | **ENVO_00000077 |
| Marine | marine reef | **ENVO_01000143 | Terrestrial | microbial mat | **ENVO_01000008 | Anthropogenic | agricultural soil | **ENVO_00002259 |
| Marine | marine sediment | **ENVO_00002113 | Terrestrial | moor | **ENVO_00000231 | Anthropogenic | agricultural waste | **ENVO_00002265 |
| Marine | marine snow | **ENVO_01000158 | Terrestrial | moraine | **ENVO_00000177 | Anthropogenic | air filter | **ENVO_00003968 |
| Marine | marine terrace | **ENVO_00000509 | Terrestrial | mound | **ENVO_00000180 | Anthropogenic | anaerobic bioreactor | **ENVO_00002124 |
| Marine | estuary | **ENVO_00000045 | Terrestrial | mountain | **ENVO_00000081 | Anthropogenic | anaerobic digester sludge | **ENVO_00003965 |
| Marine | fjord | **ENVO_00000039 | Terrestrial | mud | **ENVO_01000001 | Anthropogenic | anaerobic sludge | **ENVO_00002129 |
| Marine | hydrothermal fluid | **ENVO_01000134 | Terrestrial | mud volcano | **ENVO_00000402 | Anthropogenic | aquaduct | **ENVO_00000072 |
| Marine | hydrothermal vent | **ENVO_00000215 | Terrestrial | muddy water | **ENVO_00005793 | Anthropogenic | aquarium | **ENVO_00002196 |

| | | | | | | | | |
|---|---|---|---|---|---|---|---|---|
| Anthropogenic | inlet | **ENVO_ 00002267 | Terrestrial | mushroom compost | **ENVO_ 00003033 | Anthropogenic | artificial reef | **ENVO_ 00000149 |
| Marine | Intertidal zone | **ENVO_ 00000316 | Terrestrial | national park | **ENVO_ 00000367 | Anthropogenic | asphalt lake | **ENVO_ 00000165 |
| Marine | lagoon | **ENVO_ 00000038 | Terrestrial | nature reserve | **ENVO_ 00000363 | Anthropogenic | bar | **ENVO_ 00000167 |
| Marine | mangrove swamp | **ENVO_ 00000057 | Terrestrial | nest of termite | **ENVO_ 02000006 | Anthropogenic | biofilm | **ENVO_ 00002034 |
| Marine | marine bulk water | **ENVO_ 01000055 | Terrestrial | nesting material | **ENVO_ 02000004 | Anthropogenic | biofilter | **ENVO_ 00002152 |
| Marine | marine habitat | **ENVO_ 00000569 | Terrestrial | nunatak | **ENVO_ 00000181 | Anthropogenic | bioreactor | **ENVO_ 00002123 |
| Marine | marine hydrothermal vent | **ENVO_ 01000122 | Terrestrial | oil | **ENVO_ 00002985 | Anthropogenic | biosolids | **ENVO_ 00002059 |
| Marine | marine reef | **ENVO_ 01000143 | Terrestrial | oil reservoir | **ENVO_ 00002185 | Anthropogenic | borehole | **ENVO_ 00002226 |
| Marine | marine sediment | **ENVO_ 00002113 | Terrestrial | oil seep | **ENVO_ 00002063 | Anthropogenic | brackish water | **ENVO_ 00002019 |
| Marine | marine snow | **ENVO_ 01000158 | Terrestrial | orchard | **ENVO_ 00000115 | Anthropogenic | brewery | **ENVO_ 00003885 |
| Marine | marine terrace | **ENVO_ 00000509 | Terrestrial | ornithogenic soil | **ENVO_ 00005782 | Anthropogenic | brine | **ENVO_ 00003044 |
| Marine | mesopelagic zone | **ENVO_ 00000213 | Terrestrial | paddy field | **ENVO_ 00000297 | Anthropogenic | brine pool | **ENVO_ 00000369 |
| Marine | mudflat | **ENVO_ 00000241 | Terrestrial | paddy field soil | **ENVO_ 00005740 | Anthropogenic | canal | ** ENVO _0000014 0 |
| Marine | ocean | **ENVO_ 00000181 | Terrestrial | pasture | **ENVO_ 00000266 | Anthropogenic | carcass | **ENVO_ 00002033 |
| Marine | ocean basin | **ENVO_ 00002450 | Terrestrial | pasture soil | **ENVO_ 00005773 | Anthropogenic | channel | **ENVO_ 00000395 |
| Marine | ocean time series station | **ENVO_ 00011764 | Terrestrial | peat soil | **ENVO_ 00005774 | Anthropogenic | city | **ENVO_ 00000856 |
| Marine | ocean trench | **ENVO_ 00000275 | Terrestrial | peat swamp | **ENVO_ 00000189 | Anthropogenic | coal mine | **ENVO_ 00002169 |
| Marine | ocean water | **ENVO_ 00002151 | Terrestrial | peatland | **ENVO_ 00000044 | Anthropogenic | compost | **ENVO_ 00002170 |
| Marine | oceanic zone | **ENVO_ 00000207 | Terrestrial | pebble | **ENVO_ 00002139 | Anthropogenic | compost soil | **ENVO_ 00005747 |
| Marine | pelagic zone | **ENVO_ 00000211 | Terrestrial | peninsula | **ENVO_ 00000305 | Anthropogenic | contaminated sediment | **ENVO_ 00002114 |
| Marine | photic zone | **ENVO_ 00000209 | Terrestrial | permafrost | **ENVO_ 00000134 | Anthropogenic | contaminated soil | **ENVO_ 00002116 |
| Marine | reef | **ENVO_ 00000130 | Terrestrial | pig manure | **ENVO_ 00003860 | Anthropogenic | contaminated water | **ENVO_ 00002186 |
| Marine | saline marsh | **ENVO_ 00000054 | Terrestrial | piggery | **ENVO_ 00003042 | Anthropogenic | creosote contaminated soil | **ENVO_ 00002117 |
| Marine | saline pan | **ENVO_ 00000279 | Terrestrial | plain | **ENVO_ 00000086 | Anthropogenic | cultivated habitat | **ENVO_ 00000113 |
| Marine | saline water | **ENVO_ 00002010 | Terrestrial | plantation | **ENVO_ 00000117 | Anthropogenic | cultured habitat | **ENVO _0100031 2 |
| Marine | Sea | **ENVO_ 00000016 | Terrestrial | plateau | **ENVO_ 00000182 | Anthropogenic | ditch | **ENVO_ 00000037 |
| Marine | sea floor | **ENVO_ 00000482 | Terrestrial | poultry litter | **ENVO_ 00002192 | Anthropogenic | farm | **ENVO_ 00000078 |
| Marine | sea grass bed | **ENVO_ 01000059 | Terrestrial | prairie | **ENVO_ 00000260 | Anthropogenic | farm soil | **ENVO_ 00005749 |
| Marine | sea ice | **ENVO_ 00002200 | Terrestrial | red clay | **ENVO_ 02000045 | Anthropogenic | fish farm | **ENVO_ 00000294 |
| Marine | sea shore | **ENVO_ 00000485 | Terrestrial | red soil | **ENVO_ 00005790 | Anthropogenic | fishpond | **ENVO_ 00000056 |
| Marine | sea water | **ENVO_ 00002149 | Terrestrial | regosol | **ENVO_ 00002256 | Anthropogenic | gold mine | **ENVO_ 00002168 |
| Marine | seamount | **ENVO_ 00000264 | Terrestrial | rhizosphere | **ENVO_ 00005801 | Anthropogenic | harbor | **ENVO_ 00000463 |
| Marine | Surface water | **ENVO_ 00002042 | Terrestrial | rice field | **ENVO_ 00000296 | Anthropogenic | hospital | **ENVO_ 00002173 |

| | | | | | | | | |
|---|---|---|---|---|---|---|---|---|
| Marine | tidal mudflat | **ENVO_ 00000241 | Terrestrial | ridge | **ENVO_ 00000283 | Anthropogenic | industrial waste | **ENVO_ 00002267 |
| Marine | tidal pool | **ENVO_ 00000317 | Terrestrial | rift valley | **ENVO_ 00000302 | Anthropogenic | island | **ENVO_ 00000475 |
| Marine | undersea feature | **ENVO_ 00000104 | Terrestrial | river bank | **ENVO_ 00000143 | Anthropogenic | landfill | **ENVO_ 00000533 |
| Marine | upwelling | **ENVO_ 01000005 | Terrestrial | river bed | **ENVO_ 00000384 | Anthropogenic | leachate | **ENVO_ 00002141 |
| Marine | watercourse | ** ENVO _00000029 | Terrestrial | river valley | **ENVO_ 00000171 | Anthropogenic | metal contaminated soil | **ENVO_ 00003081 |
| Terrestrial | acid hot spring | **ENVO_ 00002120 | Terrestrial | saline evaporation pond | **ENVO_ 00000055 | Anthropogenic | mine | **ENVO_ 00000076 |
| Terrestrial | acrisol | **ENVO_ 00002234 | Terrestrial | saline lake | **ENVO_ 00000019 | Anthropogenic | mine drainage | **ENVO_ 00001996 |
| Terrestrial | aerosol | **ENVO_ 00010505 | Terrestrial | saline lake sediment | **ENVO_ 00002209 | Anthropogenic | mine tailing | **ENVO_ 00000003 |
| Terrestrial | agricultural soil | **ENVO_ 00002259 | Terrestrial | saline marsh | **ENVO_ 00000054 | Anthropogenic | mushroom compost | **ENVO_ 00003033 |
| Terrestrial | alkaline salt lake | **ENVO_ 00002121 | Terrestrial | saline pan | **ENVO_ 00000279 | Anthropogenic | nuclear power plant | **ENVO_ 00002271 |
| Terrestrial | alpine soil | **ENVO_ 00005741 | Terrestrial | sand | **ENVO_ 01000017 | Anthropogenic | oil contaminated soil | **ENVO_ 00002875 |
| Terrestrial | animal habitation | **ENVO_ 00005803 | Terrestrial | sandstone | **ENVO_ 00002055 | Anthropogenic | oil field production water | **ENVO_ 00002194 |
| Terrestrial | animal waste | **ENVO_ 00002276 | Terrestrial | sandy sediment | **ENVO_ 01000118 | Anthropogenic | oil seep | **ENVO_ 00002063 |
| Terrestrial | aquifer | **ENVO_ 00012408 | Terrestrial | savanna soil | **ENVO_ 00005746 | Anthropogenic | oil spill | **ENVO_ 00002061 |
| Terrestrial | arable soil | **ENVO_ 00005742 | Terrestrial | scrubland | **ENVO_ 00000300 | Anthropogenic | organic waste | **ENVO_ 00002873 |
| Terrestrial | arenosol | **ENVO_ 00002229 | Terrestrial | sea sand | **ENVO_ 00002118 | Anthropogenic | petroleum | **ENVO_ 00002984 |
| Terrestrial | arid | **ENVO_ 01000230 | Terrestrial | sediment | **ENVO_ 00002007 | Anthropogenic | pinnacle | **ENVO_ 00000481 |
| Terrestrial | bagasse | **ENVO_ 00002872 | Terrestrial | sedimentary rock | **ENVO_ 00002016 | Anthropogenic | pothole | **ENVO_ 00000534 |
| Terrestrial | bar | **ENVO_ 00000167 | Terrestrial | shale | **ENVO_ 00002056 | Anthropogenic | poultry litter | **ENVO_ 00002192 |
| Terrestrial | basalt | **ENVO_ 01000236 | Terrestrial | shore | **ENVO_ 00000304 | Anthropogenic | research station | **ENVO_ 00003919 |
| Terrestrial | beach | **ENVO_ 00000091 | Terrestrial | silage | **ENVO_ 00003030 | Anthropogenic | road | **ENVO_ 00000064 |
| Terrestrial | beach sand | **ENVO_ 00002138 | Terrestrial | slope | **ENVO_ 00002000 | Anthropogenic | roadside soil | **ENVO_ 00005743 |
| Terrestrial | beech forest soil | **ENVO_ 00005770 | Terrestrial | soil | **ENVO_ 00001998 | Anthropogenic | scum | **ENVO_ 00003930 |
| Terrestrial | biofilm | **ENVO_ 00002034 | Terrestrial | solonchak | **ENVO_ 00002252 | Anthropogenic | sewage | **ENVO_ 00002018 |
| Terrestrial | botanical garden | **ENVO_ 00010624 | Terrestrial | sound | **ENVO_ 00000393 | Anthropogenic | sludge | **ENVO_ 00002044 |
| Terrestrial | boulder | **ENVO_ 01000243 | Terrestrial | sphagnum bog | **ENVO_ 00002268 | Anthropogenic | Superfund site | **ENVO_ 00002156 |
| Terrestrial | brackish lake | **ENVO_ 00000540 | Terrestrial | stalactite | **ENVO_ 00000331 | Anthropogenic | tannery | **ENVO_ 00003323 |
| Terrestrial | buffer zone | **ENVO_ 00000135 | Terrestrial | steppe | **ENVO_ 00000262 | Anthropogenic | textile | **ENVO_ 02000001 |
| Terrestrial | bulk soil | **ENVO_ 00005802 | Terrestrial | stream sediment | **ENVO_ 00002127 | Anthropogenic | waste | **ENVO_ 00002264 |
| Terrestrial | canyon | **ENVO _00000169 | Terrestrial | subterrestrial habitat | **ENVO_ 00000572 | Anthropogenic | waste treatment plant | **ENVO_ 00002272 |
| Terrestrial | cave | **ENVO_ 00000067 | Terrestrial | subtropical | **ENVO_ 01000205 | Anthropogenic | waste water | **ENVO_ 00002001 |
| Terrestrial | cave system | **ENVO_ 00000013 | Terrestrial | surface soil | **ENVO_ 02000059 | Anthropogenic | wastewater treatment plant | **ENVO_ 00002043 |
| Terrestrial | chaparral | **ENVO_ 00000301 | Terrestrial | swamp | **ENVO_ 00000233 | Anthropogenic | well | **ENVO_ 00000026 |

| | | | | | | | | |
|---|---|---|---|---|---|---|---|---|
| Terrestrial | clay | **ENVO_00002982 | Terrestrial | temperate | **ENVO_01000206 | Unclassified | ENVO 00000428 | **ENVO_00000428 |
| Terrestrial | clay sediment | **ENVO_01000120 | Terrestrial | terrace | **ENVO_00000508 | Unclassified | ENVO 00000446 | **ENVO_00000446 |
| Terrestrial | clay soil | **ENVO_00002262 | Terrestrial | terrestrial habitat | **ENVO_00002009 | Unclassified | ENVO 00000447 | **ENVO_00000447 |
| Terrestrial | cliff | **ENVO_00000087 | Terrestrial | tidal mudflat | **ENVO_00000241 | Unclassified | ENVO 00000479 | **ENVO_00000479 |
| Terrestrial | coast | **ENVO_00000303 | Terrestrial | travertine | **ENVO_00003982 | Unclassified | ENVO 00000873 | **ENVO_00000873 |
| Terrestrial | crater | **ENVO_00000514 | Terrestrial | tropical | **ENVO_01000204 | Unclassified | ENVO 00002030 | **ENVO_00002030 |
| Terrestrial | desert | **ENVO_00000097 | Terrestrial | trough | **ENVO_00000499 | Unclassified | ENVO 01000020 | **ENVO_01000020 |
| Terrestrial | ditch | **ENVO_00000037 | Terrestrial | tundra | **ENVO_00000112 | Unclassified | ENVO 01000047 | **ENVO_01000047 |
| Terrestrial | dry valley | **ENVO_00000128 | Terrestrial | upland soil | **ENVO_00005786 | Unclassified | ENVO 01000048 | **ENVO_01000048 |
| Terrestrial | dune | **ENVO_00000170 | Terrestrial | valley | **ENVO_00000100 | Unclassified | ENVO 01000181 | **ENVO_01000181 |
| Terrestrial | elevation | **ENVO_00000176 | Terrestrial | vineyard | **ENVO_00000116 | Unclassified | ENVO 01000193 | **ENVO_01000193 |
| Terrestrial | endolithic habitat | **ENVO_00000886 | Terrestrial | volcanic field | **ENVO_00000354 | Unclassified | ENVO 01000193 | **ENVO_01000193 |
| Terrestrial | farm soil | **ENVO_00005749 | Terrestrial | volcanic soil | **ENVO_00005785 | Unclassified | ENVO 01000196 | **ENVO_01000196 |
| Terrestrial | fen | **ENVO_00000232 | Terrestrial | volcano | **ENVO_00000247 | Unclassified | ENVO 01000199 | **ENVO_01000199 |
| Terrestrial | field soil | **ENVO_00005755 | Terrestrial | woodland | **ENVO_00000109 | Unclassified | ENVO:00000428 | **ENVO_00000428 |
| Terrestrial | flood plain | **ENVO_00000255 | Terrestrial | zoological garden | **ENVO_00010625 | Unclassified | ENVO:00000446 | **ENVO_00000446 |
| Terrestrial | forest | **ENVO_00000111 | Fresh water | alpine glacier | **ENVO_00000085 | Unclassified | ENVO:00000447 | **ENVO_00000447 |
| Terrestrial | forest soil | **ENVO_00002261 | Fresh water | aquatic habitat | **ENVO_00000144 | Unclassified | ENVO:00000873 | **ENVO_00000873 |
| Terrestrial | fumarole | **ENVO_00000243 | Fresh water | drainage basin | **ENVO_00000291 | Unclassified | ENVO:00002030 | **ENVO_00002030 |
| Terrestrial | garden | **ENVO_00000216 | Fresh water | epilimnion | **ENVO_00002131 | Unclassified | ENVO:01000020 | **ENVO_01000020 |
| Terrestrial | garden soil | **ENVO_00002263 | Fresh water | fresh water | **ENVO_00002011 | Unclassified | ENVO:01000047 | **ENVO_01000047 |
| Terrestrial | geothermal field | **ENVO_00000373 | Fresh water | fresh water | **ENVO_00002011 | Unclassified | ENVO:01000176 | **ENVO_01000176 |
| Terrestrial | glacier | **ENVO_00000133 | Fresh water | freshwater habitat | **ENVO_00002037 | Unclassified | ENVO:01000177 | **ENVO_01000177 |
| Terrestrial | grassland | **ENVO_00005750 | Fresh water | freshwater lake | **ENVO_00000021 | Unclassified | ENVO:01000181 | **ENVO_01000181 |
| Terrestrial | grassland soil | **ENVO_00005750 | Fresh water | freshwater wetland | **ENVO_00000243 | Unclassified | ENVO:01000193 | **ENVO_01000193 |
| Terrestrial | greenhouse soil | **ENVO_00005780 | Fresh water | headwater | **ENVO_00000153 | Unclassified | ENVO:01000196 | **ENVO_01000196 |
| Terrestrial | ground water | **ENVO_00002041 | Fresh water | hypolimnion | **ENVO_00002130 | Unclassified | habitat | **ENVO_00002036 |
| Terrestrial | hot spring | **ENVO_00000051 | Fresh water | iceberg | **ENVO_00000298 | | | |

** http://purl.obolibrary.org/obo/

**Table S8: Pearson correlation of bacterial taxa and wind speed.** Bacterial taxa showing significant linear correlation with wind speed are classified as "oceanic-originated", and "non-oceanic originated" in both transects, with the Pearson correlation coefficient and the statistical significance.

| **Pacific Taxa originate from the ocean** | **Pearson coefficient** | ***p*-Value** |
|---|---|---|
| Planctomycetes_Phycisphaerae_Phycisphaerales_Phycisphaeraceae_Urania-1B-19_marine_sediment_group_NA | 0.597 | 0.001 |
| Proteobacteria_Alphaproteobacteria_Puniceispirillales_SAR116_clade_NA_NA | 0.580 | 0.001 |
| Verrucomicrobia_Verrucomicrobiae_Opitutales_Puniceicoccaceae_Coraliomargarita_NA | 0.569 | 0.002 |
| Cyanobacteria_Oxyphotobacteria_Synechococcales_Cyanobiaceae_Prochlorococcus_MIT9313_marinus | 0.539 | 0.003 |
| Bacteroidetes_Bacteroidia_Flavobacteriales_Flavobacteriaceae_NS4_marine_group_NA | 0.524 | 0.004 |
| Cyanobacteria_Oxyphotobacteria_Synechococcales_Cyanobiaceae_Synechococcus_CC9902_NA | 0.522 | 0.004 |
| Planctomycetes_Phycisphaerae_Phycisphaerales_Phycisphaeraceae_CL500-3_NA | 0.513 | 0.005 |
| Proteobacteria_Gammaproteobacteria_Betaproteobacteriales_Burkholderiaceae_NA_NA | 0.478 | 0.010 |
| Cyanobacteria_Oxyphotobacteria_Synechococcales_Cyanobiaceae_Prochlorococcus_MIT9313_NA | 0.456 | 0.015 |
| Proteobacteria_Gammaproteobacteria_Vibrionales_Vibrionaceae_Vibrio_NA | 0.446 | 0.017 |
| Proteobacteria_Alphaproteobacteria_Parvibaculales_PS1_clade_NA_NA | 0.444 | 0.018 |
| Proteobacteria_Deltaproteobacteria_Bdellovibrionales_Bdellovibrionaceae_Bdellovibrio_NA | 0.422 | 0.025 |
| Bacteroidetes_Bacteroidia_Flavobacteriales_NS9_marine_group_NA_NA | 0.417 | 0.027 |
| Proteobacteria_Alphaproteobacteria_SAR11_clade_Clade_I_Clade_Ia_NA | 0.412 | 0.029 |
| Proteobacteria_Alphaproteobacteria_SAR11_clade_Clade_IV_NA_NA | 0.411 | 0.030 |
| Kiritimatiellaeota_Kiritimatiellae_Kiritimatiellales_Kiritimatiellaceae_R76-B128_NA | 0.401 | 0.034 |
| Proteobacteria_Gammaproteobacteria_Alteromonadales_Idiomarinaceae_Idiomarina_NA | 0.397 | 0.036 |
| **Atlantic Taxa originate from the ocean** | | |
| Proteobacteria_Alphaproteobacteria_Rhodospirillales_AEGEAN-169_marine_group_NA_NA | 0.394 | 0.028 |
| **Pacific Taxa not originate from the ocean** | | |
| Bacteroidetes_Bacteroidia_Bacteroidales_Prevotellaceae_Alloprevotella_NA | 0.474 | 0.011 |
| Actinobacteria_Actinobacteria_Frankiales_Geodermatophilaceae_Modestobacter_NA | 0.402 | 0.034 |
| Actinobacteria_Actinobacteria_Micrococcales_Micrococcaceae_Micrococcus_NA | 0.399 | 0.035 |
| Proteobacteria_Alphaproteobacteria_Rhodobacterales_Rhodobacteraceae_Paracoccus_caeni | -0.384 | 0.043 |
| Proteobacteria_Deltaproteobacteria_Myxococcales_Polyangiaceae_Pajaroellobacter_NA | -0.384 | 0.043 |
| Actinobacteria_Actinobacteria_Micrococcales_Micrococcaceae_Arthrobacter_NA | -0.456 | 0.015 |
| **Atlantic Taxa not originate from the ocean** | | |
| Actinobacteria_Acidimicrobiia_Microtrichales_Ilumatobacteraceae_Ilumatobacter_NA | 0.575 | 0.001 |
| Firmicutes_Bacilli_Lactobacillales_Carnobacteriaceae_Alloiococcus_NA | 0.463 | 0.009 |
| Actinobacteria_Actinobacteria_Micrococcales_Intrasporangiaceae_Janibacter_NA | 0.443 | 0.013 |
| Firmicutes_Bacilli_Lactobacillales_Carnobacteriaceae_Alloiococcus_otitis | 0.410 | 0.022 |
| Proteobacteria_Gammaproteobacteria_Pseudomonadales_Moraxellaceae_Alkanindiges_NA | 0.398 | 0.027 |
| Actinobacteria_Actinobacteria_Pseudonocardiales_Pseudonocardiaceae_Pseudonocardia_NA | 0.375 | 0.038 |
| Firmicutes_Bacilli_Bacillales_Alicyclobacillaceae_Tumebacillus_NA | 0.371 | 0.040 |
| Actinobacteria_Actinobacteria_Micrococcales_Microbacteriaceae_Agrococcus_NA | 0.360 | 0.047 |
| Actinobacteria_Actinobacteria_Micrococcales_Dermabacteraceae_Dermabacter_NA | -0.362 | 0.045 |
| Actinobacteria_Actinobacteria_Micrococcales_Micrococcaceae_Kocuria_NA | -0.382 | 0.034 |
| Firmicutes_Bacilli_Lactobacillales_Aerococcaceae_Abiotrophia_defectiva | -0.447 | 0.012 |

### References


1. D. Hospodsky, N. Yamamoto, J. Peccia, Accuracy, precision, and method detection limits of quantitative PCR for airborne bacteria and fungi. *Appl Environ Microbiol* **76**, 7004-7012 (2010).
2. N. Lang-Yona *et al.*, Annual distribution of allergenic fungal spores in atmospheric particulate matter in the Eastern Mediterranean; a comparative study between ergosterol and quantitative PCR analysis. *Atmos. Chem. Phys.* **12**, 2681-2690 (2012).
3. S. Sharoni *et al.*, Infection of phytoplankton by aerosolized marine viruses. *Proc Natl Acad Sci U S A* **112**, 6643-6647 (2015).
4. N. M. Davis, D. M. Proctor, S. P. Holmes, D. A. Relman, B. J. Callahan, Simple statistical identification and removal of contaminant sequences in marker-gene and metagenomics data. *Microbiome* **6**, 226 (2018).
5. R. Eisenhofer *et al.*, Contamination in Low Microbial Biomass Microbiome Studies: Issues and Recommendations. *Trends Microbiol* **27**, 105-117 (2019).
6. S. J. Salter *et al.*, Reagent and laboratory contamination can critically impact sequence-based microbiome analyses. *BMC Biology* **12**, 87 (2014).
7. E. R. Lewis, S. E. Schwartz, in *Sea Salt Aerosol Production: Mechanisms, Methods, Measurements and Models*, E. R. Lewis, S. E. Schwartz, Eds. (American Geophysical Union, 2013), pp. 9-99.
8. S.-L. von der Weiden, F. Drewnick, S. Borrmann, Particle Loss Calculator - a new software tool for the assessment of the performance of aerosol inlet systems. *Atmos Meas Tech Discuss* **2**, 1099 (2009).
9. J. M. Flores *et al.*, Tara Pacific expedition's atmospheric measurements. Marine aerosols across the Atlantic and Pacific Oceans Overview and Preliminary results. *Bull Am Meteorol Soc* **101**, 536-554 (2020).
10. M. E. Allentoft *et al.*, The half-life of DNA in bone: measuring decay kinetics in 158 dated fossils. *Proc R Soc Lond B Biol Sci* **279**, 4724-4733 (2012).
11. J. Peccia, M. Hernandez, Incorporating polymerase chain reaction-based identification, population characterization, and quantification of microorganisms into aerosol science: A review. *Atmos Environ (1994)* **40**, 3941-3961 (2006).
12. A. Z. Ijaz, T. C. Jeffries, U. Z. Ijaz, K. Hamonts, B. K. Singh, Extending SEQenv: a taxa-centric approach to environmental annotations of 16S rDNA sequences. *PeerJ* **5**, e3827-e3827 (2017).
13. L. Sinclair *et al.*, Seqenv: linking sequences to environments through text mining. *PeerJ* **4**, e2690 (2016).
14. P. L. Buttigieg *et al.*, The environment ontology: contextualising biological and biomedical entities. *J Biomed Semant* **4**, 43 (2013).
